# The neuroendocrine transition in prostate cancer is dynamic and dependent on ASCL1

**DOI:** 10.1101/2024.04.09.588557

**Authors:** Rodrigo Romero, Tinyi Chu, Tania J. González-Robles, Perianne Smith, Yubin Xie, Harmanpreet Kaur, Sara Yoder, Huiyong Zhao, Chenyi Mao, Wenfei Kang, Maria V. Pulina, Kayla E. Lawrence, Anuradha Gopalan, Samir Zaidi, Kwangmin Yoo, Jungmin Choi, Ning Fan, Olivia Gerstner, Wouter R. Karthaus, Elisa DeStanchina, Kelly V. Ruggles, Peter M.K. Westcott, Ronan Chaligné, Dana Pe’er, Charles L. Sawyers

## Abstract

Lineage plasticity is a recognized hallmark of cancer progression that can shape therapy outcomes. The underlying cellular and molecular mechanisms mediating lineage plasticity remain poorly understood. Here, we describe a versatile *in vivo* platform to identify and interrogate the molecular determinants of neuroendocrine lineage transformation at different stages of prostate cancer progression. Adenocarcinomas reliably develop following orthotopic transplantation of primary mouse prostate organoids acutely engineered with human-relevant driver alterations (e.g., *Rb1^-/-^*; *Trp53^-/-^*; *cMyc^+^* or *Pten^-/-^*; *Trp53^-/-^*; *cMyc^+^*), but only those with *Rb1* deletion progress to ASCL1+ neuroendocrine prostate cancer (NEPC), a highly aggressive, androgen receptor signaling inhibitor (ARSI)-resistant tumor. Importantly, we show this lineage transition requires a native *in vivo* microenvironment not replicated by conventional organoid culture. By integrating multiplexed immunofluorescence, spatial transcriptomics and PrismSpot to identify cell type-specific spatial gene modules, we reveal that ASCL1+ cells arise from KRT8+ luminal epithelial cells that progressively acquire transcriptional heterogeneity, producing large ASCL1^+^;KRT8^-^ NEPC clusters. *Ascl1* loss in established NEPC results in transient tumor regression followed by recurrence; however, *Ascl1* deletion prior to transplantation completely abrogates lineage plasticity, yielding adenocarcinomas with elevated AR expression and marked sensitivity to castration. The dynamic feature of this model reveals the importance of timing of therapies focused on lineage plasticity and offers a platform for identification of additional lineage plasticity drivers.

## INTRODUCTION

Prostate cancer is the leading cause of cancer death globally in men^1^. Survival has improved through development of next generation ARSIs; however, patients eventually progress to castration-resistant prostate cancer^2^. Although men receiving ARSIs are living longer, an increasing fraction display features of lineage plasticity at relapse, characterized by reduced or absent expression of luminal lineage markers such as AR and the downstream target gene prostate specific antigen^3,4^. In its most extreme form, lineage plasticity manifests as a transition to neuroendocrine (NE) histology called NEPC, with expression of synaptophysin (SYP) and chromogranin^4^. NEPC histology is more commonly seen in patients with metastasis to soft tissue (e.g., liver) rather than bone, raising a potential role of the tumor microenvironment (TME) in this transition^5,6^. Similar lineage transitions are observed in other tumor types treated with targeted therapies, such as *EGFR*-, *ALK*-, and *KRAS^G12C^*-mutant lung adenocarcinoma, underscoring the broad relevance of lineage plasticity in tumor progression and therapy resistance^7–11^.

The molecular details underlying these lineage transitions are poorly understood, largely owing to a shortage of tractable model systems that accurately and dynamically replicate plasticity-associated transitions observed in patients. Autochthonous models of prostate cancer have contributed substantially to our understanding of prostate tumor progression, but few capture the transition at all stages or are amenable to intervention in a timely and cost-effective manner^12–17^. Conversely, studies using prostate tumor cell line transplant models can be completed more quickly, but the number of models is limited, and they fail to replicate all stages of the lineage transition that occurs in patients. To gain a better understanding of NEPC and to develop intervention strategies that curtail lineage plasticity, model systems that accurately reproduce the molecular and morphologic features of these lineage transitions over time are needed.

Organoid technology has greatly expanded our ability to model epithelial biology, including prostate cancer initiation and progression^18,19^. Previously, we described a strategy to assess putative genetic drivers of prostate adenocarcinoma (PRAD), as well as tumor cells of origin using mouse prostate organoids coupled with orthotopic transplantation^20^ (OT). Here, we optimize this approach into a robust platform that enables rapid, side-by-side assessment of cancer initiation and progression phenotypes using multiple combinations of human-relevant cancer drivers *in vivo*. Using multiplexed spatial techniques, we detect isolated NE cells emerging from luminal epithelial cells, which subsequently evolve to fully penetrant NEPC, together with temporal changes within the TME, and perform functional perturbations that dramatically impact the lineage plasticity program.

## RESULTS

### Rapid tumor phenotyping across an allelic series of prostate cancer drivers

We sought to develop a platform to interrogate prostate cancer drivers rapidly and comprehensively at larger scale compared to traditional genetically-engineered mouse models (GEMMs), focusing particularly on the unmet need to dynamically model the PRAD-to-NEPC transition observed in patients. Using multiplexed editing approaches^20^ and lentiviral oncogene delivery, we established organoids with six relevant combinations of cancer drivers selected based on their enrichment and co-occurrence in human prostate cancer (**Fig. 1a, Extended Data** Fig. 1a-c**, and Supplementary Table 1**; hereafter: *<u>Pt</u>en*^-/-^; *Tr<u>p</u>53*^-/-^ = PtP, *<u>R</u>b1*^-/-^; *Tr<u>p</u>53*^-/-^ = RP, *<u>Pt</u>en*^-/-^; *<u>R</u>b1*^-/-^ = PtR, *<u>Pt</u>en*^-/-^; *Tr<u>p</u>53*^-/-^; *c<u>M</u>yc*^+^ = PtPM, *<u>R</u>b1*^-/-^; *Tr<u>p</u>53*^-/-^; *c<u>M</u>yc*^+^ = RPM, *<u>Pt</u>en*^-/-^; *<u>R</u>b1*^-/-^; *c<u>M</u>yc*^+^ = PtRM). In line with previous work, histological assessment of edited mouse organoids grown in 3D culture conditions revealed a mixture of KRT5+ basal and KRT8+ luminal cells, with both populations staining for nuclear AR (ref. 18; **Extended Data** Fig. 1d-e). All cultured organoids lacked expression of the NE transcription factors achaete-scute family bHLH transcription factor 1 (ASCL1) and Neuronal Differentiation 1 (NEUROD1), critical regulators of the neuronal and NE lineages in mammalian development, despite prolonged *in vitro* culture (**Extended Data** Fig. 1d-e).

**Figure 1:**
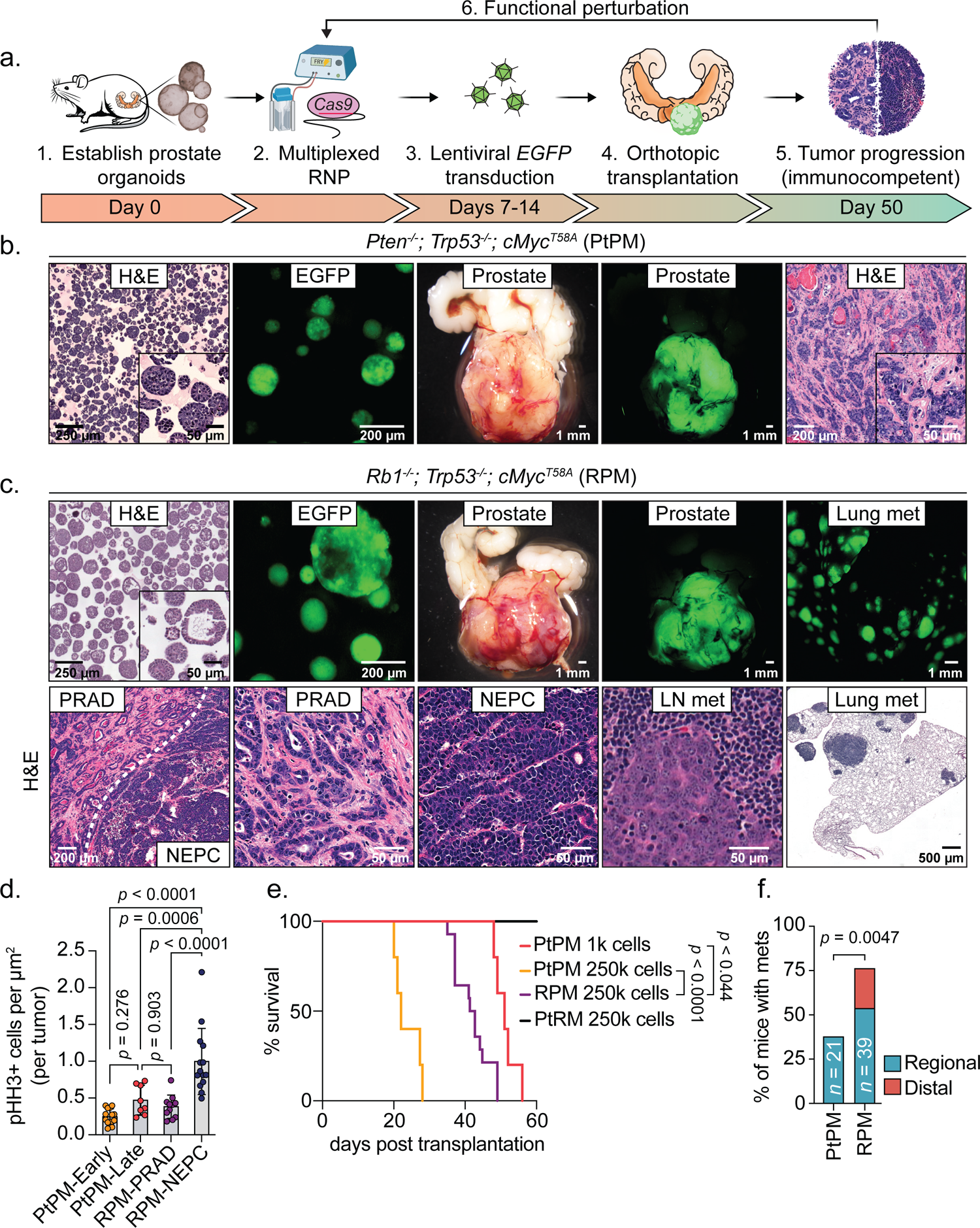
Rapid establishment of genetically-defined prostate cancer with prostate organoids transplanted into immunocompetent syngeneic hosts. **a.** Schematic of timeline required to establish, propagate, edit, and select for organoids harboring mutations in cancer-associated genes prior to transplantation into immunocompetent hosts for tumor establishment. **b.** Representative microscopy of *Pten^-/-^*; *Trp53^-/-^*; *cMyc^T58A^* (PtPM) organoids, and stereoscopic and fluorescence images of orthotopic (OT) prostate tumors with prostate adenocarcinoma (PRAD) histology. Tumor images are representative of *n*=22 independent mice. H&E, hematoxylin and eosin. **c.** Top: Representative microscopy of *Rb1^-/-^*; *Trp53^-/-^*; *cMyc^T58A^*(RPM) organoids, and stereoscopic and fluorescence images of OT prostate tumors and lung metastases. Bottom: Representative histological assessment of RPM-PRAD and RPM-neuroendocrine prostate cancer (NEPC) primary tumor or metastases histology at varying magnifications. Primary and metastatic histology are representative of *n*=25 independent mice. LN, lymph node (iliac). **d.** Phospho-Histone H3 (Ser10; pHH3) positive tumor cells per total tumor area (µm^2^). Each data point represents the average number of pHH3+ cells per individual tumor subset by tumor histology (PRAD vs NEPC) and experimental end point. PtPM-Early (<4 weeks), *n*=14; PtPM-Late (>6 weeks), *n*=8; RPM-PRAD, *n*=11; RPM-NEPC, *n*=14. Statistics derived using one-way ANOVA with Tukey’s multiple comparisons correction. Error bars denote mean and standard deviation. **e.** Survival of mice transplanted with the indicated cell numbers of PtPM, RPM, and *Pten^-/-^*; *Rb1^-/-^*; *cMyc^T58A^* (PtRM) *ex-vivo* edited organoids. PtPM 1k, *n*=5; PtPM 250k, *n*=5; RPM 250k, *n*=14; PtRM 250k, *n*=8. Statistics derived from the Log-rank (Mantel-Cox) test for each pair-wise comparison. **f.** Metastatic disease penetrance of the indicated organoid transplants. Regional metastases include dissemination into the iliac lymph nodes. Distal metastases include dissemination to kidney, pancreas, liver, or lungs. Statistics derived from two-sided Fisher’s exact test. Number of biological replicates indicated within the figure. Scale bars indicated within each figure panel.

Having generated this allelic series, we next evaluated tumorigenicity following OT (**Fig. 1a**). Because expansion of organoids grown in 3D culture is labor intensive (requiring serial propagation of single cell suspensions embedded in matrigel), we compared 3D expansion to short term (five day) monolayer expansion as a simpler alternative (**Extended Data** Fig. 2a-c). Although monolayer expansion was fast and yielded highly penetrant tumor growth for most genotypes (PtP, RP, PtPM, RPM), pathologic evaluation revealed a high frequency of sarcomatoid-like histology that is not seen in typical human prostate cancers^21^ (**Extended Data** Fig. 2d-g). In contrast, tumors arising from organoids expanded exclusively in 3D culture consistently and reliably established histologic phenotypes and lineage marker expression that closely mirror the human disease, particularly for the PtPM and RPM genotypes as detailed below (**Extended Data** Fig. 2d-h). Phenotypes of each of the six combinations of genetic drivers, expanded using 3D or monolayer culture, are summarized in **Supplementary Table 2**. Due to the sarcomatoid-like histology seen following monolayer culture, all subsequent experiments were performed using 3D expansion only.

### *Rb1* loss is a critical gatekeeper event for NEPC transformation

Based on the rapid, highly penetrant development of PRAD using PtPM and RPM organoids, we comprehensively evaluated disease progression across both models (hereafter called PtPM and RPM mice). In these models, we consistently observed PRAD with moderate to poorly differentiated histology during the first 2-3 weeks post transplantation (**Fig. 1b-c**); however, RPM tumors also contained pockets of small cell-like tumors with “salt-and-pepper” chromatin and a mixture of trabecular or diffuse architecture suggestive of NEPC (**Fig. 1c**). The mitotic index in RPM tumors, particularly in large areas of NEPC that emerged late (8-10 weeks), was greater than PtPM tumors, consistent with the rapid disease progression seen clinically in patients with NEPC transformation (**Fig. 1d**). Despite this difference in proliferation rate, the overall survival of PtPM mice was shorter, likely due to higher tumor engraftment potential of PtPM organoids, since a 250-fold reduction in the number of cells injected results in comparable survival to RPM mice (**Fig. 1e and Extended Data** Fig. 2i).

Consistent with the moderately differentiated luminal histology, early RPM tumors displayed significantly more KRT8+ cells compared to KRT5+ cells, markers of luminal and basal identity respectively (**Supplementary** Fig. 1a). Moreover, ASCL1 expression was observed as early as 4 weeks post-engraftment, with a significant increase in the proportion of ASCL1+ cells by 8-10 weeks (**Supplementary** Fig. 1b). These late stage NEPC regions also expressed canonical NE markers such as FOXA2, DLL-3, SYP, NCAM-1, and rarely NEUROD1^4,5,22^ (**Fig. 2a-b**). In contrast, tumors in PtPM mice rarely contained ASCL1+ cells and never progressed to NEPC (**Fig. 2a and Supplementary** Fig. 1c-d). We therefore conclude that functional *Rb1* loss is a critical gatekeeper event required for NEPC transformation, consistent with preclinical and clinical datasets demonstrating enrichment of *RB1* pathway mutations in small cell lung cancer (SCLC) and NEPC^22,23^.

**Figure 2:**
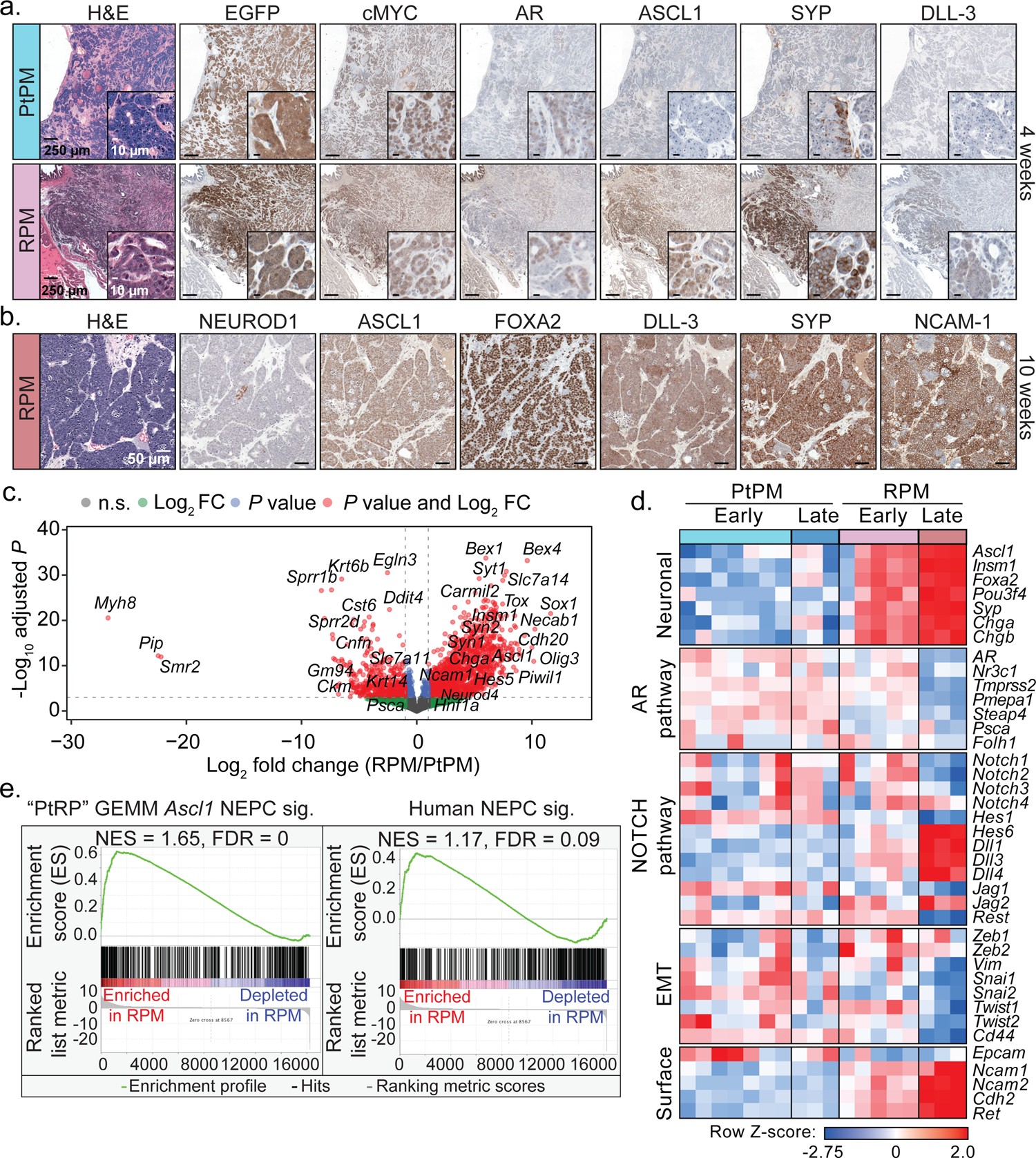
Molecular characterization of PtPM and RPM primary prostate tumor transplants demonstrates emergence of neuroendocrine carcinoma marker expression. **a.** Representative histological analysis of PtPM (top) and RPM (bottom) tumors isolated at 4-weeks post-transplantation. Serial sections depict immunohistochemical staining of the indicated markers. Data are representative of *n*=22 independently transplanted tumors. **b.** Representative histological analysis of RPM tumors isolated at 10-weeks post-transplantation. Serial sections depict immunohistochemical staining of the indicated markers. Data are representative of *n*=25 independently transplanted tumors. **c.** Volcano plot depiction of the log_2_ fold change in RNA expression of primary (OT) RPM tumors relative to primary (OT) PtPM tumors. Genes that meet or surpass the indicated thresholds of significance and fold change in expression are color coded as depicted in the figure legend. Data derived from the comparison of PtPM (*n*=10) and RPM (*n*=8) independent prostate tumors. **d.** Heatmap depicting the *Z*-score normalized differential expression of select genes in PtPM versus RPM tumors. Time points of isolation are color coded in the figure as they are in Fig. 2a. Genes are grouped by the listed class or pathway. Early PtPM ≤4 weeks, early RPM ≤6 weeks. Late PtPM =5 weeks, late RPM =10 weeks. Data related to samples used in Fig. 2f. **e.** Enrichment plots (GSEA) of established expression signatures of (left) genetically engineered mouse model (GEMM) of NEPC harboring conditional deletion of *Pten*, *Rb1*, and *Trp53* (PtRP), and (right) histologically verified human NEPC within RPM primary tumors. FDR and NES indicated in the figure. Analysis derived from the transcriptional profiles of multiple independent RPM tumors (*n*=8) relative to PtPM tumors (*n*=10). Data related to samples used in Fig. 2d. All scale bars noted in each panel and are of equivalent magnification across each marker.

PtPM and RPM mice both developed regional metastases in the draining iliac lymph nodes, but RPM mice also established distant metastases (primarily liver and lung; **Fig. 1b-c, f**). Metastases in RPM mice mostly retained the same NEPC profile seen in primary tumors, except for rare ASCL1-negative patches that were also negative for SYP, NCAM-1, and NEUROD1 but occasionally positive for vimentin (VIM), a marker of mesenchymal-like cells (**Extended Data** Fig. 3a-d). Whether these ASCL1-negative regions reflect ongoing lineage plasticity after metastasis of ASCL1+ cells, or independent metastatic events prior to NEPC transformation, requires further investigation. Interestingly, lung metastases in RPM mice contained a higher proportion of ASCL1+/KRT8+ (double-positive) cells compared to liver metastases (mostly ASCL1 single-positive), and AR expression was absent in tumor cells at both metastatic sites (**Extended Data** Fig. 3e-f).

To further benchmark the PtPM and RPM models relative to autochthonous prostate cancer models and human samples, we performed bulk RNA-sequencing of tumors harvested early (PtPM ≤ 4 weeks, RPM ≤6 weeks) and late (PtPM = 5 weeks, RPM = 10 weeks). Consistent with the immunohistochemical findings, we observed progressive upregulation of genes involved in neuronal differentiation in RPM compared to PtPM tumors, including *Ascl1*, *Foxa2*, *Sox1*, *Chga*, and *Olig3*, several NOTCH pathway ligands^4,24^ (e.g., *Dll1*, *Dll3*, *Hes5*), as well as downregulation of AR and several AR-target genes (e.g., *Tmprss2*, *Pmepa1*, *Folh1*; **Fig. 2c-d, Extended Data** Fig. 4a-b**, and Supplementary Table 3**). Critically, RPM tumors were significantly enriched for transcriptional signatures derived from prostate GEMMs that undergo NEPC transformation and from human NEPC specimens, demonstrating that RPM transplants rapidly establish and recapitulate key molecular features observed in gold-standard preclinical models and clinical samples^13^ (**Fig. 2e, Extended Data** Fig. 4c**, and Supplementary Table 4**). Further highlighting the critical role of the *in vivo* TME in initiating NEPC transformation, *Ascl1* transcript levels were ∼2000-fold higher in RPM tumors compared to long-term cultured RPM organoids. Moreover, the *in vivo* TME is required for maintenance of the NEPC state as *Ascl1* expression progressively declined in RPM tumor-derived organoids (tumoroids; **Extended Data** Fig. 4d).

### Dynamic tumor microenvironment changes during adenocarcinoma to NEPC transition

Because the *in vivo* setting is required to trigger lineage plasticity in the RPM model, we were particularly interested in surveying changes in the TME. Toward that end, we developed a 20-plex immunofluorescence panel to visualize prostate tumor cells (PRAD and NEPC) in the context of adjacent immune populations, vasculature, and stroma (**Fig. 3a-g, and Supplementary Tables 5 and 6**). We focused our analysis on the later stages of tumor progression within the RPM model to identify changes to the TME within large patches of NEPC histology (**Fig. 3b**). We used GFP expression to define tumor cells, together with co-expression of either KRT8 and AR or ASCL1 to distinguish PRAD from NEPC (**Extended Data** Fig. 5a). We selected co-expression of EGFP+/ASCL1- and EGFP+/ASCL1+ as the principal metric to score PRAD and NEPC (**see Methods)**. After mapping these respective regions across multiple tissue sections from RPM tumors containing patches of NEPC differentiation, we then looked for selective changes in cell type composition within the TME.

**Figure 3:**
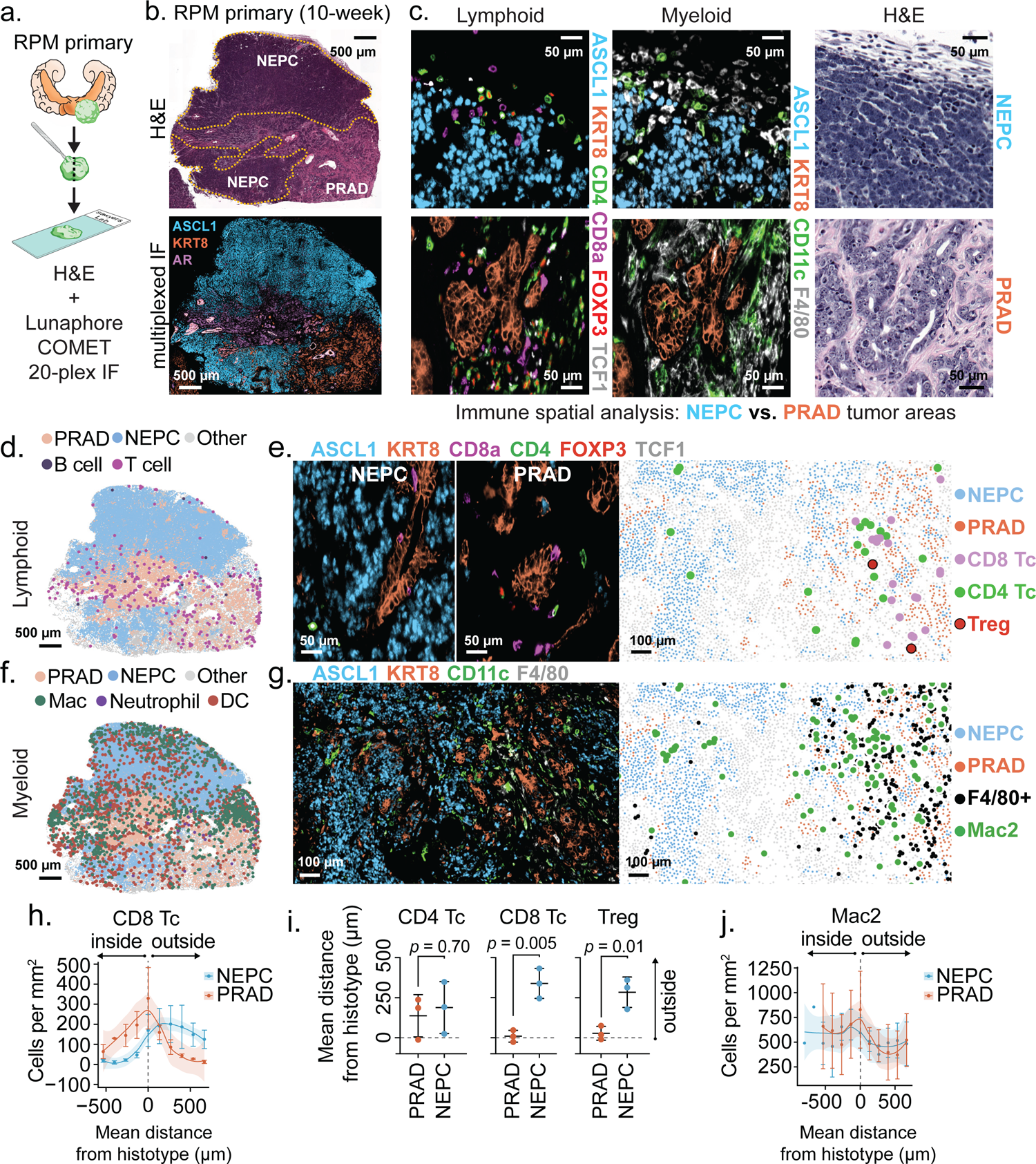
Multiplexed immunofluorescence identifies unique spatial distribution of immune cells within RPM prostate tumors, with local depletion of immune cell types in NEPC areas. **a.** Schematic representation of the methods used to process RPM tumors for 20-plex cyclic immunofluorescence. **b.** (Top) Representative H&E and (bottom) serial section depicting a 3-marker pseudo-colored 10-week RPM tumor. Histological regions (PRAD vs. NEPC) are denoted in the H&E and demarcated by dotted yellow line. **c.** Representative enhanced magnification of lymphoid (left) and myeloid cell markers (middle), and serially sectioned H&E (right). **d.** Representative segmented field of view (FoV) for the indicated general lymphoid cell types in a 10-week RPM tumor. **e.** Representative immunofluorescence of the indicated pseudo-colored lymphocyte markers within NEPC (left) or PRAD (middle). (Right) Data presented as a segmented FoV indicating the localization of each lymphoid and tumor cell type in space. **f.** Representative segmented field of view (FoV) for the indicated general myeloid cell types in a 10-week RPM tumor. **g.** Representative immunofluorescence of the indicated pseudo-colored myeloid and tumor histotype markers. (Right) Segmented FoV indicating the localization of each myeloid and tumor cell type in space. **h.** Frequency distribution of CD8+ T cells within binned distance outside or inside the defined interface region (NEPC or PRAD). Scale bar represents mean and standard error of the mean of the cell counts per bin. **i.** Mean distance of the indicated cell types to the nearest histotype boundary. Error bars denote mean and standard deviation. **j.** Frequency distribution of Mac2 cells (CD11b^lo^; CD11c+; F4/80+) within each binned distance outside or inside of the defined interface region (NEPC or PRAD). Scale bar represents mean and standard error of the mean of the cell counts per bin. Data calculated as in h. Shaded regions in panels h and j approximated through Loess method. Dotted line in h-j represents the boundary of the tumor histotype or tumor edge. All scale bars denoted within each panel. Data derived from *n*=3 independent tumor samples. Infiltration analyses representative of *n*>3 distinct NEPC and PRAD boundaries.

Focusing initially on stroma, we noted that mesenchymal cells were abundant in regions of PRAD but depleted in regions of NEPC. We observed a similar trend for LYVE1+ lymphatics although this did not reach statistical significance. However, there were no obvious differences in CD31+ endothelial populations which localized primarily to the boundaries of NEPC and PRAD (**Supplementary** Fig. 2a-c).

We next turned our attention to immune cells and noted striking depletion of CD8+ and FOXP3+;CD4+ regulatory T cells (Treg) as well as F4/80+ macrophages across all NEPC regions, consistent with reports showing similar absence of immune cells within human NE cancers^23,25,26^. Conversely, FOXP3-;CD4+ T cells were equally distributed within PRAD and NEPC, with a high fraction located at PRAD boundaries, suggestive of differential recruitment and retention of T cell subsets between histologies (**Fig. 3d-e, h-i and Extended Data** Fig. 5b-d). Of the CD8+ T cells within PRAD regions, the vast majority (∼96%) were TCF1-negative, consistent with prior work demonstrating downregulation of TCF1 and upregulation of an effector program in tumor infiltrating compared to draining lymph node resident CD8 T cells^27^ (**Extended Data** Fig. 5e-f).

We identified five distinct myeloid populations which we labeled Mac1 (CD11b+;F4/80-), Mac2 (CD11b^lo^;CD11c+;F4/80+), Mac3 (CD11b+;F4/80+), neutrophil (CD11b+;Ly6G+;S100A9+) and DC (CD11c+;F4/80-; **Extended Data** Fig. 6a **and Supplementary Table 6**). Neutrophil infiltration was low and confined to the outer boundary of PRAD regions (**Fig. 3f and Extended Data** Fig. 6a-b). Mac1 and Mac3 populations were largely absent from the NEPC TME; however, Mac2, which harbors similar marker expression as alveolar and wound-healing macrophages was present within NEPC^28^ (**Fig. 3f-g,j and Extended Data** Fig. 6c-e). We were surprised to also see substantial numbers of CD11c+;F4/80-cells within NEPC regions of primary tumors, raising the possibility of dendritic cell infiltration (**Fig. 3f and Extended Data** Fig. 6a-b).

To determine if the differences in PRAD versus NEPC immune infiltrates in RPM mice are seen in human prostate cancer, we examined a recently published human single cell RNA-sequencing dataset that includes PRAD and NEPC samples^13^. Both histologies had evidence of myeloid infiltration, but NEPC harbored fewer tumor associated macrophages (hereafter abbreviated TAM) relative to PRAD (**Extended Data** Fig. 6f**; see Methods**). However, CD11c (*ITGAX*) expression was evident across TAM populations within both PRAD and NEPC, and highest in *IL1B*+ TAMs (**Extended Data** Fig. 6f-h). We also observed decreased immune infiltration in the NEPC regions of a human prostatectomy specimen from a patient with mixed PRAD/NEPC histology but confirmed the presence of CD11c+;CD68+ macrophage populations within ASCL1+ tumor regions (**Extended Data** Fig. 6i-j). Whether these CD11c+ myeloid populations correspond to professional antigen presenting cells remains uncertain and will require further phenotypic (e.g., MHC-II, CD103, BATF3 expression) and functional characterization. Nonetheless, the evidence of early CD8+ T cell infiltration in PRAD and persistence of potential dendritic cells in late stage NEPC in this model suggest that deeper analysis may be informative in addressing the disappointing clinical results to date using conventional immune checkpoint blockade therapy in prostate cancer.

We next profiled the TME of RPM metastases, a clinically relevant site of NEPC histology (**Fig. 4a-c**). Turning first to RPM lymph node metastases, there was a striking absence of CD45+ cells within ASCL1+ tumor nests, thus highlighting the capacity of NE tumors to promote immune exclusion within lymphocyte-dense microenvironments (**Fig. 4a,d**). Within distant metastases (liver and lung), we also observed exclusion of Treg, CD4+ and CD8+ T cell subsets but retention of IBA-1+ macrophages that co-stain with markers consistent with the Mac1, Mac2, or Mac3 identities seen in the primary tumors, with findings confirmed by neighborhood composition analysis (**Fig. 4b-c, e-i, see Methods**). Taken together, spatial profiling of primary tumors and metastases demonstrates exclusion of nearly all T cell populations within NEPC regions but not PRAD. However, subsets of myeloid cells such as Mac2 and those with DC-like cell surface marker expression (CD11c+ F4/80-) are retained in NEPC. Critically, our syngeneic models are readily suited for studies using model antigens to evaluate strategies to overcome the immunosuppressive prostate tumor microenvironment.

**Figure 4:**
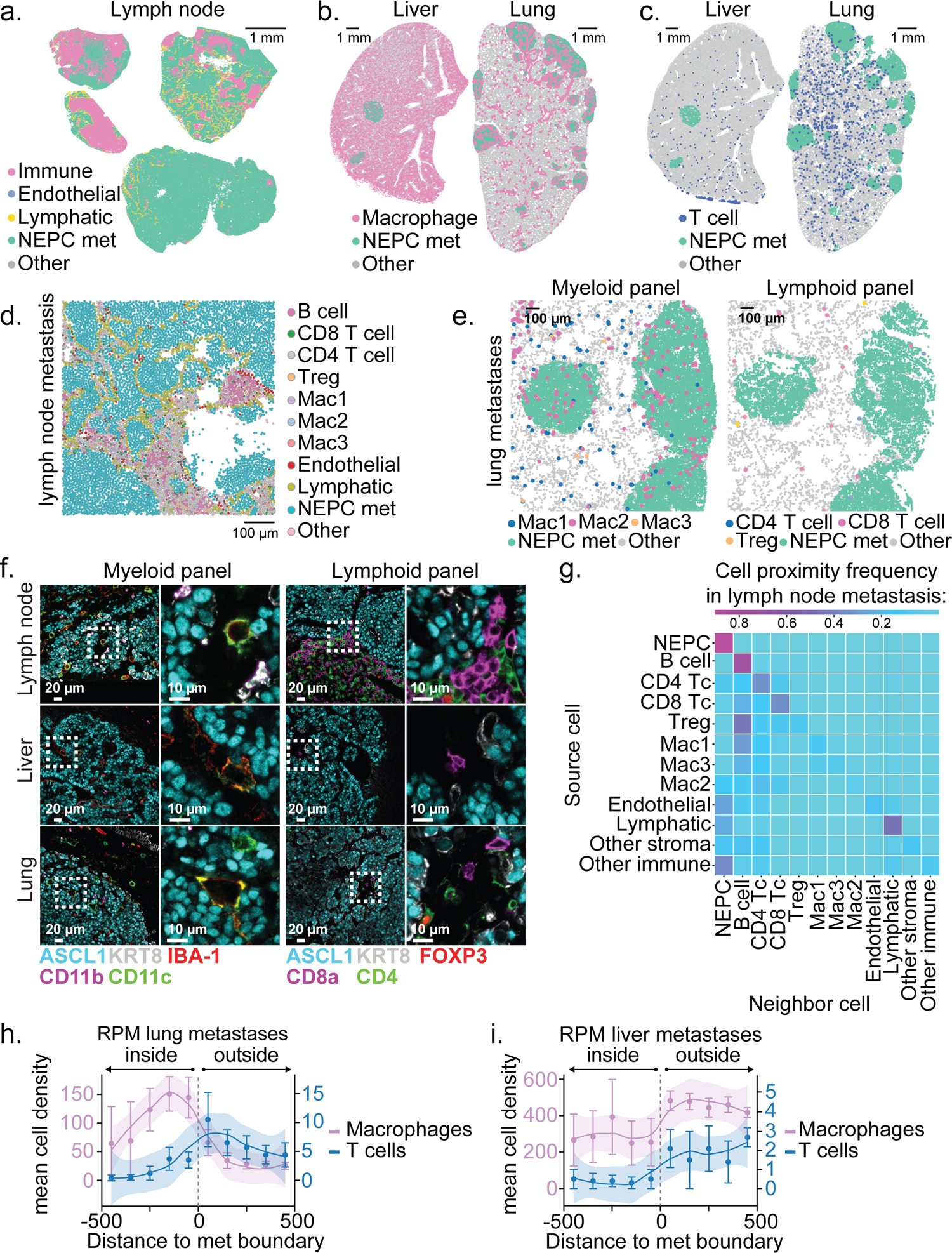
NEPC metastatic lesions are T cell excluded but retain macrophage infiltrates. **a.** Representative segmented field of view (FoV) for the indicated cell types within 4 independent draining lymph node metastases derived from *n*=2 mice transplanted OT with RPM organoids. **b.** Representative segmented FoV of macrophages (IBA-1+) within liver or lung sections obtained from mice transplanted OT with RPM organoids. Note, liver-resident macrophages (Kupffer cells) are IBA-1+. **c.** Representative segmented FoV of T cells (CD4+ or CD8+) within liver or lung sections obtained from mice transplanted OT with RPM organoids. **d.** Representative zoomed in segmented FoV for all cell types listed within a draining lymph node metastasis. Scale denotes relative cell size. **e.** Representative zoomed in segmented FoV across serial lung sections obtained from mice transplanted OT with RPM organoids, identifying NEPC metastatic nodules infiltrated with (left) macrophage subsets or (right) T cell subsets. **f.** Representative multiplexed immunofluorescence of the indicated cell type markers across distinct metastatic sites obtained from mice OT transplanted with RPM organoids. **g.** Neighborhood composition heatmap of cell types found within RPM draining lymph node metastases demonstrating the proximity of the source cell relative to a neighboring cell (20-pixel distance). Data are derived from *n*=4 independent metastatic lymph node samples isolated from *n*=2 mice. **h.** Frequency distribution for Macrophages (IBA1+) or T cells (CD4+ or CD8+) within each binned distance outside or inside of RPM lung metastatic samples. **i.** Frequency distribution for Macrophages (IBA1+) or T cells (CD4+ or CD8+) within each binned distance outside or inside of RPM liver metastatic samples quantified as in Fig. 4h. Shaded region in h-i approximated through Loess method. Scale bar in h-i represents mean and standard error of the mean of the cell counts per bin. Dotted line in h-i represents the boundary of a tumor histotype or tumor edge. All metastatic tumors per section within an individual mouse were combined for infiltration analysis and subsequently averaged between replicates (*n*=3 independent mice).

### Origin and progression of neuroendocrine cells within prostate adenocarcinoma

In addition to tracking changes in the TME, the dynamic nature of the RPM model allows a careful examination of the earliest stages of NEPC transformation. ASCL1, a marker of emerging NE cells, was first detected at 4-6 weeks with the appearance of EGFP+;KRT8+;ASCL1+ tumor cell clusters (**Fig. 5a, Extended Data** Fig. 5a**, and Supplementary** Fig. 1a-b). By 10 weeks, larger homogeneous clusters of ASCL1+;KRT8-tumor cells with small cell NEPC histology were easily visible. The observation that the earliest detectable ASCL1+ cells also co-express KRT8 suggests that NE cells may arise from KRT8+ luminal cells. Indeed, KRT8+;ASCL1+ cells were 4- to 5-fold more abundant than KRT5+;ASCL1+ cells at intermediate timepoints (6-weeks, *p* = 0.025 two-tailed *t*-test, **Fig. 5b**). At later time points (8-10 weeks) primary and metastatic tumor cells were mostly AR-negative and ASCL1-positive with heterogenous expression of KRT8 and E-cadherin (**Fig. 3b, Fig. 5b, and Extended Data** Fig. 3e-f **and 5a**).

**Figure 5:**
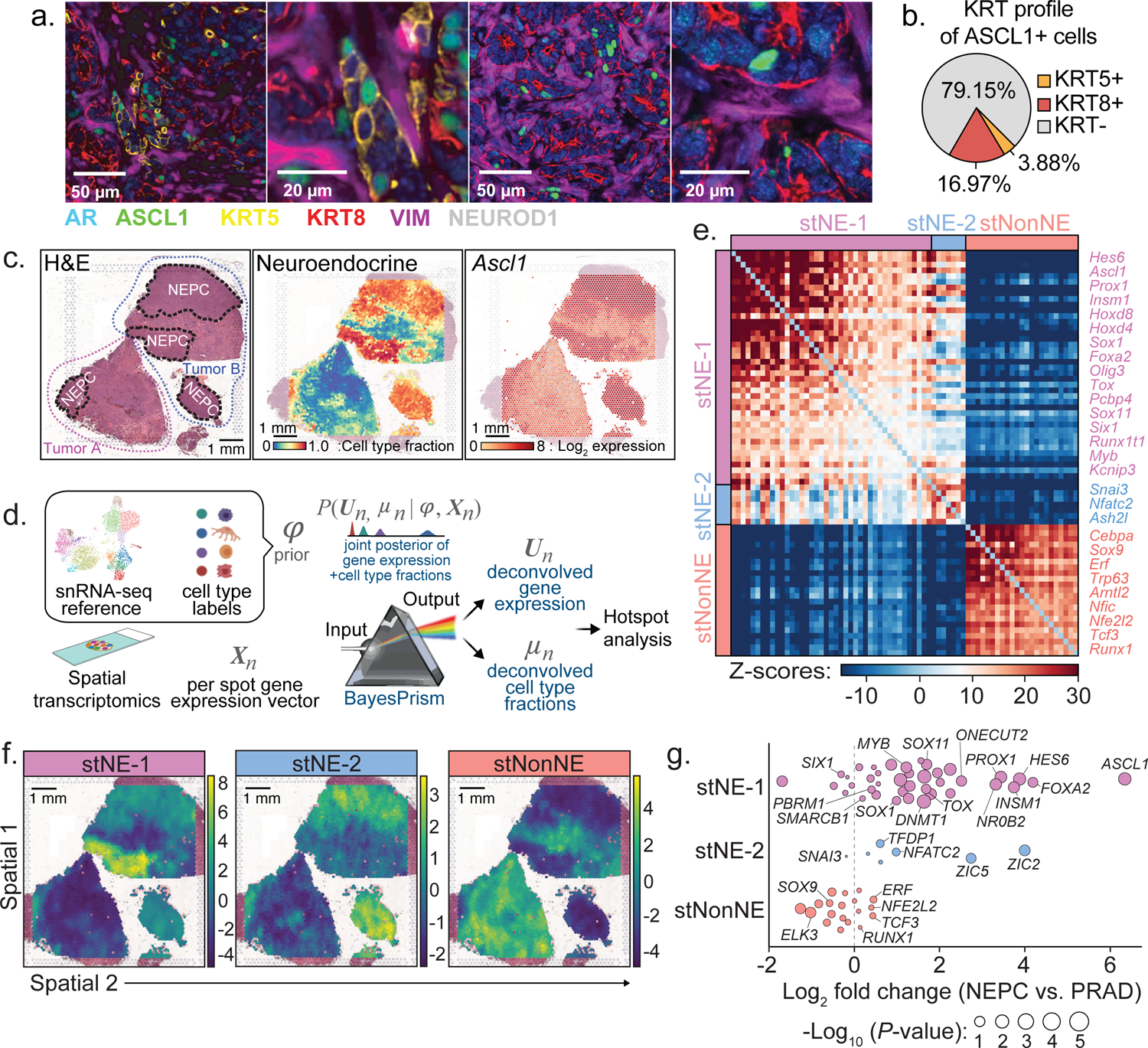
PrismSpot reveals spatial transcriptomic heterogeneity within NEPC marked by *Ascl1* co-expressed with distinct NE-related TFs. **a.** Representative confocal images of 7-plex IF of the indicated markers. Second and fourth images are digitally magnified versions of the first and third panel from the left. Data are representative of *n*=29 individual RPM tumors at varying time points post OT transplantation. **b.** Percentage of all ASCL1+ cells co-expressing KRT5, KRT8, or KRT-negative within an individual RPM OT tumor. Data is derived from the average percentage of cells within each tumor across *n*=10 independent tumors 6-weeks post OT transplantation. **c.** (Left) H&E stains of two independent 10-week RPM tumors. Tumors A and B are outlined in red and blue dotted lines, respectively. NEPC regions are highlighted in black dotted lines. (Middle) BayesPrism inferred cell type fraction for NEPC. (Right) Log_2_ fold expression of *Ascl1* overlayed on the tumor histology. **d.** Workflow of PrismSpot method. BayesPrism infers the posterior of cell type-specific gene expression, U, and cell type fraction, μ, of each spot. The expression profile of the cell type of interest (NEPC) was selected as the input for Hotspot analysis. **e.** Heatmap shows PrismSpot output of the pairwise local correlation *Z*-scores of 71 TFs of high consensus scores (>0.8) and significant spatial autocorrelation (FDR<0.01). TFs are clustered into 3 modules based on pairwise local correlations between all TFs of significant spatial autocorrelation. **f.** Spatial expression patterns of TFs within each module are visualized using smoothed summary module scores. **g.** Beeswarm plot shows the log2 fold change in expression of TFs in each module between bulk RNA-seq of human NEPC and PRAD samples. Dot size shows the two-sided *p*-values based on Wilcoxon test. All scale bars indicated within each figure panel.

The appearance of histologically homogeneous, spatially separate clusters of highly proliferative NE cells within weeks of detecting isolated ASCL1+;KRT8+ cells is consistent with a clonal expansion model. To further characterize the level of transcriptomic heterogeneity, we performed spatial transcriptomics (st; 10X Visium) using tissue sections containing both PRAD and NEPC, coupled with single cell nuclear RNA sequencing (snRNA-seq) from 10-week RPM tumors (**Fig. 5c and Supplementary** Fig. 3a). We observed distinct NE tumor cell clusters from snRNA-seq with variable KRT8 expression (**Supplementary** Fig. 3a-b), consistent with the evidence of heterogeneity within NEPC seen by multiplexed immunofluorescence.

Given the mixture of multiple cell types within individual tissue spots used for spatial transcriptomic sequencing, we applied BayesPrism^29,30^ to deconvolve tumor cell from non-tumor cell transcripts using the snRNA-seq data as the reference (**Fig. 5d**). BayesPrism integrates a single cell genomics reference with spatial transcriptomics data to deconvolve each spot into the cell type fractions present and provide a cell type specific count matrix for each spot, while accounting for differences between Visium and single cell reference. This method has superior performance in deconvolving spatial transcriptomics data using ground truth datasets^29,30^. Prior to deploying BayesPrism for further downstream analysis, we assessed the robustness of the inferred deconvolution by comparing BayesPrism on technical replicates profiled from adjacent tissues and found strong correspondence of inferred cell type fraction (**Supplementary** Fig. 4a-b). Specifically, the tumor cell type fraction inferred by BayesPrism recapitulates the distribution of NEPC observed by histology (**Fig. 5c**).

We next investigated the expression of TFs within regions with NEPC histology as well as those containing a high content of NEPC as inferred by BayesPrism. Consistent with its role regulating neuronal expression programs, all NEPC regions showed *Ascl1* expression with minimal *Neurod1* and *Pou2f3* expression (**Supplementary** Fig. 5).

Conversely, other TFs previously implicated in NEPC (e.g. *Mycn*, *Onecut2*, *Pou3f2*, *Pou3f4*) and cerebellar development (*Olig3*) were expressed only within subsets of the NEPC regions examined^16,31–34^ (**Supplementary** Fig. 5). The spatial heterogeneity in expression of these selected TFs, as well as similar TF heterogeneity reported in SCLC (a tumor of NE origin)^35–39^, led us to examine the structure underlying this heterogeneity using Hotspot^40^, which identifies spatially-varying genes. However, the limited resolution of Visium technology makes identification of gene modules specifically associated with a single cell type of interest challenging because direct application of Hotspot would detect co-localization of genes expressed within multiple cell types or between a pair of colocalized cell types, resulting in false positives when studying cell type-specific gene modules. To overcome this, we leveraged a powerful feature of BayesPrism: inference of cell-type specific count matrices, thereby associating each transcript with its respective cell type (**see Methods**). Therefore, as input to Hotspot, we used the deconvolved tumor count matrices, a strategy we have termed “PrismSpot” resulting from a combination of BayesPrism and Hotspot (**Fig. 5d**). Compared to directly applying Hotspot on un-deconvolved Visium data, the spatial auto- and pairwise-correlation computed by PrismSpot showed significantly stronger signal-to-noise ratio for tumor-specific gene modules (**Extended Data** Fig. 7a-g **and Supplementary** Fig. 6a-c). Application of PrismSpot identified five distinct spatial modules (**Supplementary Table 7**). To ensure robustness of the clustering of gene modules, we selected genes with the highest co-occurrence within each gene module upon iterative subsampling of the Visium data (**see Methods**), which narrowed down our gene list to 71 TFs spanning three of the original five modules (**Supplementary Table 7**).

We label these three final (robust) TF modules defining two NEPC states (stNE-1, stNE-2) and a single PRAD state (stNonNE; **Fig. 5e-f and Supplementary Table 7**). stNE-1, whose leading genes include coordinated expression of *Ascl1* and other TFs implicated in neuronal biology^41^ (*Hes6*, *Ascl1*, *Prox1*, *Insm1*), was enriched across all NEPC regions. The stNE-1 regions correspond to those with a high density of *Mycn* and *Olig3* expression (by spatial transcriptomics) and KRT8+;ASCL1+ (double-positive) tumor cells (by multiplexed IF; **Fig. 3b and Supplementary** Fig. 5). stNE-2, defined primarily by *Nfatc2* (a regulator of *Tox* expression within lymphocytes^42–44)^ but also includes the epithelial-to-mesenchymal (EMT) TF *Snai3* was enriched in some but not all NEPC regions (**Fig. 5f and Extended Data** Fig. 7h). Of note, *Nfatc2* expression has been linked with an EMT-like state in melanoma^45^. As further validation of these spatially derived signatures, both stNE modules are selectively enriched in the NEPC signature derived from previously reported scRNA-seq data of prostate GEMMs^13^ (**Extended Data** Fig. 7i) as well as human NEPC samples previously characterized using RNAseq^46^ (stNE-1 *p* = 1.17E-7, stNE-2 *p* = 5.50E-4, stNonNE *p* = 0.742, two-sided Wilcoxon test; **Fig. 5g and Supplementary Table 8**). Collectively, multiplexed IF and spatial transcriptomics combined with PrismSpot analysis suggest that NE differentiation arises from KRT8+ luminal epithelial cells which progressively evolve into spatially distinct ASCL1+ subpopulations with heterogeneous expression of other NE-associated TFs in various combinations.

### *Ascl1* is essential for NEPC transformation

In addition to its role as a master TF in neural lineage specification^47,48^, several human SCLC cell lines and at least one human NEPC xenograft model are dependent on ASCL1 for proliferation^39,49,50^. Whether ASCL1 upregulation is required during the transition from PRAD to NEPC progression is unknown. The reliable kinetics of disease progression in the RPM model, coupled with the flexibility to perform multiplexed genome editing, allow us to rapidly address this question through CRISPR editing of the *Ascl1* locus in RPM organoids (hereafter called *Ascl1^KO^*; **Supplementary Table 9 and Supplementary** Fig. 7a-b). To assess the requirement of *Ascl1* for NEPC transformation, we compared the growth and histologic features of *Ascl1^wt^* versus *Ascl1^KO^* RPM tumors following either OT or subcutaneous (SQ) transplantation (**Fig. 6a-d**). As expected, *Ascl1^wt^* RPM mice developed PRAD initially that, over 6-10 weeks, progressed to NEPC. Of note, we also observed a reproducible NE-lineage transition, with similar kinetics, following SQ injection, indicating that the *in vivo* signal that triggers lineage plasticity is not restricted to the prostate microenvironment. Multiplexed IF revealed that the TME of these SQ tumors shared many of the features seen in the OT tumors (**Supplementary** Fig. 8a-c). In stark contrast, all *Ascl1^KO^* RPM tumors (OT and SQ) developed PRAD with moderate to well-differentiated glandular histology, slower growth kinetics than *Ascl1^wt^* RPM tumors and, importantly, no evidence of NE transformation (**Fig. 6b-f and Extended Data** Fig. 8a-f). Furthermore, no metastases were detected in *Ascl1^KO^*RPM mice after OT, compared to 50% incidence in *Ascl1^wt^* RPM mice, despite comparable end-stage tumor weights at the primary OT site in either intact or castrated hosts (**Fig. 6g and Extended Data** Fig. 8g). Thus, *Ascl1* is obligately required for transition to NEPC and for metastasis in the RPM model.

**Figure 6:**
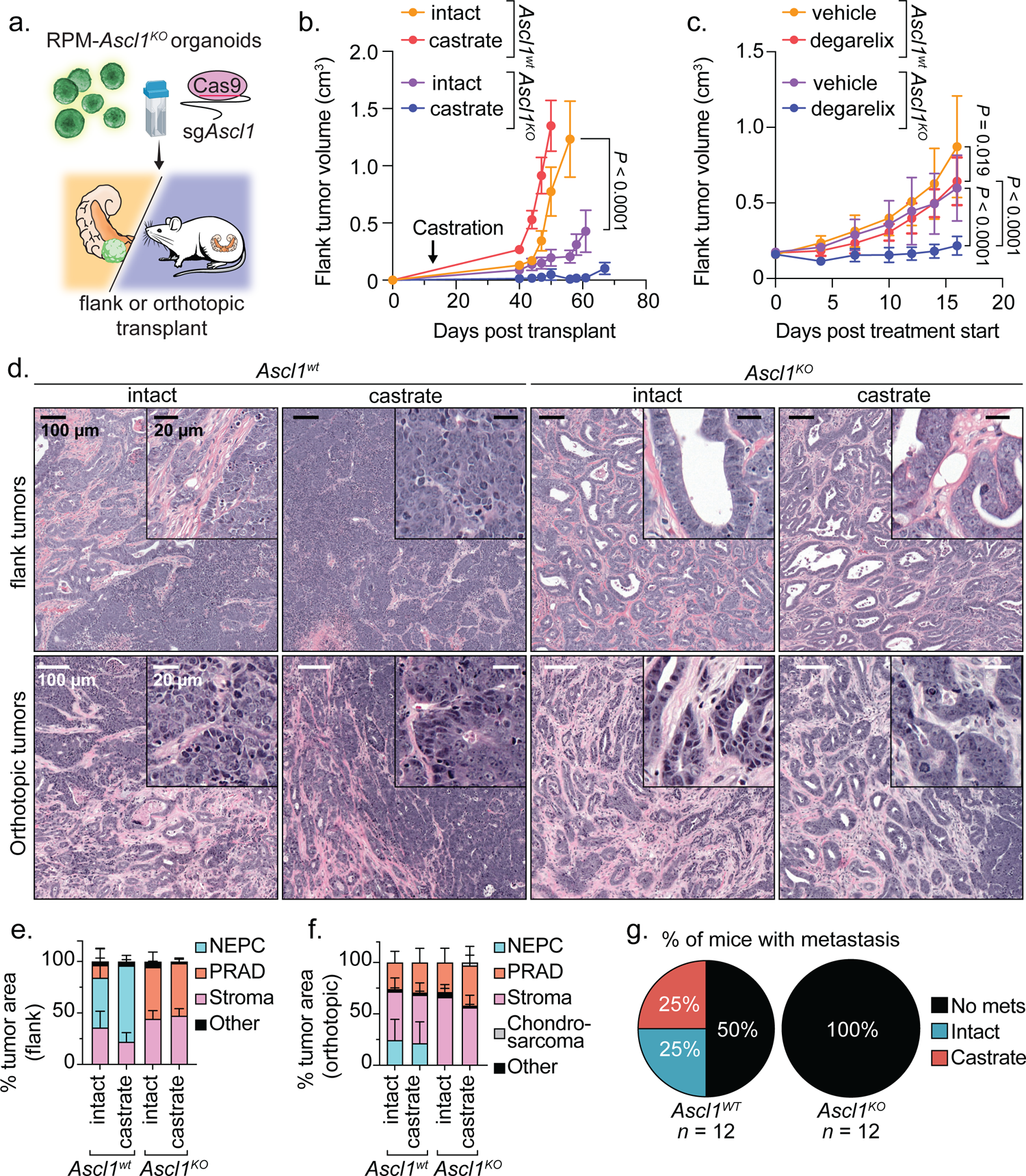
Loss of *Ascl1* results in abrogated NEPC establishment and castration-sensitivity. **a.** Schematic for the generation of RPM-*Ascl1^WT^*and RPM-*Ascl1^KO^* tumors transplanted into the flanks or prostates of immunocompetent C57BL6/J hosts. **b.** Longitudinal subcutaneous (SQ) tumor volumes of the indicated tumor genotypes and host backgrounds. Statistics derived using two-way ANOVA with Tukey’s multiple comparisons correction for data collected between days 0-56 to ensure equal sample size comparisons. Error bars denote mean and standard error of the mean. *n*=6 independent tumors across each group. Castration or sham surgery performed 14 days post SQ transplantation. **c.** Longitudinal SQ tumor volumes of the indicated tumor genotypes and host backgrounds. Statistics derived using two-way ANOVA with Tukey’s multiple comparisons correction for data collected between 0-16 days post treatment start to ensure equal sample size comparisons. Error bars denote mean and standard deviation. *Ascl1^WT^*-vehicle, *Ascl1^WT^*-vehicle, and *Ascl1^KO^*-degarelix, *n*=8; *Ascl1^KO^*-vehicle, *n*=9 independent tumors. Vehicle or degarelix treatment was initiated upon tumor establishment (≥150 mm^3^). **d.** Representative H&E of SQ (top) and OT (bottom) tumors isolated at endpoint. Genotype and treatment groups listed within the figure panel. Data related to mice in Fig. 5b-c. Scale bars denoted within the figure panel. Data are representative of 4-6 independent tumors per experimental group. **e.** Stacked bar charts representing percentage of OT tumor area composed of the histological categories depicted in the figure legend. Data are quantified histology of tumors generated in Fig. 5b and represent average tumor area. **f.** Stacked bar charts representing percentage of SQ tumor area composed of the histological categories depicted in the figure legend. Data are quantified histology of tumors generated in Fig. 5c and represent the average tumor area. **g.** Pie charts representing percentage of mice with metastatic disease (regional and distal) in intact or castrated hosts of the indicated genotypes. Statistics derived from two-sided Fisher’s exact test, *p*=0.0137. Number of biological replicates indicated in the figure panel. All scale bars denoted in the figure panels.

We and others previously found that perturbations preventing lineage plasticity may restore sensitivity to androgen deprivation therapy in prostate cancer^13,50^. To address if this is also true in the context of *Ascl1* loss, we compared the tumorigenicity and histologic features of *Ascl1^wt^* and *Ascl1^KO^* RPM tumors following OT or SQ injection into intact versus castrated mice. Of note, *Ascl1^KO^*tumors grew significantly slower in castrated versus intact hosts in both the OT and SQ settings, an intriguing result given that loss of *RB1* and *TP53* are strongly linked to castration-resistance in multiple prostate models and in patients (**Fig. 6b and Extended Data** Fig. 8a-f). To distinguish between effects of castration on tumor engraftment versus tumor maintenance, we initiated chemical castration therapy (degarelix) in established SQ tumors (≥150mm^3^). Degarelix treatment completely abrogated the growth of *Ascl1^KO^*RPM tumors and significantly extended survival whereas progression of *Ascl1^WT^* RPM tumors was only marginally impacted (**Fig. 6c, Extended Data** Fig. 8h-i**, and Supplementary** Fig. 9). Interestingly, one castrated mouse injected with *Ascl1^KO^* RPM organoids developed a tumor with chondrocyte-like histology, reminiscent of a similar phenotype reported in RPM-driven; *Ascl1^KO^*SCLC mouse models^49^ (**Extended Data** Fig. 8j).

To investigate why tumors with *Rb1* and *Trp53* loss display increased androgen dependence in the context of *Ascl1* loss, we examined the expression of AR as well as luminal (KRT8) and basal (KRT5) cytokeratins. Consistent with their well-differentiated glandular morphology, RPM-*Ascl1^KO^* tumors were dominated by AR+;KRT8+ tumor cells (**Fig. 7a and Extended Data** Fig. 9a-c). Furthermore, the intensity of nuclear AR staining was significantly elevated relative to RPM-*Ascl1^WT^* tumors (**Fig. 7b-d and Extended Data** Fig. 9d-f). Notably, degarelix treated RPM SQ tumors harbored an increase in NEPC tumor area as measured by histological evaluation (**see Methods**), with an increase in the fraction of ASCL1+, but decreased AR+ tumor cells relative to vehicle treated RPM SQ tumors (**Extended Data. Fig. 8k**). Taken together, these data suggest that *Ascl1^KO^* tumors are phenotypically and transcriptionally bottlenecked into a luminal AR-dependent state.

**Figure 7:**
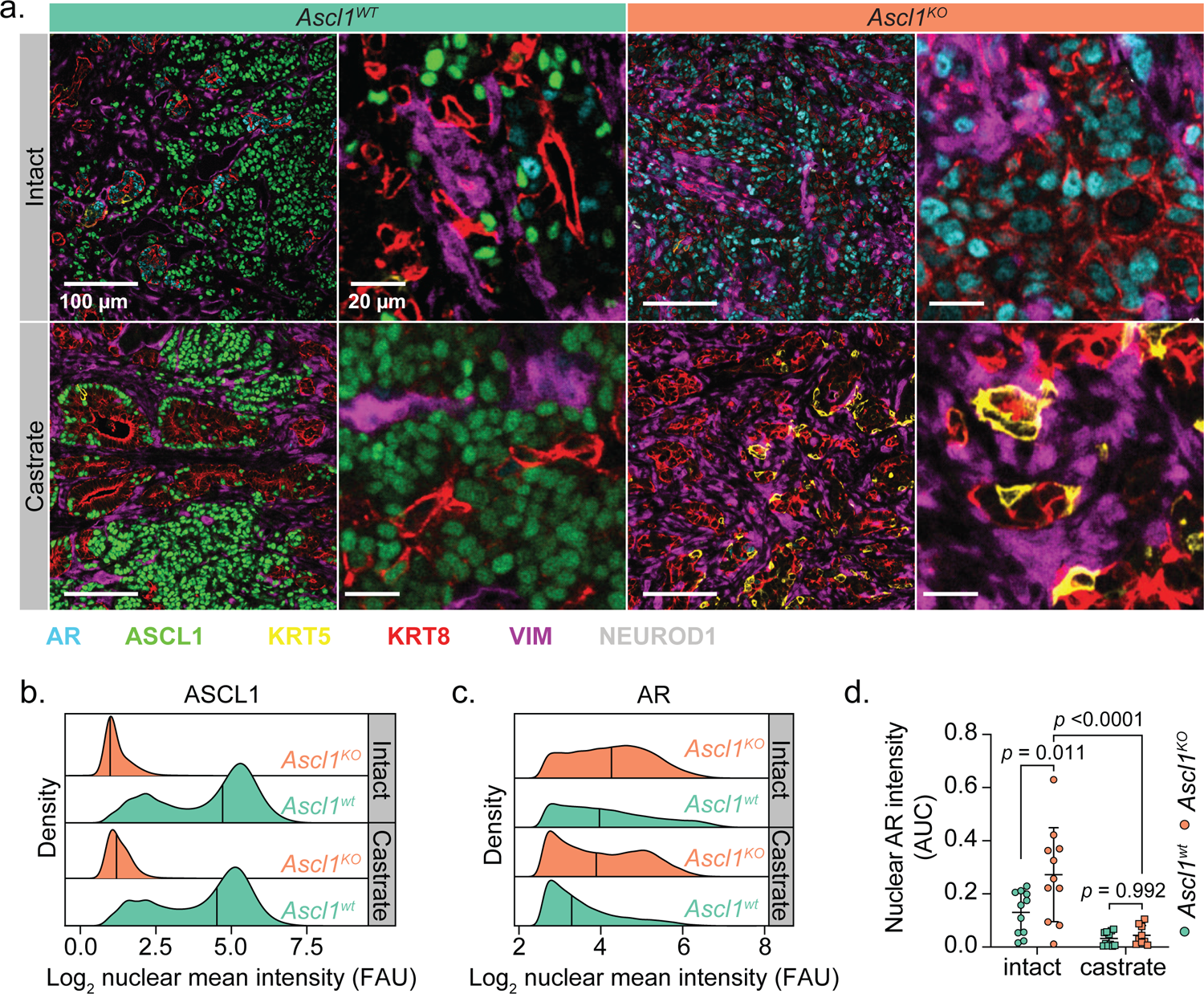
Loss of *Ascl1* results in enhanced AR expression and proportion of KRT8+ tumor cells. **a.** Representative confocal images of the tumors isolated from mice in Fig. 5b-c. Scale bars and pseudocolor legend indicated within the figure. **b.** Density plots of the log_2_(x+1) transformed ASCL1 mean fluorescence intensity from all (OT and SQ) tumor cells. Tumor cells subset by all cells staining negatively for VIMENTIN. Tumor genotype and treatment indicated in the figure panel. Data derived from *n*<10 independent RPM tumors per group. **c.** Density plots of the log_2_(X+1) transformed AR mean fluorescence intensity from all OT tumor cells within the indicated genotypes and treatment groups. Tumor cells subset by all cells staining negatively for VIMENTIN and positively for KRT8 and AR. **d.** Area under the curve for all KRT8+:AR+ tumor cells (VIMENTIN-) across both SQ and OT tumor transplants, containing a log_2_ transformed nuclear AR intensity score ≥3. Statistics derived using two-way ANOVA with Tukey’s multiple comparisons correction. Error bars indicate mean and standard deviation. Combined OT and SQ tumor sample sizes for all quantification and analysis performed in Fig. 7: *n*=11 (*Ascl1^wt^* and *Ascl1^KO^*intact groups), *n*=12 (*Ascl1^wt^* castrate group), *n*=9 (*Ascl1^KO^* castrate group). FAU=fluorescence arbitrary units.

**Figure 8:**
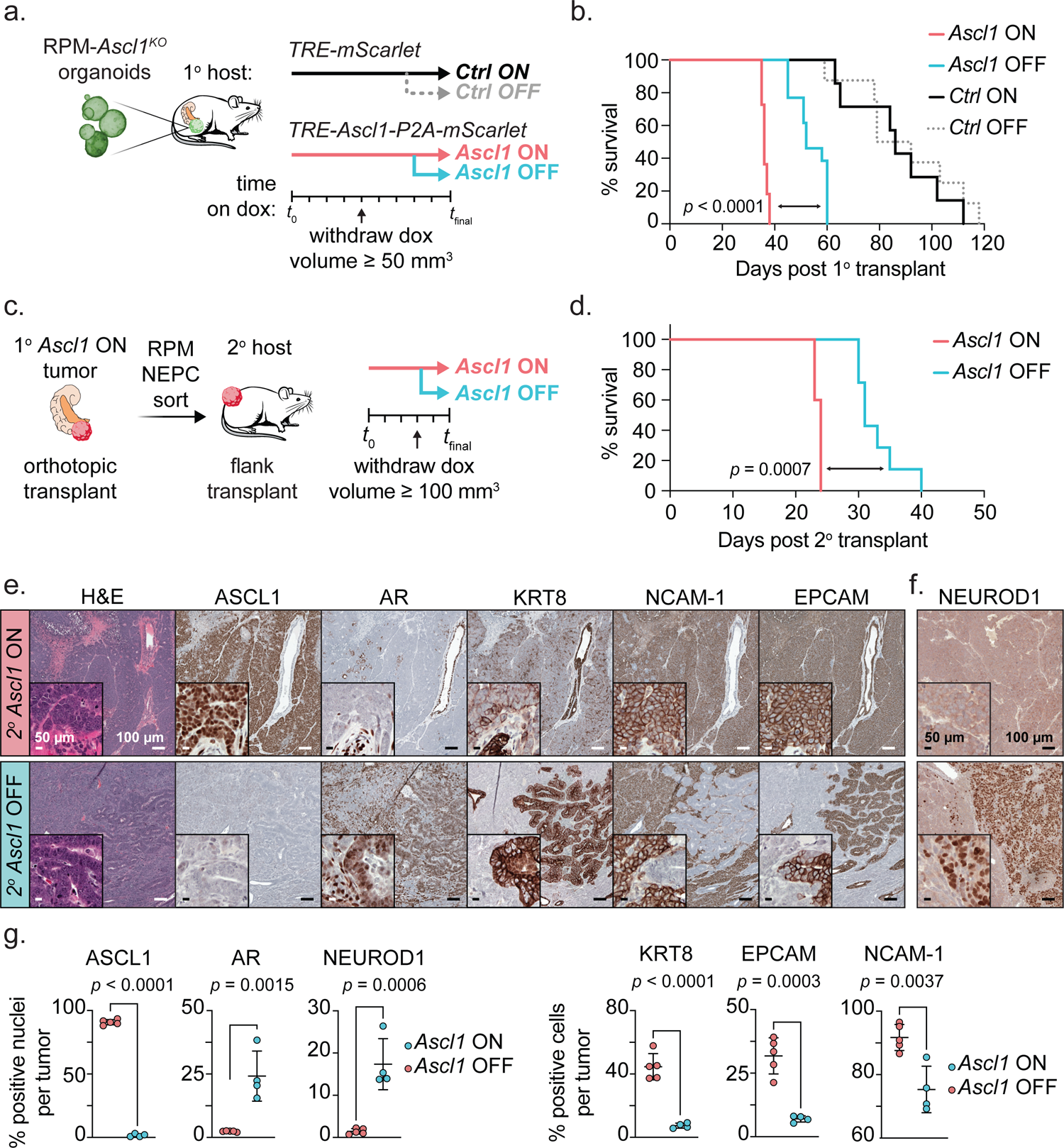
Loss of *Ascl1* in established NEPC results in modest tumor control and increased tumor heterogeneity. **a.** Schematic of *Ascl1* doxycycline (dox)-inducible *in vivo* platform. RPM-*Ascl1^KO^* organoids infected with inducible *mScarlet* (*Ctrl*) or *Ascl1*-*P2A-mScarlet* (*Ascl1*) vectors were transplanted OT into mice fed dox-chow (primary recipient host, 1°). Tumor volume was monitored by ultrasound. Upon primary tumor establishment, mice were randomized into dox ON (maintained) or dox OFF (withdrawal) groups. **b.** Survival curves of *Ctrl* or *Ascl1* induced OT tumors following dox-maintenance (ON groups) or withdrawal groups (OFF groups). Statistics derived from log-rank (Mantel-Cox) test comparing primary *Ascl1* ON to primary *Ascl1* OFF groups. *Ctrl* ON *n*=7, *Ctrl* OFF *n*=8, *Ascl1* ON *n*=11, *Ascl1* OFF *n*=13 independent mice. **c.** Schematic of SQ *Ascl1* doxycycline (dox)-inducible *in vivo* platform (secondary recipient host, 2°). *Ascl1 ON* primary tumors were dissociated for flow cytometry to enrich for RPM-NEPC cells used for transplantation assays into the flanks of secondary recipient mice fed dox-chow. Tumor volume was monitored by caliper. Upon tumor establishment, mice were randomized into dox ON (maintained) or dox OFF (withdrawal) groups. **d.** Survival curves of *Ctrl* or *Ascl1* induced secondary tumors following dox-maintenance (ON groups) or withdrawal groups (OFF). Stats derived from Log-rank (Mantel-Cox) test. *Ascl1* ON *n*=5, *Ascl1* OFF *n*=7 independent mice per group. **e.** Serial sections from secondary transplanted mice (SQ) stained for the indicated markers by H&E and IHC. **f.** Representative NEUROD1 IHC within a secondary transplant containing mostly NEPC histology. Data representative of *n*=5 independent tumors. **g.** (Left) Average percent marker positive nuclei or (right) cells across biologically independent secondary SQ *Ascl1* ON (*n*=5) or OFF (*n*=4) tumors. Statistics derived from two-sided *t*-test. Error bars indicate mean and standard deviation. All scale bars depicted in the figure panels.

### Loss of Ascl1 in established NEPC promotes tumor heterogeneity

Given the exquisite dependence on *Ascl1* for the transition to NEPC, we next asked if *Ascl1* is also required for the maintenance of established RPM-NEPC tumors. To address this question, we introduced a doxycycline (Dox) regulatable *Ascl1* cDNA (with a cis-linked *mScarlet* reporter allele) into RPM-*Ascl1^KO^* organoids and performed OT experiments in mice receiving Dox (**Fig. 8a and Extended Data** Fig. 10a). As expected, mScarlet-positive OT primary tumors developed quickly (within 5 weeks) in mice transplanted with RPM-*Ascl1^KO^* organoids harboring the Dox-*Ascl1* allele (hereafter *Ascl1^ON^*) whereas tumors in mice transplanted with RPM-*Ascl1^KO^* organoids containing the Dox-*mScarlet* allele alone were delayed (*Ctrl^ON^*) (**Fig. 8b and Extended Data** Fig. 10b-c). *Ascl1^ON^* mice also developed metastases whereas *Ctrl^ON^* mice did not (**Extended Data** Fig. 10d), thus fully recapitulating the findings reported earlier (**Fig. 6g**).

Having established the fidelity of the Dox-*Ascl1* rescue allele, we asked if ASCL1 is required for the sustained growth of *Ascl1^ON^* tumors in a second cohort of *Ascl1^ON^* mice and *Ctrl^ON^* mice that received Dox until tumors were established (≥100 mm^3^) followed by withdrawal (hereafter *Ascl1^OFF^*and *Ctrl^OFF^*; **Fig. 8a and Extended Data** Fig. 10b). Consistent with prior evidence that *ASCL1*-knockdown delays the growth of human NEPC xenografts, most *Ascl1^OFF^*tumors regressed within one week of Dox withdrawal but resumed growth within 2-3 weeks. Although short lived, Dox withdrawal resulted in a statistically significant (albeit modest) survival benefit (log-rank Mantel-Cox test, *p* < 0.0001; **Fig. 8b and Extended Data** Fig. 10e-f).

To gain insight into the mechanism of relapse after Dox withdrawal, we examined the histologic features and lineage of relapsed *Ascl1^OFF^*tumors. To avoid the confounding issue of PRAD cells within RPM primary transplants (recall that RPM tumors retain mixed PRAD and NEPC histology), we focused our analysis solely on NEPC cells by isolating a pure population of RPM-NEPC from primary *Ascl1^ON^* OT tumors then retransplanting these cells SQ into secondary recipients (**Fig. 8c and Supplementary** Fig. 10; **see Methods**). As expected, the SQ transplants mirrored the results seen by OT in that *Ascl1^ON^* tumors grew rapidly, whereas *Ascl1^OFF^* tumors had slower growth with a modest but significant extension in survival (log-rank Mantel-Cox test, *p* = 0.0007; **Fig. 8d and Extended Data** Fig. 10g). As expected, *Ascl1^OFF^* tumors lacked nuclear ASCL1 expression, confirming the fidelity of the Dox^ON^/Dox^OFF^ platform (**Fig. 8e)**. While loss of ASCL1 expression resulted in the reacquisition of some histologic features of PRAD (with pockets of moderate- to well-differentiated adenocarcinoma harboring KRT8 and AR expression), the predominant histologic phenotypes were high-grade ASCL1-NEPC and regions of sarcomatoid-like differentiation (**Fig. 8e-g, Extended Data** Fig. 10h**, and Supplementary** Fig. 11a-c). In contrast to the RPM tumors discussed earlier where we found no evidence of NEUROD1 expression (**Fig. 2b**), we now observed several regions of NEUROD1+ NEPC (**Fig. 8f-g and Extended Data** Fig. 10i). In summary, and in contrast to clear dependency on *Ascl1* for the initiation of NE plasticity, *Ascl1*-dependency in established NEPC tumors is rapidly circumvented, revealing unique pathologies and marker profiles not seen previously in RPM or RPM-*Ascl1^KO^* tumors. Moreover, these results provide evidence of selective pressure to maintain the NE state through upregulation of NEUROD1 and perhaps other TF programs that remain to be identified.

## DISCUSSION

Because lineage plasticity in cancer is a dynamic process that evolves over time, a precise understanding of the underlying molecular events requires a model amenable not only to repetitive interrogation but also rapid perturbation and reconstitution of the full repertoire of cells found within the TME. Through application of organoid techniques, genome engineering and *in vivo* transplantation assays, we have generated a scalable, flexible, and robust platform that captures the evolution from PRAD to NEPC with highly reproducible kinetics in a manner that closely resembles human disease. As with human NEPC, the mouse NEPC transition is accelerated by castration, although it is worth noting that plasticity also occurs in hormonally intact mice. Using this platform, we have identified at least two steps that are required for plasticity to develop. The first is *Rb1* loss which we postulate creates a cell state poised for lineage transformation. This is followed by a second “trigger” derived from the TME that initiates and cooperates with lineage-defining TFs such as *Ascl1* to complete the transition from an epithelial to NE lineage. Detailed characterization of the chromatin state of tumor cells in this model prior to and during the lineage transition, coupled with side-by-side analysis of signaling crosstalk with the TME in *Rb1*^-/-^ versus *Rb1*^+/+^ backgrounds (e.g., RPM versus PtPM) and cross referenced with published *Rb1* chromatin residence data should shed light on the underlying molecular events^51^.

Application of spatial methods to this model allowed us to gain additional insight into the origin of NEPC and its subsequent evolution as well as changes to the local TME. For example, the earliest detectable ASCL1+ cells nearly always co-express KRT8 or are adjacent to KRT8+ epithelial cells. In addition to implicating luminal cells as a likely cell of origin, this may provide an important clue as to the source of the TME trigger. Our spatial analysis also allowed us to track the expansion of ASCL1+ cells following the initial lineage transformation event, where we see further evolution into transcriptionally distinct NE clusters that continue to express ASCL1 but now gain expression of other TFs associated with neural lineage development. This NEPC evolution is also associated with substantial changes in the TME, such as near complete loss of mesenchymal cells and loss of infiltrating CD8+ T cells and CD4+ Tregs.

In addition to the unique capability of capturing critical aspects of lineage plasticity that are not recapitulated *in vitro*, the platform is well positioned for rapid throughput functional studies. For example, application of multiplexed gene editing at the time of tumor initiation established the critical role of ASCL1 in NE transformation in a matter of months (versus 1-2 years required for multigenic crosses using GEMMs). The role of ASCL1 in the development and maintenance of NE cancers has been previously addressed in the context of SCLC but, importantly, that dependency is a consequence of tumor initiation in pre-existing ASCL1+ NE cells^39,49^. By contrast, ASCL1 is not expressed in prostate cancer except during the epithelial to NE lineage transition. While prior work has shown delayed growth of ASCL1-expressing human xenograft models following *ASCL1* knockdown^50^, the dynamic nature of our platform allowed us to document an essential role of *Ascl1* in initiating the transformation of PRAD to NEPC. Deletion of *Ascl1* prior to histological transformation resulted in homogeneous well-differentiated adenocarcinomas with no evidence of escape to another lineage despite *Rb1/Trp53* loss and *c-Myc* overexpression.

In addition to evaluating the importance of genes such as *Ascl1* in initiating the lineage plasticity program, the model is also well positioned, through use of a Dox-inducible rescue alleles, to address dependencies on such genes once NEPC is fully established. In contrast to the absolute dependence on *Ascl1* for the NEPC transition, termination of *Ascl1* expression within established NEPC resulted in transient tumor regressions. Notably, the relapsed tumors contained small foci of AR+, KRT8+ PRAD (indicative of some lineage reversion) but are dominated by sarcomatoid features and regions of NEUROD1+ NEPC that rapidly progress. In addition to revealing additional layers of phenotypic plasticity, this result underscores the advantage of early pharmacologic intervention to prevent plasticity rather than intervening after plasticity is fully established. Whether such an approach is clinically feasible with ASCL1 remains to be determined as we are unaware of any drug development efforts that have succeeded in directly targeting ASCL1. However, clinical benefit has been reported using a bi-specific T cell engager targeting the downstream cell surface protein DLL3 in SCLC, and other DLL3-directed radio-ligand and cell-based therapies are also in development^52–54^.

The establishment of this model in a fully immunocompetent setting provides an opportunity to address several unresolved topics regarding the immunobiology of prostate cancer. In contrast to cell lines derived from tumors that have escaped immune suppression (and are commonly used to evaluate novel immunotherapies), the immune evasive mechanisms in the current model develop without any pre-transplantation immune-mediated selective pressure. This scenario allows deeper analysis of the earliest steps in immune escape and may shed light on novel strategies to buttress immunity before tumors become depleted of T cells. Indeed, our spatial analysis shows that CD8 T cells are present early in PRAD but absent in NEPC. We hope to unravel these details using prostate tumors expressing model antigens, combined with tetramer-based monitoring of T cell responses and selective depletion of specific myeloid and Treg subpopulations.

Although the work reported here is exclusively based on prostate cancer models, the platform is, in principle, adaptable to other epithelial lineages in which short-term organoid culture and orthotopic transplantation methods have been developed. One disease that closely approximates the lineage transitions observed in prostate cancer is *EGFR*- or *ALK*-mutant lung adenocarcinoma where epithelial to NE transition is seen as a mechanism of escape from EGFR or ALK inhibition, particularly in patients with co-occurring loss of function mutations in *TP53* and *RB1*, and recently demonstrated in an EGFR-driven GEMM^7,8,11,55,56^. *KRAS^G12C^*-mutant lung adenocarcinoma is a second example where transition to squamous histology is a resistance mechanism for RAS inhibitor therapy^10,57^. Other applications in bladder, pancreas, breast, and gastrointestinal cancer can also be easily envisioned. In closing, we report a robust, scalable platform to study lineage plasticity in a format amenable to deep molecular interrogation and perturbation and identify *Ascl1* as a critical gatekeeper of NE transformation and tumor heterogeneity in prostate cancer.

## Supporting information

Source Data File: Unedited western blots

## ACKNOWLEDGEMENTS

We thank members of the Sawyers and Pe’er labs for valuable discussions. We thank Y.M. Soto-Feliciano, F.J. Sánchez-Rivera, T. Tammela, T. Papagiannakopoulos, C. Concepcion-Crisol, S. Naranjo, E.E. Gardner, B.D. Stein, C. Burdziak, P. McGillivray, and T. Baslan for their critical feedback. We thank the core facilities at MSKCC for their support, including the Molecular Cytology Core, Flow Cytometry Core, Mouse facilities, and N. Mao and J. Lawson for their administrative support. R.R. was supported by NIH Translational Research Oncology Training Program (T32CA160001) and Charles H. Revson Senior Fellowship in Biomedical Science (22-23). T.C. was supported by William Raveis Charitable Fund Damon Runyon Quantitative Biology postdoctoral fellowship (DRQ: 10-21). T.J.G.R. was supported by NIH Institutional Cell Biology training grant (T32GM136542) and Howard Hughes Medical Institute Gilliam Fellowship program (GT15758). S.Z. is supported by NIH K08 CA282978 and Burroughs Wellcome Fund Career Award for Medical Scientists. We acknowledge the use of the Integrated Genomics Operation Core, funded by the NCI Cancer Center Support Grant (CCSG, P30 CA08748), Cycle for Survival, Marie-Josée and Henry R. Kravis Center for Molecular Oncology and the Alan and Sandra Gerry Metastasis and Tumor Ecosystems Center. This work was supported by the Howard Hughes Medical Institute; Calico Life Sciences; NIH grants CA193837, CA092629, CA224079, CA155169, CA008748, and CA274492. The content is solely the responsibility of the authors and does not necessarily represent the official views of the National Institutes of Health, Howard Hughes Medical Institute, Damon Runyon Cancer Research Foundation, or the Charles H. Revson Foundation. D.P. and C.L.S. are Howard Hughes Medical Institute investigators.

## AUTHOR CONTRIBUTIONS

R.R. and C.L.S. designed the study and wrote the manuscript with comments from all authors. R.R. designed, analyzed, and oversaw all experiments. R.R., P.S., K.E.L., H.K., and O.G. performed experiments. R.R., P.S., and K.E.L. performed mouse work. T.C., T.J.G.R, and Y.X. performed computational analysis. T.C. developed PrismSpot, analyzed snRNA-seq, scRNA-seq and spatial transcriptomics data. T.J.G.R. performed bulk RNA-sequencing and immunofluorescence spatial analysis. Y.X. analyzed COMET spatial immunofluorescence datasets. H.Z. and E.D. performed or oversaw orthotopic surgeries, respectively. S.Y. and R.R. optimized Lunaphore COMET multiplexed immunofluorescence. C.M., W.K., and M.V.P. performed 7-plex immunofluorescence. R.R. performed immunohistochemical staining and confocal microscopy. A.G. assessed and cross-validated histopathology and grade of tumors. S.Z., K.Y., and J.C. performed macrophage subset scRNAseq analyses across human prostate tumor samples. N.F. performed tissue embedding and preparation for 10X Visium spatial transcriptomics. W.R.K. was involved in optimization of improved organoid culture methods. K.V.R. oversaw bulk RNA sequencing analysis. P.M.K.W. was involved in optimization of organoid transplantation assays. D.P. oversaw all snRNA-seq, spatial transcriptomics, and multiplexed immunofluorescence analyses. D.P. and C.L.S. oversaw the project.

## COMPETING INTERESTS

C.L.S. is on the board of directors of Novartis, is a cofounder of ORIC Pharmaceuticals, and is a co-inventor of the prostate cancer drugs enzalutamide and apalutamide, covered by US patents 7,709,517; 8,183,274; 9,126,941; 8,445,507; 8,802,689; and 9,388,159 filed by the University of California. C.L.S. is on the scientific advisory boards for the following biotechnology companies: Beigene, Blueprint Medicines, Column Group, Foghorn, Housey Pharma, Nextech, PMV Pharma and ORIC. D.P. is on the scientific advisory board of Insitro.

EXTENDED DATA FIGURES 1-10

SUPPLEMENTARY FIGURES 1-11

**Supplementary Table 1:** Mutational co-occurrence analyses in prostate cancer patients for all genetic combinations modeled in mouse prostate organoids.

**Supplementary Table 2:** Summary table of organoid genotypes and in vivo transplantation efficiency and histology.

**Supplementary Table 3:** PtPM and RPM tumor bulk RNAseq normalized (log_2_(X+1)) read counts. Related to Fig. 2c-d and Extended Data Fig. 5e.

**Supplementary Table 4:** Gene set enrichment analysis (REACTOME) results from RPM vs PtPM tumors processed for bulk RNAseq. Related to Supplementary Table 3.

**Supplementary Table 5:** Antibodies used for multiplexed immunofluorescence (Lunaphore COMET).

**Supplementary Table 6:** Marker co-expression used for cell typing COMET datasets.

**Supplementary Table 7:** PrismSpot module analysis. Related to Fig. 5e.

**Supplementary Table 8**: PrismSpot consensus TF differential gene expression in human NEPC versus PRAD.

**Supplementary Table 9:** *Ascl1* edited alleles in RPM-*Ascl1^KO^*pooled clones.

**Supplementary Table 10:** Antibodies and concentrations used for IHC, multiplexed IF, western blot, and flow cytometry.

**Supplementary Table 11:** Supplementary sequences.

**Source Data Figure 1:** Unedited western blot scans (related to Extended Data Fig. 2c).

## METHODS

### Mice

Animal studies were carried out in full compliance with Research Animal Resource Center guidelines and the MSKCC Institutional Animal Care and Use Committee under protocol #06-07-012). Only male mice were used for transplantation studies. All mice used for transplantation harbored conditional EGFP alleles to tolerize against EGFP-derived antigens expressed within organoids (Jax, #026179). All studies employed ≥ 3 animals per genotype per experimental cohort. Mice were maintained on a pure C57BL/6J genetic background. At established experimental end-point, mice were euthanized by CO_2_ asphyxiation followed by cervical dislocation.

### Orthotopic prostatic organoid transplantation

For tumor transplant studies, 8-12-week-old animals with appropriate genotypes were randomized for surgical implantation of *ex-vivo* manipulated organoids as described previously^20^. Briefly, the ventral abdomen was depilated (using clippers) the day prior to surgery. Animals were anesthetized with isoflurane and the surgical area was disinfected three times with alternating Betadine/Isopropanol. Eye lubrication was used to maintain eye health. Sterile tools were used for all procedures. A 0.5 cm midline incision was made along the lower abdominal midline and peritoneal wall to allow for exteriorization of the bladder, prostate, and seminal vesicles. Local analgesia was used at the incision site (bupivacaine). Using straight forceps, the bladder was gently pressed down caudally, exposing the dorsal prostate lobes. A 30-gauge needle was inserted into the right dorsal prostatic parenchyma and 20 μL (containing single cell suspensions of organoids in 50% PBS + 50% matrigel) was injected. Organoid/matrigel mix was kept on ice throughout the entire procedure. Successful injection was visualized by local expansion in volume of the right dorsal lobe without leakage. The prostate, bladder, and seminal vesicles were gently internalized, and the peritoneal layer was sutured using 5-0 vicryl sutures. The outer skin layer was closed with 3 to 4 wound clips. All mice received pre- and post-operative analgesia with buprenorphine and meloxicam and were followed post-operatively for any signs of discomfort or distress. Tumor-containing area was measured on hematoxylin and eosin (H&E) slides. Histological assessment was performed in consultation with clinical pathologist Dr. Anuradha Gopalan.

### Subcutaneous prostatic organoid transplantation

For allograft experiments, 250k single cell suspension of organoids in 100 μL 50% + 50% matrigel were injected into the depilated flanks (clippers) of isoflurane anesthetized C57BL/6J mice. Mice were followed for tumor measurement and signs of discomfort or distress. Subcutaneous tumor volumes (mm^3^) were calculated using the formula: (*a*^2^\**b*)*(π/6), where *a* and *b* are the smaller and larger dimensions, respectively.

### Castration surgery

Mice harboring orthotopic prostate tumors were randomized into castration or sham surgeries 2 weeks post orthotopic surgery. Mice were anesthetized with isoflurane. Eye lubrication was used to maintain eye health. The perineal region was cleaned three times with alternating Betadine/Isopropanol. Sterile tools were used for the procedure. A 0.5 cm incision was made in the scrotal sack. Forceps were used to locate and exteriorize the testes. Using a cauterizing iron, the testes were amputated via the seminal tubules. The scrotum was closed shut with 2-3 wound clips. A local triple antibiotic was applied over the region to facilitate healing. All mice received pre- and post-operative analgesia with meloxicam and were followed post-operatively for any signs of discomfort or distress.

### Chemical castration studies (GnRH antagonist)

1E6 RPM-*Ascl1^WT^* or RPM-*Ascl1^KO^* organoids were injected into the flanks (one tumor per mouse) of immunocompetent mice as described above. Mice were randomized into vehicle (5% D-Mannitol, Sigma M4125-500G) or Degarelix (15 mg/kg, Sigma SML2856-25MG) groups once tumors measured ≥ 150 mm^3^. Treatment was performed by subcutaneous delivery once every 14-days in a total of 100 uL. Tumor volumes were obtained and calculated as above. At sacrifice, blood was collected from mice and coagulated for 1hr on ice, followed by centrifugation at 1000 g for 30 min to isolate serum. Successful serum testosterone depletion was assessed by ELISA according to manufacturer recommendations (Abcam, ab285350).

### Transplantation of dox inducible Ascl1 organoids

100E5 RPM-*Ascl1^KO^* organoids harboring dox-inducible *Ascl1-P2A-mScarlet* or *mScarlet* alone were injected (50% matrigel) into the prostates of C57BL/6J *Prkcd^KO^* mice (Jax, #001913) to avoid rtTA-mediated rejecton. Dox chow (Inotiv, 0.625 g/kg) was started 1 week prior to transplantation ensure immediate induction of transgene expression. Mice were randomized into *Ascl1^ON^* (maintained on dox) or *Ascl1^OFF^* (withdrawn from dox) cohorts when tumor volumes reached ≥ 50mm^3^, as measured by small animal ultrasound. For secondary transplantation assays of pure RPM-NEPC tumor cells, 5-week *Ascl1^ON^* tumors were harvested as described for primary prostate organoid single cell suspensions, and sorted (Sony MA900, Sony Biotechnology, 100-μm sorting chip (Sony Biotechnology, #LE-C3210) for DAPI-, EGFP+, mScarlet+ (*Ascl1* reporter), EPCAM+, NCAM-1+ cells (gating strategy **Supplementary** Fig. 10). Antibodies used for flow cytometry listed in **Supplementary Table 10**. Post sort, 100E5 cells were immediately injected (50% matrigel) into the flanks of secondary C57BL/6J *Prkcd^KO^*mice pre-fed dox diet for 1 week pre-transplantation. Tumor volumes were assessed by caliper measurements as above. Mice were randomly separated into *Ascl1^ON^* or *Ascl1^OFF^* groups when flank tumors reached ≥ 50mm^3^. At experimental end-point, secondary tumors from *Ascl1^ON^* or *Ascl1^OFF^*groups as well as RPM organoids and primary *Ctrl* ON tumors (mScarlet-induced expression alone) were harvested for FFPE and processed for flow-cytometry for the markers listed above (**Supplementary** Fig. 11a-c).

### Small animal ultrasound

Animals were anesthetized using isoflurane and the ventral abdominal areas were depilated with Nair. Eye lubrication was used to maintain eye health. Animals were imaged using the Vevo2100 ultrasound and photoacoustic imaging system (Fujifilm-Visualsonics). Animals were placed on the imaging platform in the supine position and a layer of ultrasound gel was applied over the entirety of the abdominal area. The ultrasound transducer was placed on the abdomen orthogonal to the plane of the imaging platform. The bladder and urethra were used as landmark organs to define the area of the prostate. The transducer was set at the scanning midpoint of the normal prostate or prostatic tumor and a 3D image of 10-20 mm, depending on tumor size, at a Z-slice thickness of 0.04mm. 3D images were uploaded to the Vevo Lab Software and volumetric analysis function was used to determine the tumor border at various Z-slices through the entirety of the tumor and derive the final calculated tumor volume.

### Human Specimens

Informed consent was obtained for all patients and approved by MSKCC’s Institutional Review Board (IRB) #12-245 (NCT: 01775072) and #06-107. The human prostate tumor specimen was collected from a 62-year-old male with localized PRAD undergoing XRT followed by salvage prostatectomy post ADT and docetaxel. Tumor in the bladder arose by extension of a prostate tumor recurrence in the surgical bed. Pathological evaluation showed small cell carcinoma arising from PRAD. Tumor cells were focally positive for SYP, CHGA (patchy), PSA (focal), and PSMA (focal weak). Tumor sample was sectioned and processed for COMET-based multiplexed IF according to the antibodies listed in **Supplementary Table 10**.

### Immunohistochemistry and Immunofluorescence

#### Immunohistochemistry

Prostate tumors were cleaned under a stereomicroscope, fixed overnight in zinc-formalin, washed in PBS, transferred to 70% ethanol, and embedded in paraffin. Sections were cut to six micrometers and stained with H&E. Chromogenic immunohistochemistry (IHC) was performed on fresh cut sections. Briefly, slides were heated for 30 min at 58°C and deparaffinized. Antigen retrieval was performed in freshly prepared citrate buffer (pH 6.0) followed by Tris-EDTA (pH 9.0, Abcam #ab93684) in a decloaking chamber and subsequently slowly cooled on ice. Slides were washed in PBS + 0.1% Tween (PBST) followed by an endogenous peroxidase block (Bloxall, Vector Labs, SP-6000-100). Slides were subsequently blocked in 2.5% normal horse serum and stained overnight in primary antibodies at 4°C in PBS + 0.01% Tween-20. The following day, slides were washed in PBST and stained with the secondary-HRP conjugated antibodies, washed in PBST, and developed with 3,3’-diaminobenzidine (DAB, Vector Labs, SK-4100). For mouse-IgG primary, a M.O.M. kit was used after the peroxidase block (Vector Labs, MP-2400). Antibodies used for IHC are listed in **Supplementary Table 10**. Slides were scanned on a Pannoramic Scanner (3DHistech) with a 20X/0.8NA objective and visualized in ImageJ or QuPath (v0.4.2).

#### Multiplexed immunofluorescence (Leica Bond RX)

Samples were pretreated with EDTA-based epitope retrieval ER2 solution (Leica, AR9640) for 20 minutes at 95°C. The 6-plex antibody staining and detection were conducted sequentially. Antibodies used for multiplexed IF are listed within **Supplementary Table 10**. After 1 hr incubation, Leica Bond Polymer anti-rabbit HRP was applied followed by Alexa Fluor tyramide conjugate 488 and 647 (Life Technologies, B40953, B40958), or CF® dye tyramide conjugate 430, 543, 594, and 750 (Biotium, 96053, 92172, 92174, 96052) for signal amplification-based detection. At each round, epitope retrieval was performed for denaturation of primary and secondary antibodies before the following primary antibody was applied. Slides were washed in PBS and incubated in 5 μg/ml 4’,6-diamidino-2-phenylindole (DAPI; Sigma Aldrich) in PBS for 5 min, rinsed in PBS, and mounted in Mowiol 4–88 (Calbiochem). Slides were scanned on a Pannoramic Scanner (3DHistech) with a 20X/0.8NA objective and visualized in ImageJ or QuPath. Confocal microscopy was performed on a Leica Stellaris 8.

#### Multiplexed immunofluorescence (Lunaphore COMET)

Tissue was cut at 4 mm onto positively charged glass slides. Slides were baked at 64°C for 1 hr. Dewaxing and antigen retrieval was performed on the Leica Bond RX with 30 min retrieval EDTA-based epitope retrieval ER2 solution. Before loading onto the COMET, slides were washed 3X for 1 min in DI water. 20-plex antibody panel and dilutions can be found in **Supplementary Table 5**.

#### Immunoblotting

Single cell organoid suspensions or monolayer cells were lysed in 125-250 μL ice-cold RIPA buffer (Pierce, #89900) supplemented with 1x Complete Mini inhibitor mixture (Roche, #11836153001) and mixed on a rotator at 4°C for 30 minutes. Protein concentration was quantified using the Bio-Rad DC Protein Assay (Catalog #500-0114). 10–20 μg of total protein was separated on 4–12% Bis-Tris gradient gels (Bio-Rad) by SDS-PAGE and then transferred to nitrocellulose membranes. Antibodies used for western blots are listed in **Supplementary Table 10**. Blots were developed in Amersham ECL western detection region (Cytiva, RPN2236) and imaged on a Cytiva Amersham ImageQuant 800.

#### Lentiviral production

Lentiviruses were produced by co-transfection of 293T cells (Takara, #632180) with lentiviral backbone constructs and packaging vectors (psPAX2 and pMD2.G; Addgene #12260 and #12259) using TransIT-LT1 (Mirus Bio #MR 2306). Virus was concentrated through ultracentrifugation (Beckman, Optima L-100 XP) at 25,000 RPM for 2hrs. The viral pellet was resuspended in OptiMEM (Thermo, 31985062), aliquoted, frozen at −80°C, and titered through serial dilution assays.

#### Organoid culture

Murine prostate organoids were established and maintained as previously described^18,19^. Full prostate organoid media composed of: 1X ADMEM, 10 mM HEPES, 1X Glutamax, 0.5X Pen/Strep, 1X B27 (Fisher Scientific, #17504-044), 1.25 mM N-acetylcysteine (Sigma, A9165-100G), 10 mM Nicotinamide (Sigma, N0636-500G), 500 nM A83-01 (Tocris, #2939), 5 ng/mL recombinant murine EGF (Peprotech, #315-09), 10 ng/mL recombinant NRG1 (Peprotech, #100-03), 1 nM Dihydrotestosterone (Selleck Chemicals, S4757) 10% NOGGIN conditioned media, 5% RSPO-I conditioned media. Single cell embedded prostate organoids were supplemented with 10 μM Y-27632 (Fisher Scientific, #50-863-7) for the first 2-3 days in culture prior to change with fresh organoid media lacking Y-27632. *Ex-vivo* transformed organoids were seeded at 3E3 cells per 20 μL matrigel dome and passaged every 3-4 days. Wild-type organoids were seeded at 10E3 cells per 20 μL matrigel dome and passaged every 5-7 days. For monolayer adaptation, western blot validated knockout organoids were dissociated using methods described above and seeded at 1E5 cells/mL in full organoid media containing 10 μM Y-27632 on a collagen I coated 10 cm plate (Fisher Scientific, #08-772-75). After 5 days adaptation and expansion, cells were processed for protein lysates, or dissociated for orthotopic transplantation.

#### Organoid Cas9 ribonucleoprotein electroporation

Organoids were electroporated as previously described^20^. Briefly, organoids were electroporated with Cas9-RNP complexes (IDT) and recovered in organoid conditions for 3-5 days. *Trp53* loss was selected by supplementing media with 5 μM Nutlin-3a (Tocris, 6075). *Rb1* loss was selected for by supplementing media with 2.5 μM Palbociclib (Tocris, 4786). *Pten* loss can be optionally selected for by growing EGF and NRG1 knockout organoid media. Sequences for sgRNAs can be found in **Supplementary Table 11**.

#### Organoid lentiviral infection

RNP-edited and selected single cell organoid suspensions were infected with concentrated lentiviral supernatants at predetermined titers as previously described^18^. 3-5 days post lentiviral infection, organoids were dissociated with TrypLE (see above) and resuspended in 0.1% BSA in PBS + 1 mM EDTA and supplemented with 10 μM Y-27632. Single cell suspensions were passed through a sterile 5 mL polypropylene tube with cell strainer (Corning, #352235) and enriched by sorting for DAPI-EGFP+ cells using a Sony MA900 (Sony Biotechnology) with a 130-μm sorting chip (Sony Biotechnology, #LE-C3213). Sorted organoids were expanded for 5-6 days before transplantation.

#### Molecular Cloning

Lentiviral vector (LVt-UBC-cMYC-P2A-EGFP) was generated using Gibson assembly. Briefly, PCR fragments were amplified containing UBC promoter, *cMyc^T58A^* codon optimized cDNA, and a P2A-EGFP sequence, mixed within a Gibson master mix reaction and transformed into Stbl3 chemically competent *E. coli* (Thermo, #C737303). All plasmids were purified (Qiagen, #12943) and sequence validated through long-read sequencing (SNPsaurus). Lentiviral construct UT4GEPIR (Addgene #186712) was used for cloning Dox-inducible *Ascl1* and *mScarlet* constructs. Briefly, UT4GEPIR was digested with BamHI and I-SceI and a geneblock (IDT) encoding mouse codon-optimized *Ascl1-T2A-mScarlet* or *mScarlet* sequences with compatible overhangs was cloned by ligation and transformation as above.

#### Isolation and validation of *Ascl1* knockout organoid clones

*Ascl1* sgRNA targeted RPM organoids with Cas9 RNP were expanded for 5 days post electroporation and gently triturated in 0.5% BSA in PBS. Intact spheres were subsequently diluted 1:10 in PBS and placed in a 6 well plate (Fisher Scientific, #07-000-646). Using a standard tissue culture microscope and a 20 uL pipet tip, individual intact spheres with a healthy morphology were isolated and individually dissociated in 100 μL TrypLE and quenched in 1 mL of 0.5% BSA in PBS. Organoid clones were centrifuged at 600g for 3 min in protein low-bind microcentrifuge tubes and resuspended in 50 μL of matrigel and plated into 24-well suspension plates. Full organoid media supplemented with 10 μM Y-27632 was added after 10 min. Individual organoid clones were expanded in parallel and genomic DNA was isolated using a DNeasy Blood & Tissue kit (Qiagen, 69506). 35 cycle PCR reactions with an input of 100 ng of gDNA and an annealing temperature of 58°C were performed using primers flanking the sg*Ascl1* edit site and KAPA mouse genotyping kit following manufacturer protocols (Fisher Scientific, #50-196-5243; **Supplementary Table 11**). A PCR product of 170 bp was purified using the QIAquick Gel Extraction Kit (Qiagen, 28706) and submitted for library preparation and next generation sequencing at the Integrated Genomics Operation (IGO) at MSKCC. A total of 6 sequence-validated bi-allelic RPM-*Ascl1* KO clones were subsequently pooled and expanded for transplantation experiments.

#### CRISPR sequencing

Sequencing libraries were prepared from amplicons with an average size of 200-280bp (see **Supplementary Table 11** for PCR primer sequences). The reported concentration of 500 ng was used as input for the KAPA Hyper Library Preparation Kit (Kapa Biosystems KK8504) according to the manufacturer’s instructions with 8 cycles of PCR. Barcoded libraries were pooled at equal volumes and run on NovaSeq 6000 in a PE150 run, using the NovaSeq 6000 S4 Reagent Kit (300 Cycles) (Illumina). The average number of read pairs per sample was 1.3M. Alignment and modification quantification was done with CRISPResso2 (http://crispresso.pinellolab.org/) using default parameters.

#### RNA isolation from organoid cultures and bulk tumors

Tumors were isolated from euthanized mice and validated for EGFP fluorescence under fluorescence stereomicroscope. Tumors were quickly placed within 250-500 μL of RLT buffer supplemented with B-mercaptoethanol into 2mL tube with ceramic beads (MP, 6910500). Tumor samples were lysed on a Fisher Bead Mill 24 using manufacturer recommended settings. Lysates were passed through a Qiashredder (Qiagen, 79656) and RNA isolated using the Qiagen RNeasy kit (Qiagen, 74106) with manufacturer recommended protocols. Organoids were dissociated as above and resuspended in 300 μL of RLT buffer supplemented with B-mercaptoethanol and spun through a Qiashredder.

Qiagen RNeasy kit was performed to isolate RNA using manufacturer recommended protocols. For qPCR, purified RNA was reverse transcribed (Thermo, 4368814) and quantified (Applied Biosystems, Quantstudio 6) with SYBR green reagent (Thermo, A46110). See **Supplementary Table 11** for primer sequences used for qPCR.

#### Bulk Transcriptome sequencing and gene set enrichment analysis

After RiboGreen quantification and quality control by Agilent BioAnalyzer, 500 ng of total RNA with RIN values of 8.3-10 underwent polyA selection and TruSeq library preparation according to instructions provided by Illumina (TruSeq Stranded mRNA LT Kit, catalog # RS-122-2102), with 8 cycles of PCR. Samples were barcoded and run on a NovaSeq 6000 in a PE100 run, using the NovaSeq 6000 SX Reagent Kit (Illumina). An average of 24 million paired reads was generated per sample. Ribosomal reads represented 0.4-1.5% of the total reads generated and the percent of mRNA bases averaged 86%. Analysis of bulk RNA sequencing was performed at NYULMC HPC UltraViolet (formerly BigPurple) cluster using software provided in the Seq-N-Slide pipeline (https://github.com/igordot/sns), through the *rna-star* followed by *rna-star-groups-dge* routes. Briefly, after quality control assessment with MultiQC^58^ (python/cpu/v2.7.15) and sequencing adaptor trimming with Trimmomatic^59^ (v0.36), reads were aligned to the mm10/GRCm38 mouse reference genome with a splice-aware^60^ (STAR v2.7.3a) alignment, followed by featureCounts^61^ (subread/v1.6.3) to generate the RNA counts table. Counts were normalized for gene length and sequencing depth and tested for differential mRNA expression between groups using negative binomial generalized linear models implemented by the DESeq2^62^ 1.40.1 R package (r/v4.1.2). Differential expression was assessed by principal component analysis (prcomp function from the stats v4.3.1 R package) or unsupervised hierarchical clustering (pheatmap v1.0.12 and ComplexHeatmap v2.16.0) and visualized by a volcano plot (EnhancedVolcano v1.18.0) and TPM expression heatmap illustrating genes of interest. Differentially expressed genes (DEGs) identified were further analyzed for gene set enrichment analysis (GSEA) and pathway analysis with R packages: fgsea v1.26.0 and msigdbr v7.5.1. Moreover, DEGs were analyzed for enrichment of previously curated neuroendocrine signatures^13^. For this, we used the java GSEA Desktop Application (v4.3.2) with the GSEA Preranked module using the variance-stabilized log fold changes as metric to rank the DEG list in a descending order.

#### snRNA sequencing

Briefly, a single 10-week RPM tumor was extracted from the mouse prostate, and EGFP signal assisted in tumor microdissection (Nikon, SMZ18). Tumor sample was sliced into ∼5 mm x 5 mm pieces, dabbed on a kimwipe to remove moisture, and flash frozen in liquid nitrogen. Tumor piece was loaded onto a Singulator 100 (S2 Genomics) cartridge supplemented with 3.5 μL of 1 M DTT (Sigma, 43816-10mL) and 87.5 units of Protector RNAse inhibitor (Sigma, 3335402001). Nuclei were isolated with standard-nuclei isolation protocol according to manufacturer recommendations. Nuclei suspension was cleaned with sucrose density gradient (Sigma, NUC201-1KT) in protein low-bind microcentrifuge tubes (Eppendorf, 0030108442) at 500g for 5 min. Nuclei were subsequently resuspended in nuclei wash buffer: 10 mM Tris-HCl pH 7.4, 10 mM NaCl, 3 mM MgCl_2_, 1 mM DTT, 1% BSA, 1 U/μL Protector RNAse inhibitor, strained through blue-capped 35 μm FACS tube, and flow sorted for 7-AAD+ population to obtain single nuclei suspension. Sorted nuclei were submitted to the Single Cell Analytics Innovation Lab (SAIL) at MSKCC. Nuclei were validated for integrity under phase-contrast microscopy, and processed on Chromium instrument (10X Genomics) following user guide manual for 3’ v3.1 snRNAseq. Nuclei were captured in droplets, emulsions were broken, and cDNA purified using Dynabeads (Thermo, 37012D) followed by PCR amplification per manual instruction. ∼10,000 cells were targeted for each sample. Final libraries were sequenced on Illumina NovaSeq S4 platform (R1 - 28 cycles, i7 - 8 cycles, R2 - 90 cycles) at the Integrated Genomics Operation (MSKCC). FASTQ files were processed using the 10X Cell Ranger 6.1.2. Cell Ranger count was used to align reads to the GRCm38 (mm10) reference genome for snRNAseq samples and a modified version of GRCm38 (mm10) that includes the *Myc-P2A-EGFP* transgene sequences, given that organoids were infected with lentiviruses harboring its expression (**Extended Data** Fig. 1b). We set “include introns=TRUE’’ to accommodate the higher rate of intronic reads in snRNA. Finally, Cell Ranger count was used to generate feature-barcode matrices for subsequent bioinformatic analyses.

#### Spatial transcriptomics by 10X Genomics Visium

We generated Visium data from two adjacent sections as technical replicates, with each slide containing 10-week RPM tumor tissues from two individual mice. Visium Spatial Gene Expression slides were prepared with FFPE sections by the Molecular Cytology Core at MSKCC. Tumor samples with a target RIN value > 0.5 were processed for spatial transcriptomics. Probe pairs targeting the whole transcriptome were added to slide capture areas and allowed to hybridize overnight at 50°C. Bound pairs were ligated to one another, then released from the tissue by RNase treatment and permeabilization and captured by oligos bound to the slide. Probe extension and library preparation proceeded using the Visium Spatial for FFPE Gene Expression Kit, Mouse Transcriptome (10X Genomics, 1000337) according to the manufacturer’s protocol. After evaluation by real-time PCR, sequencing libraries were prepared with 11-14 cycles of PCR. Indexed libraries were pooled equimolar and sequenced on a NovaSeq 6000 in a PE28/88 run using the NovaSeq 6000 SP Reagent Kit (100 cycles) (Illumina). FASTQ files from sequencing (NovaSeq) were processed via spaceranger count (version 2.0.0) to align reads to the GRCm38 (mm10) reference genome and generate count matrices for subsequent bioinformatic analyses. An average of 74 million paired reads was generated per sample, corresponding to 37,000 reads per spot.

### Pathology annotation and spatial immunofluorescence analysis

Sections processed for H&E and multiplexed IF were reviewed by a board-certified genitourinary pathologist (A.G.). Graded histological areas were used to identify regions of PRAD and NEPC on serially sectioned samples processed for 10X Visium and multiplexed IF.

#### Cell segmentation

We utilized Mesmer^63^, a deep-learning algorithm designed for cell segmentation, to identify cell boundaries in COMET images. The input to Mesmer is a single nuclear image and single membrane or cytoplasm image. We used DAPI as a nuclear marker. To demarcate various cell types, we merged images from multiple cell-type-specific markers of membranes or cytoplasm by applying min-max normalization to each channel before summing them. The min-max scaling was performed using the *MinMaxScaler* function in *sklearn.preprocessing* package using default parameters. For COMET, we combined CD45, CD20, CD4, CD8, CD11b, CD11c, Ly6G (immune cells), CD31 (endothelial cell), GFP, KRT8 (tumor epithelial cells), VIM, a-SMA (stromal cells). The markers were chosen because they cover the cell types of our interests. We ran Mesmer on these images with standard parameters to predict cell boundaries (modified slightly to exclude cells smaller than 36 pixels), and calculated the cell size, eccentricity, and centroid of each cell boundary. We preprocessed COMET images to half their size (to 0.46µm per pixel) to accommodate system memory constraints (128 GB).

#### Normalization

Initially, we quantified raw per-cell marker expressions by aggregating pixel brightness within each cell boundary. To neutralize cell size variance in our analysis, expressions per channel were normalized against cell boundary area. We found bimodal distributions of cell size and DAPI expression in our dataset. The lower mode of DAPI contained primarily empty regions and the upper mode of cell size contained cell segmentation errors. We then filtered all cells with DAPI value less than 2096 (estimated from the distribution) and cell size larger than 2500 (estimated from the distribution). The marker intensity signals then underwent logarithmic transformation.

#### Cell type identification

For tumor cell identification within our dataset, we used K-nearest neighbor analysis (k=30) and clustered normalized marker expressions using the Leiden algorithm^64^, yielding 27 distinct clusters. These clusters were labeled as stromal, tumor (notably marked by GFP, ASCL1, KRT8), or artifacts (characterized by low marker expression), with artifact cells excluded from further analysis. Cell types were then classified based on marker intensity distributions: lymphatic endothelial cells (LYVE1>7.5), blood vessel endothelial (CD31>8), and immune (CD45>8). The distribution patterns of this coarse classification is reported in Figure 4a. With the same approach, we identified sub population in immune cell one by one: CD4 T cell (CD4>7), B cell (CD20>8), CD8 T cell (CD8>8.3), Treg (FoxP3>8), CD11b F4/80+ myeloid (Mac3; CD11b>7 & F4/80>7), CD11b F4/80-myeloid (Mac1; CD11b>7 & F4/80<7), CD11c F4/80+ myeloid (Mac2; CD11c>8.5 & F4/80>7).

#### Spatial analysis

To examine cellular organization within lymph node (LN) tissue, particularly dense with immune cells, we constructed a spatial neighborhood graph by linking cells within a 20-pixel radius, approximately 2.5 times the median cell radius.

This analysis indicated an average of 4.5 neighbors per cell. A neighborhood enrichment matrix was generated to map interactions between cell types, with axis labeling indicating source and neighboring cell types and matrix values representing total neighboring counts. Normalizing these counts by the total number of neighbors per cell type produced a cell proximity frequency matrix (illustrated in Figure 4g), showcasing the likelihood of each cell type neighboring another. The spatial neighborhood graph and the neighborhood enrichment matrix were implemented utilizing Squidpy^65^.

For infiltration analyses, we employed HALO and HALO A.I. module v.3.6.4134 (Indica Labs). HALO A.I. module was used to obtain nuclear and cytoplasmic segmentation masks from a set of training slides. Cell nuclei were segmented using the DAPI channel. The trained module was then applied to each experimental slide. Annotations of nuclear, cytoplasmic, and background staining were reviewed manually for each slide. Thresholding parameters for each marker were kept consistent across all slides. Single cells were filtered to exclude errors in cell segmentation through DAPI-based criteria, including: 1) nuclear DAPI measurements above a user-input threshold, and 2) the nuclear/cytoplasmic ratio DAPI measurement was above a user-input threshold. Cell types were classified based on a hierarchy for positive or negative stains. Each cell is evaluated based on predetermined marker co-expression and then assigned as cell types in a layered fashion (**Supplementary Table 6**). PRAD and NEPC regions were manually annotated as above. Exported data files were used for image analysis. For distance infiltration metrics, each individual primary tumor region was separated into 15 bins spanning a total of 1000 μm inside and outside the region of interest and within the confines of the tumor boundary. For smaller metastatic nodules in the lung and liver, 10 bins spanning a total of 500 μm inside and outside and within the confines of the tissue were used for cell infiltration analysis. All tumors within an individual metastatic tissue sample were pooled together for infiltration analysis. Cells residing within each bin were quantified and normalized to the bin area. Subsequently, the data mean cell density per bin was averaged across independent mouse samples. Biological replicate samples (*n* ≥ 3) were used to calculate standard error. Data visualized in R using the ggplot2 package. Loess function was implemented to visualize smoothened error curves across the defined tissue region. For histological characterization and quantification of tumor area, a Random Forest tissue classifier was created on the HALO platform and iteratively refined using a training set of *n*=10 primary and metastatic RPM tumors containing examples of NEPC, PRAD, and stroma. A single primary RPM tumor derived from a castrated host was used as training input for chondrosarcoma histology. All tissue classification was subsequently cross-referenced with G.U. pathologist, Dr. Gopalan.

### snRNA-seq analysis

#### Preprocessing

We first used CellBender^66^ to remove ambient RNA molecules from the raw count matrices with the parameter --expected-cells 5000 and --total-droplets-included 20000. These parameters were chosen by inspecting the barcode rank plot generated by Cell Ranger, following author recommendations. Defaults were used for all other parameters. All downstream processing of snRNA-seq data and scRNA-seq data, discussed in the section “Reclustering of the public scRNA-seq dataset”, were performed in Scanpy^67^. We next removed low quality cells, unexpressed genes, and potential doublets from the CellBender output. We filtered to exclude i) genes detected in fewer than 3 cells, ii) cells with under 200 genes or 1000 UMIs, and iii) cells with a mitochondrial fraction above 10%. Mitochondrial and protein-coding ribosomal protein coding genes were also omitted from downstream analyses, as these are often a large source of spurious variance that can dominate clustering and confound deconvolution^29^. We used Scrublet^68^ to remove doublets. After Scrublet, 4,872 cells were retained for downstream analyses, with a median of 7055.5 UMIs per cell. Raw counts were normalized by log_2_ (X+1) transformation, where X = library size of each cell / 10^4^.

#### Dimension reduction, clustering and cell type annotation

We selected the top 3000 highly variable genes in each dataset using the Scanpy function pp.highly_variable_genes with raw counts as input, n_top_genes = 3000, flavor = ‘seurat_v3’ and span = 1. We applied PCA to these genes using the scanpy.tl.pca function with the arpack solver. We selected 30 PCs, which explain 40% of variance. For clustering, we used Phenograph^69^ with k = 30 and clustering_algo = ‘leiden’, which generated 19 clusters. Cell typing was performed using marker genes as shown in **Supplementary** Fig. 4. To distinguish malignant cells from non-malignant cells, we performed copy number analysis inference using inferCNV^70^ (Trinity CTAT Project, https://github.com/broadinstitute/inferCNV). We used myeloid and endothelial cells determined based on marker expression as the normal cell reference. inferCNV showed a major CNV group of low CNV score containing two additional phenograph clusters, which were labeled as normal epithelial and mesenchymal cells. All other clusters show significant CNV changes, and were labeled as NE (neuroendocrine), EMT (epithelial-to-mesenchymal transition) and Tumor Luminal/Basal according to expression of marker genes, with each subtype containing 6, 4 and 5 cell states respectively.

#### Reclustering of the public scRNA-seq dataset

To support BayesPrism and PrismSpot, we used an independent scRNA-seq data from prostates derived from *Pten^fl/fl^*; *Rb1^fl/fl^*; *Trp53^fl/fl^*; *Probasin-Cre* (PtRP) GEMMs^13^. We chose to use this dataset as 1) to demonstrate generality, as the dataset was generated from an independent experiment using a mouse model that similarly transitions from an adenocarcinoma to neuroendocrine prostate cancer, 2) it contains increased cell type diversity due to the mechanical and enzymatic digestion that skews towards non-tumor cells (total cells *n* = 67,622). To improve resolution for our BayesPrism deconvolution and PrismSpot analysis, we re-clustered the data to improve the granularity of mesenchymal cells. Specifically, as mesenchymal cells were not the focus of the original paper, authors only clustered it into two populations: endothelium and mesenchymal.

However, given the observed heterogeneity in this population, we subsetted GFP-negative mesenchymal cells from this dataset, and reclustered this population. We selected the top 3000 highly variable genes in each dataset using the Scanpy function pp.highly_variable_genes with raw counts as input, n_top_genes = 3000, flavor = ‘seurat_v3’ and span = 1. Raw counts were normalized by log_2_ (X+1) transformation, where X = library size of each cell / 10^4^. We applied PCA to these genes using the scanpy.tl.pca function with the arpack solver and selected 30 PCs. For clustering, we used Phenograph with k = 30 and clustering_algo = ‘leiden’, which generated 20 clusters. Clusters were annotated by expression of marker genes curated from Niec et al.^30^, guided by hierarchical clustering over the PC space. Endothelial, lymphatic, glial cell, pericytes/myofibroblast were clearly distinguishable based on the marker genes (**Supplementary** Fig. 6a). The rest of the clusters fall into two major groups as shown by hierarchical clustering and were annotated as Mes-1 and Mes-2 (Mesenchymal, **Supplementary** Fig. 6a and b). The original mesenchymal cell type label was replaced by this higher granular cell type annotation (**Supplementary** Fig. 6c).

### Analysis of Visium dataset

#### Cell type deconvolution using BayesPrism

We deconvolved Visium data from two replicates using the snRNA-seq data collected from the 10-week RPM tumor. We used BayesPrism to perform statistical deconvolution as it has been previously shown to yield accurate results for Visium data and outperformed other methods in multiple settings^30,71–75^. More importantly, BayesPrism jointly imputes the posterior of cell type fraction and the cell type-specific gene expression profile in each Visium spot, enabling the cell type-specific gene count matrices by PrismSpot, as described below.

To increase signal to noise ratio for deconvolution, we selected marker genes that are differentially upregulated in each of the 19 cell types, which was defined as the 19 phenograph clusters in the snRNA-seq data. We then took the union of these marker genes and deconvolved over these genes. Specifically, we performed a pairwise *t*-test using the findMarker function from the scran package^76^, provided in the wrapper function get.exp.stat from the BayesPrism package. As each cell type is compared against every other cell type, we need to summarize N-1 *p*-values and log_2_ fold change statistics for each cell type (N denotes number of total cell types). For each non-tumor cell types (4 in total), i.e., mesenchymal, myeloid, normal L/B (luminal/basal) and endothelial cells, we required both the maximum *p* value to be less than 0.01, and the minimum log_2_ fold change to be above 0.1, where the max and min were taken over every other cell type. For each tumor cell type (15 in total), i.e., NE, EMT and tumor L/B, we required the same threshold for *p*-value and log_2_ fold change, but the max and min were taken over every other non-tumor cell type. As a result, we avoided the comparison between tumor cell types when taking the max and min to retain the maximum number of tumor-specific genes for PrismSpot analysis. This was achieved by defining a coarse level cell type label, where 15 tumor cell types were grouped into a single tumor cell type and using it as the cell.type.labels argument, while setting the original 19 cell types as cell.state.labels in BayesPrism’s built-in function get.exp.stat. To speed up deconvolution we further excluded genes expressed in less than or equal to 3 spots. This yielded 5,125 genes in total used as the input for BayesPrism. We defined each of the 19 cell states as individual cell types when constructing reference for deconvolution. We set pseudo.min to 0, the numerical lower bound for genes with zero count in a given cell type, to maximize the contrast between different cell types in the reference. All other parameters of BayesPrism were used as default. We performed the updated Gibbs sampling for a more robust correction of batch effects between the snRNA-seq and the FFPE Visium.

#### Benchmarking of BayesPrism

We benchmarked both the accuracy and robustness of the cell type fraction deconvolved by BayesPrism. A big challenge in the deconvolution field is the lack of ground truth for benchmarking. To estimate some ground truth of the cell type fractions from the Visium data we used 1) histology with H&E, and 2) a marker genes-based approach. The comparison with histologically defined NEPC regions showed strong concordance with the fraction of NEPC cells inferred by BayesPrism (Fig. 5c). Additionally, we benchmarked the robustness of cell type fraction inferred by BayesPrism by assessing the concordance across 5 histologically defined regions between two technical replicates, which includes 3 neuroendocrine regions and 2 non-neuroendocrine regions. As tissues are slightly shifted between technical replicates, we manually adjusted the selected regions to achieve better spatial concordance on the x-y plane. For each region, we compute the average fractions for each cell type. We then computed both cell type-level and region-level Pearson’s correlation coefficients, which was shown in **Supplementary** Fig. 4a-b.

#### PrismSpot analysis

BayesPrism’s deconvolution infers cell type-specific gene expression matrices for each spot. For the Hotspot analysis, we use spot specific tumor gene expression as the input. Specifically, to generate tumor-specific gene expression profiles, we summed up the posterior mean of cell type-specific gene expression, *Z*, across all tumor clusters (*n* = 15) outputted by BayesPrism. As Hotspot models the raw count data using negative binomial distribution, we rounded up the posterior mean of *Z*. An important step in Hotspot analysis is to define neighborhood structure over which the spatial co-expression is defined. In our Visium data, we have two adjacent slides, referred to as technical replicates. Each technical replicate contains tumor samples from two different mice, referred to as biological replicates. To leverage the full set of Visium data points to yield higher statistical power while accurately reflecting the neighborhood structure, we performed a single Hotspot analysis for all Visium spots while restricting neighborhoods to be only within each mouse and each technical replicate. We achieved this by shifting the coordinates of Visium data points for each tumor sample in each replicate by a large numeric number, e.g.,1000, such that coordinates have no overlap within the Kth (K = 6, discussed below) nearest neighbor for Visium spots from different tumor samples. We excluded i) genes with zero count in all spots, and ii) spots containing fewer than 1000 genes or 1000 UMIs from downstream Hotspot analysis. We define n_neighbors=6 in the create_knn_graph in Hotspot to include only Visium spots that are adjacent to each other and disabled weighted_graph arguments to treat all adjacent spots and the self-spot equally. For the analysis in Figure 5, we selected genes with spatial autocorrelation FDR less than 0.01 and transcription factors within the top 500 autocorrelation *Z*-score, followed by re-computing the pairwise local correlation between the top 500 transcription factors. To cluster transcription factors into modules, we used the “create_modules” function provided by the Hotspot package, with parameters min_gene_threshold=15, core_only=True, fdr_threshold=0.05, which performs a bottom-up clustering procedure by iteratively merging two genes/modules with the highest pairwise *Z*-score.

To enhance the robustness of the identified gene modules against sampling noise, we implemented a subsampling strategy. Specifically, we subsampled 60% of the reads from the Visium data (tumor-specific gene expression deconvolved by BayesPrism) for each mouse tissue on each Visium slide using multinomial distribution. This process was repeated 100 times. Subsequently, we re-evaluated the clustering using the Hotspot method, which reclusters the genes into modules using the recalculated pairwise local correlations among genes. For each gene pair initially grouped together in a module using the full dataset, we assessed their co-occurrence in the same module across the subsampled datasets. This frequency of co-occurrence was defined as the consensus score. We then calculated the average consensus score for each gene with all other genes within the same module. A higher average consensus score indicates that a gene consistently remains within its module across different subsampling simulations, and hence stability within that cluster. Finally, we identified representative genes for each module by selecting those with an average consensus score ≥ 0.8 (Fig. 5e **and Supplementary Table 7**). In total, 71/181 genes from three modules were chosen as their representative genes.

#### Benchmarking of PrismSpot

We compared the Hotspot results between PrismSpot and “standard” Hotspot, i.e., only un-deconvolved raw count was used as input. We compared the autocorrelation and pairwise local correlation coefficients over marker genes of different cell types. We analyzed the Hotspot local correlation statistics over a more comprehensive gene set without pre-filtering genes based on autocorrelation to demonstrate the behavior of the statistics of different types of markers. We derived marker genes from the GEMM scRNA-seq dataset as mentioned above and grouped the cell types such that the granularity matches that of snRNA-seq reference. Specifically, we grouped Mes-1 and Mes-2 as stromal; *Tff3*, mutant L1, mutant L2, mutant B1, NEPC-*Pou2f3* and NEPC as tumor; macrophages, neutrophils, and DC as myeloid. We then performed differential expression using the pairwise *t*-test between tumor, stromal, myeloid, and endothelial cells, like the strategy described above. Genes with maximum *p* value less than 0.01, and the minimum log_2_ fold change above 0.1 were used as markers of each cell type for benchmarking PrismSpot.

We performed one-sided paired *t*-tests to compare auto-correlation *Z*-scores between PrismSpot and Hotspot on un-deconvolved raw counts, hereafter referred to as Hotspot. To ensure the statistical power was similar in the comparison of autocorrelation scores for each cell type, we selected the top 100 genes that pass the threshold mentioned above based on log_2_ fold change. For non-tumor cell types, i.e., endothelial, myeloid, and mesenchymal cells, we define null hypothesis H_0_: PrismSpot *Z* > Hotspot *Z*, while for tumor cells we define H_0_: PrismSpot *Z* < Hotspot *Z*. The *p*-values were 4.0×10^-3^, 2.1×10^-6^, 6.0×10^-9^, and 3.8×10^-3^ for endothelial, myeloid, mesenchymal and tumor cells, respectively (**Extended Data** Fig. 7c).

Likewise, we performed one-sided paired *t*-tests to compare local pairwise correlation *Z*-scores between PrismSpot and Hotspot across three categories, 1) between a pair of tumor marker genes, 2) between a tumor marker gene and a marker gene for non-tumor cell types, and 3) between a pair of marker gene for non-tumor cell types. As pairwise local correlation can be both positive and negative, we performed statistical tests on the absolute value of *Z*-scores. For tumor vs. non-tumor and non-tumor vs. non-tumor categories, we define null hypothesis H_0_: |PrismSpot| > |Hotspot|, while for tumor vs. tumor category, we define H_0_: |PrismSpot| < |Hotspot|. The *p*-value of tumor vs tumor category was 3.3×10^-5^. For tumor vs. non-tumor and non-tumor vs. non-tumor categories, *p* values were less than the numeric limit 2.2×10^-16^ (**Extended Data** Fig. 7g).

#### Analysis of human scRNAseq PRAD and NEPC myeloid subsets

FASTQ files from a previously published single cell dataset of 12 prostate cancer patients^13^ (histologically verified CRPC–PRAD (*n*=9) and NEPC (*n*=3)) was mapped to human reference genome GRCh38 using cellranger–7.0.1 to generate count matrices (transcripts/features x cells). Downstream analyses were performed using the Seurat R package (version 4.4.0). Cells were removed if features were not detected in at least 10 cells. Cells were filtered based on the following criteria: 1) less than 500 features; 2) ≥ 30% mitochondrial counts and ≤ 500 UMI counts. Putative doublets were removed using scDblFinder^77^. Combining samples from all CRPC-PRAD and NEPC sampled yielded 63,834 cells x 30,519 features. To normalize the data, the “LogNormalize” method was used with a pseudocount of 1 and a scale factor of 10,000. The top 2,000 highly variable genes were identified using the FindVariableFeatures() function. As patient tended to cluster by sample instead of by cell type, fastMNN was utilized across all cell types to perform batch correction (using all 30,519 features). Clustering was performed using FindNeighbors() and FindClusters() functions with a resolution of 0.3 on the batch–corrected count matrix. The resolution value was determined to be 0.3 as it best matched the expression patterns of all lineage markers. The clustered cells were visualized using RunUMAP() with the first 30 dimensions from the dimensional reduction ‘MNN’, and the clusters expressing myeloid lineage markers (*CD14*, *LYZ* and *IL1B*) were identified with these cells being subsetted for downstream analysis (N=7,004 myeloid cells). Re-clustering with a resolution of 1 was then conducted these putative myeloid cells using the batch-corrected matrix (from the upstream correction) yielded 17 clusters. Differentially expressed genes (DEGs) for each cluster were identified using FindMarkers() with MAST algorithm (version 1.24.1) and thresholds of Bonferroni adjusted *P* value < 0.05 and log_2_FC > 0.5. Of note, 4 clusters (N=1,282 cells) had low UMI counts, 1 cluster (N=382 cells) showed top DEGs possibly indicative of doublet cell types expressing markers for both epithelial cells and myeloid cells (*KLK3* and *CD14*) and 1 cluster (N=42 cells) expressed high levels of proliferation related genes (*MKI67*, *TOP2A* and *STMN1*) and therefore were removed. This yielded a total of 5,298 myeloid cells (4,348 CRPC–PRAD and 950 NEPC cells). To identify the subtypes of tumor– associated macrophages (TAMs), module scores from pre-defined gene sets for each TAM^78^ were used and scores were calculated using AddModuleScore(). Cells were labeled based on the median and maximum signature scores per cluster.

#### Analysis of human prostate SU2C dataset

The FPKM-normalized RNA-seq from Abida et al., was downloaded from https://github.com/cBioPortal/datahub/tree/master/public/prad_su2c_2019 (ref. 45). We selected patient samples sequenced by the poly-A enrichment protocol, as it contains more samples with histologically verified NEPC. In total there were 9 patients with neuroendocrine features, and 50 patients with non-neuroendocrine features. To compute the log_2_ fold change between neuroendocrine and non-neuroendocrine samples, we computed the log_2_ ((mean expression of neuroendocrine samples +1) / mean expression of non-neuroendocrine samples +1)). The statistical significance was computed using a two-sided Wilcoxon test.

#### Statistics and reproducibility

We used GraphPad Prism software v.9.5.1 for statistical analyses or in-house scripts in R v.4.3.1 which are available from the corresponding author upon reasonable request. Variance was similar between compared groups and *p*-values were determined by two-tailed Student’s *t*-test for all measurements comparing untreated to treated samples of single time points. One-way analysis of variance (ANOVA) with Sidak’s or Tukey’s multiple comparisons correction listed in the figure legends for comparisons across more than two groups. For analysis between groups over multiple time measurements (growth curves), two-way ANOVA was used with appropriate multiple comparisons testing listed in the figure legends. Figure legends denominate statistical analysis used. No statistical method was used to predetermine the required sample size. No data were excluded from this study. Investigators were not blinded to allocation during experiments and outcome assessment, except for mouse specific study analyses.

