## Extended Data Figures 1-10 for "The neuroendocrine transition in prostate cancer is dynamic and dependent on ASCL1"

### EXTENDED DATA FIGURE 1:

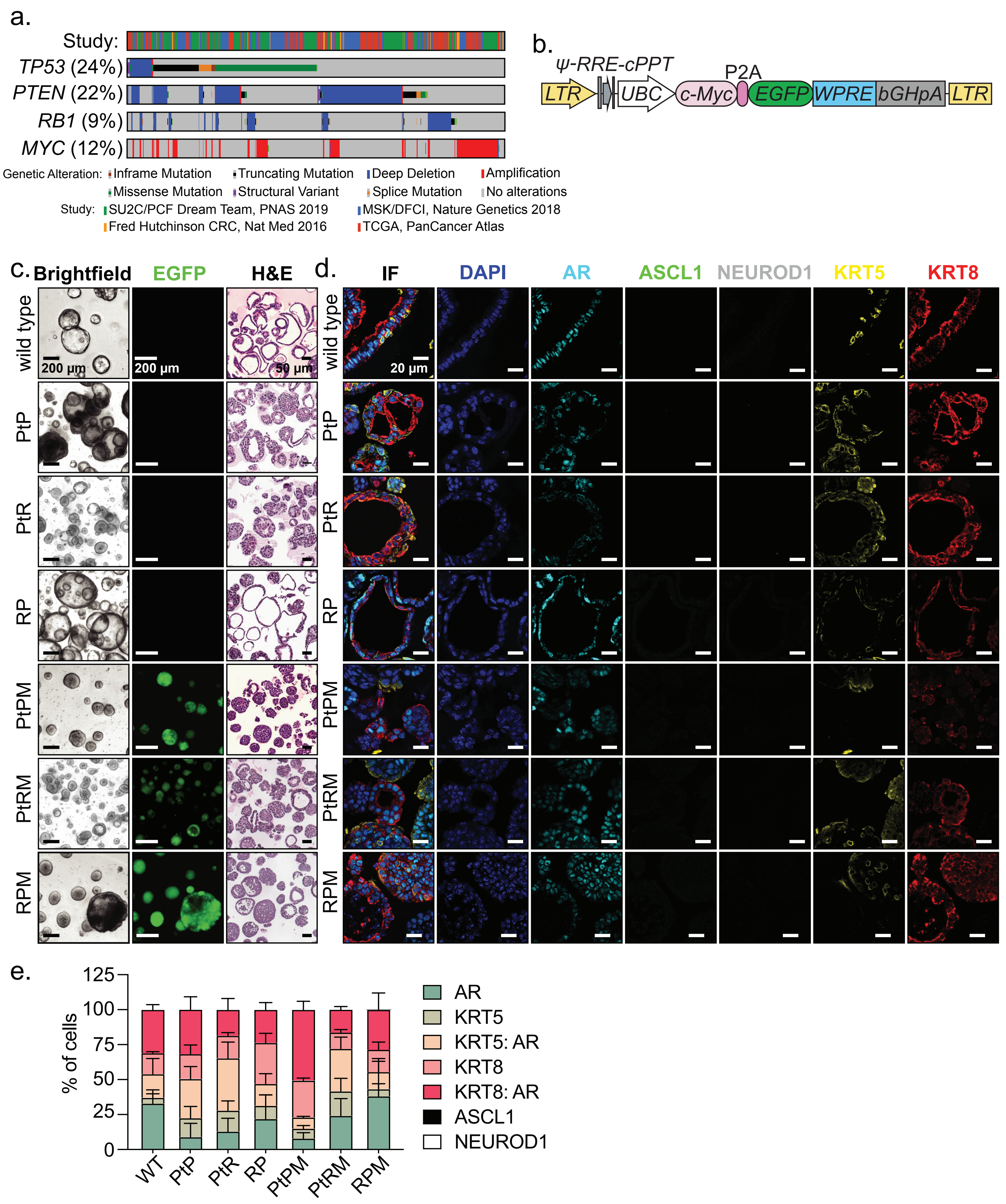

**Extended Data Figure 1:**

**a.** Oncoprint of the indicated genes frequently mutated in human primary prostate cancer. Samples obtained from the studies indicated within the figure legend. **b.** Schematic of the lentiviral vector used within this study to overexpress *cMyc*$^{T58A}$ transcriptionally linked to EGFP in organoids. **c.** Representative brightfield (left), GFP fluorescence (center), and hematoxylin & eosin (H&E) stains (right) of organoids harboring mutations in indicated tumor suppressors and oncogenes. **d.** Representative confocal images of 7-plex IF stains of organoids of the indicated proteins. Data are representative of *n*=3 technical replicates per genotype. **e.** Percentage of unique cell types expressing the indicated markers in organoid culture. Data representative of *n*=3 technical replicates and related to Extended Data Fig. 1d. Error bars denote mean and standard deviation. All scale bars and pseudocolor legend indicated within the figure panel.

EXTENDED DATA FIGURE 2:

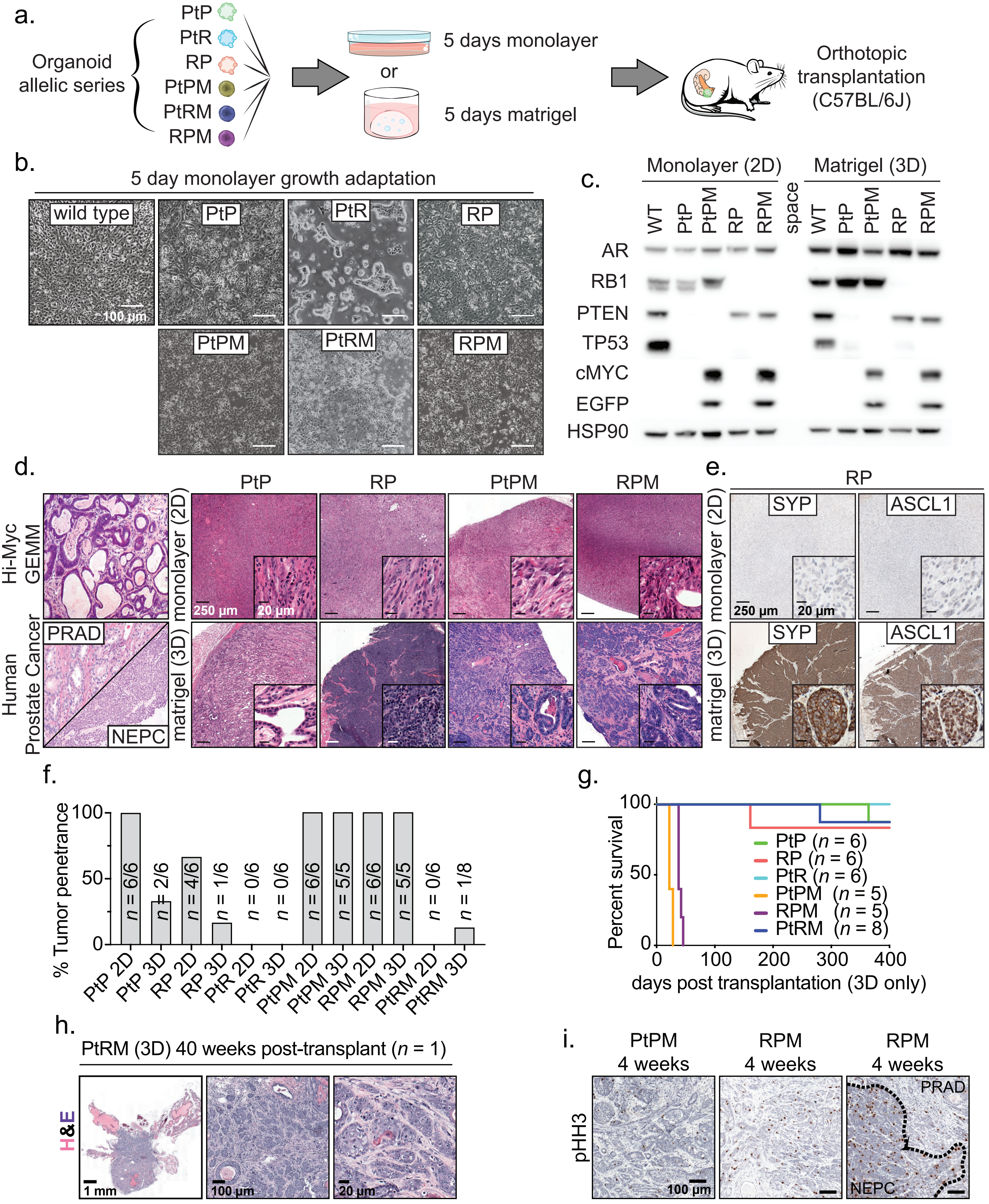

**Extended Data Figure 2:**

**a.** Schematic representation of steps taken to establish orthotopic prostate tumors in mice from edited organoids grown in matrigel (3D) or monolayer culture (2D) conditions. **b.** Representative brightfield images of the indicated organoids seeded in monolayer growth. Images taken 5 days post seeding. **c.** Western blot validation of knock out efficiency 5 days post electroporation with Cas9 in complex with purified sgRNA. As in a), edited organoids were seeded in matrigel or monolayer culture prior to lysis and western validation. Data representative of 2 independent experiments. **d.** Representative hematoxylin and eosin stains (low and high magnification images) of established mouse models (Hi-MYC), human prostate cancer, and organoid orthotopic transplant-derived prostate tumors. Data related to Extended Data Fig. 2a-c. OT prostate tumors derived from transplantation of organoids grown in (top) monolayer or (bottom) traditional 3D matrigel conditions. **e.** Representative Synaptophysin (SYP) or ASCL1 immunohistochemical stains of tumors isolated from mice transplanted with RP organoids grown in (top) monolayer or (bottom) traditional 3D matrigel conditions. Data representative of *n*=2 independent tumors per stain. **f.** Percentage of mice with tumors per genotype and organoid growth conditions. Sample size per genotype indicated within the figure panel and represent independently arising prostate tumors. **g.** Survival of mice OT transplanted with 250k dissociated organoids grown only in matrigel. Sample size per cohort indicated within the figure legend and are representative of independently arising tumors. **h.** Representative hematoxylin and eosin (H&E) stains of a single mouse that developed orthotopic tumors following transplantation of PtRM organoids grown in matrigel. **i.** Representative phospho-histone H3 immunohistochemical stains of PtPM or RPM orthotopic prostate tumors. Histological classification performed using serial sectioned H&E. Dotted line represents the boundary of PRAD and NEPC. Data related to Fig. 1d. All scale bars denoted within the figure panel.

### EXTENDED DATA FIGURE 3:

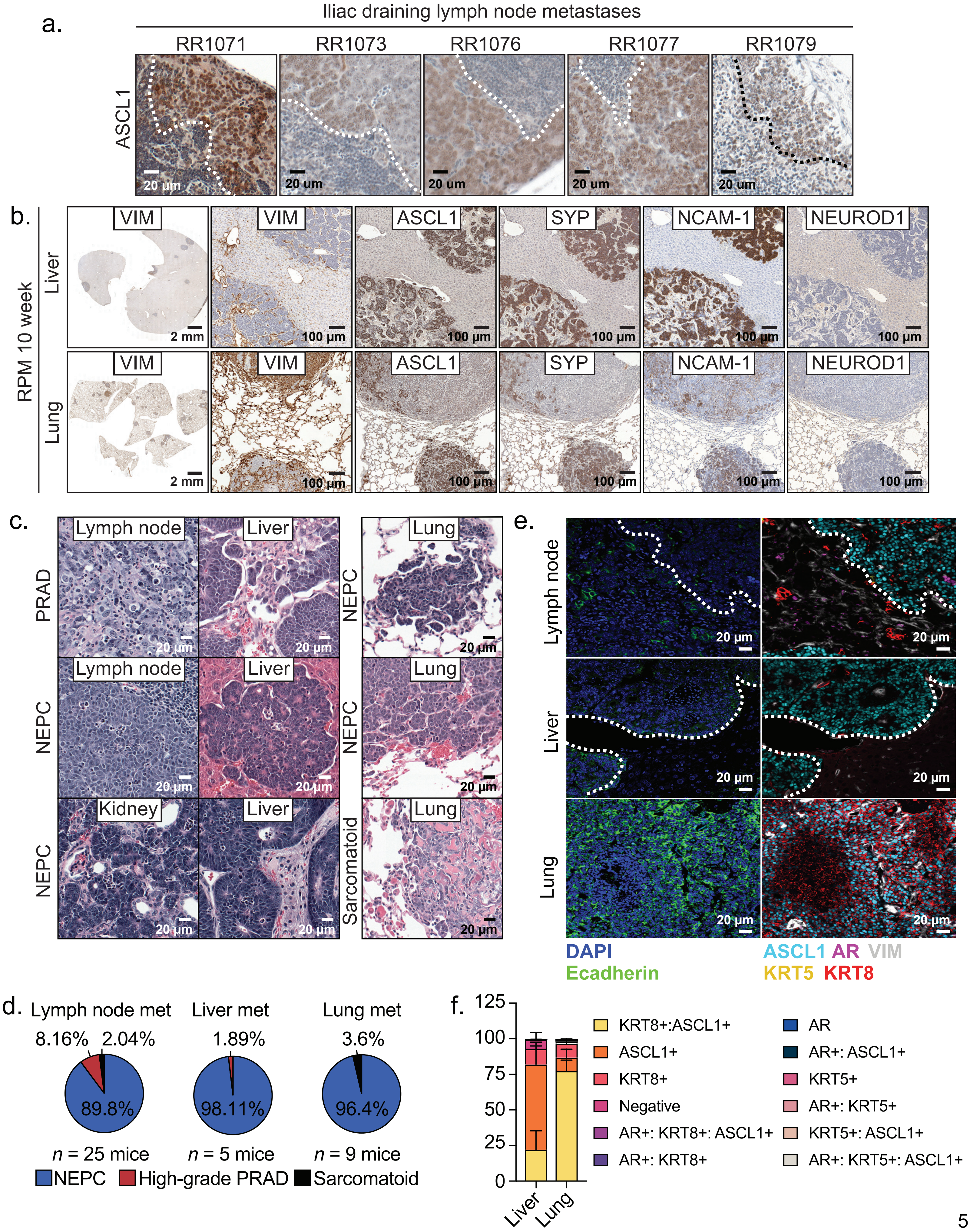

**Extended Data Figure 3:**

**a.** Representative ASCL1 immunohistochemical stains from iliac lymph node metastases isolated from 5 independent RPM OT transplanted mice. Dotted line represents the boundary of tumor and normal tissue. **b.** Representative serially sectioned immunohistochemical stains from metastases isolated across (top) liver and (bottom) lung tissue in RPM OT transplanted mice. Scale bars denoted in figure legend. Data representative of *n*≤5 independent mice. VIM = VIMENTIN, SYP = SYNAPTOPHYSIN. **c.** Representative H&E stains and histological grade of metastases isolated from regional lymph nodes, kidney, liver, and lung tissue from RPM OT transplanted mice. Data related to Extended Data Fig. 3a-b). Scale bars denoted within the figure panel. Data representative of *n*≤5 independent mice unless otherwise noted in the figure panel. **d.** Pie charts demonstrating percentage of mice harboring distinct histotypes of prostate cancer in the indicated metastatic regions. Sample size denoted in figure panel. **e.** Representative multiplexed IF staining for lineage markers in metastastic tumors isolated from regional lymph nodes, liver, and lung tissue from RPM OT transplanted mice. Data representative of *n*=3 independent mice. **f.** Percentage of unique cell types expressing the indicated lineage markers in RPM metastatic samples. Data representative of *n*=3 independent mice and related to Extended Data Fig. 3e. Error bars denote mean and standard deviation. All scale bars denoted in the figure panel.

### EXTENDED DATA FIGURE 4:

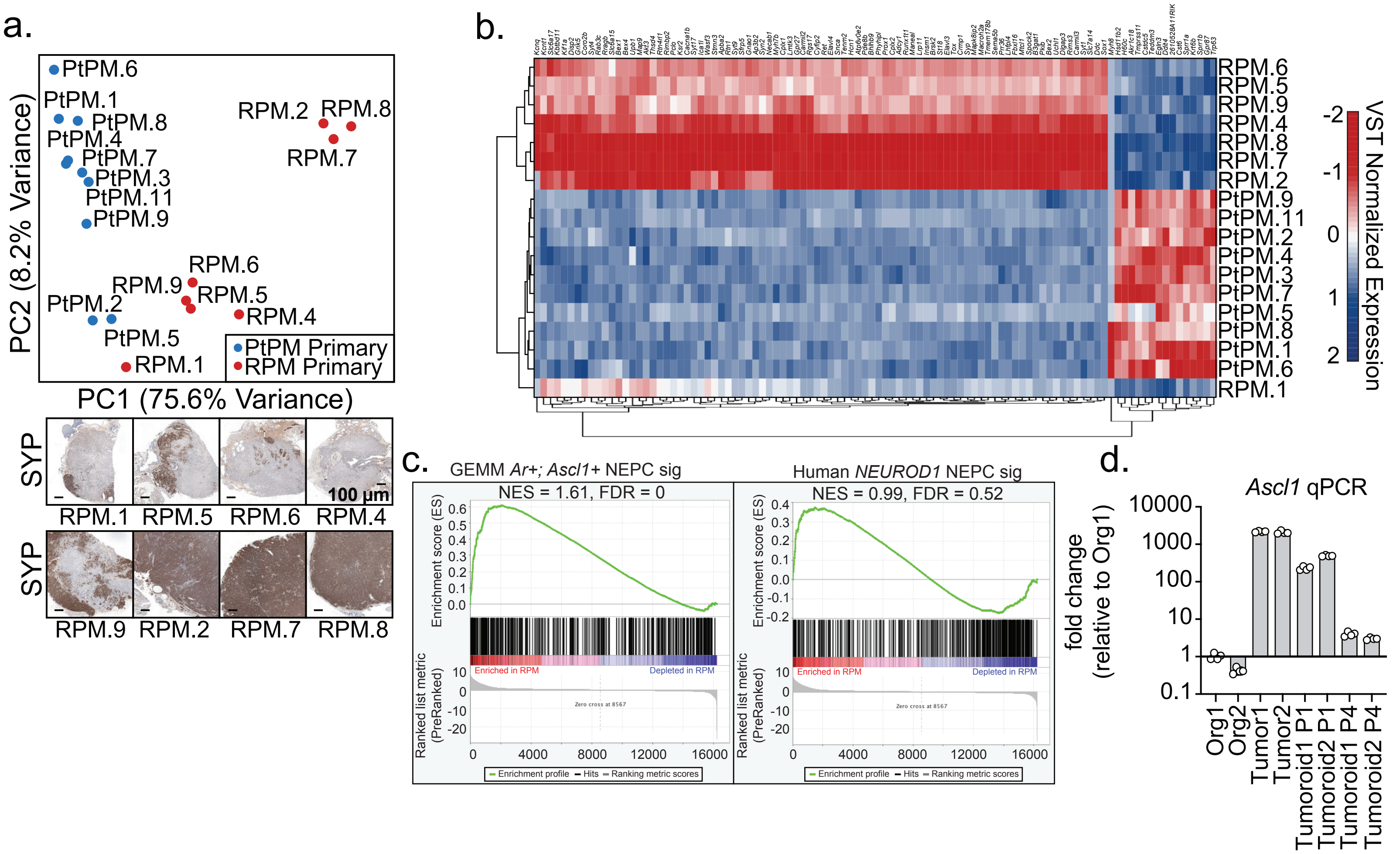

**Extended Data Figure 4:**

**a.** (Top) Principal component analysis of bulk RNA sequencing data isolated from RPM or PtPM OT transplants. (Bottom) Representative SYP immunohistochemical stains from the RPM tumors ordered by increasing percentage of SYP+ cells/tumor. RPM, *n*=8 independent tumors. PtPM, *n*=10 independent tumors. Data related figure Fig. 2f. **b.** Unsupervised hierarchical clustering of variant stabilized transcript normalized expression of the top 100 differentially expressed genes (columns) within tumors (rows). Data related figure Fig. 2c. **c.** GSEA enrichment plots of established expression signatures of (top) genetically engineered mouse model (GEMM) of *AR* and *Ascl1*-co-expressing NEPC harboring conditional deletion of *Pten*, *Rb1*, and *Trp53* (PtRP), and (right) histologically verified human NEPC expressing *NEUROD1* within RPM primary tumors. FDR and NES indicated in the figure. Analysis derived from the transcriptional profiles of multiple independent RPM tumors (*n* 8) relative to PtPM tumors (*n*=10). Data related to samples used in Fig. 2c-d. **d.** Quantitative PCR of *Ascl1* transcripts across two biologically independent RPM organoids (org), tumors, and tumor-derived organoids (tumoroids) at passage 1 (P1) or passage 4 (P4) post isolation. Each data point indicates technical quadruplicate values and bars represent mean and standard deviation. Data representative of 2 independent experiments.

EXTENDED DATA FIGURE 5:

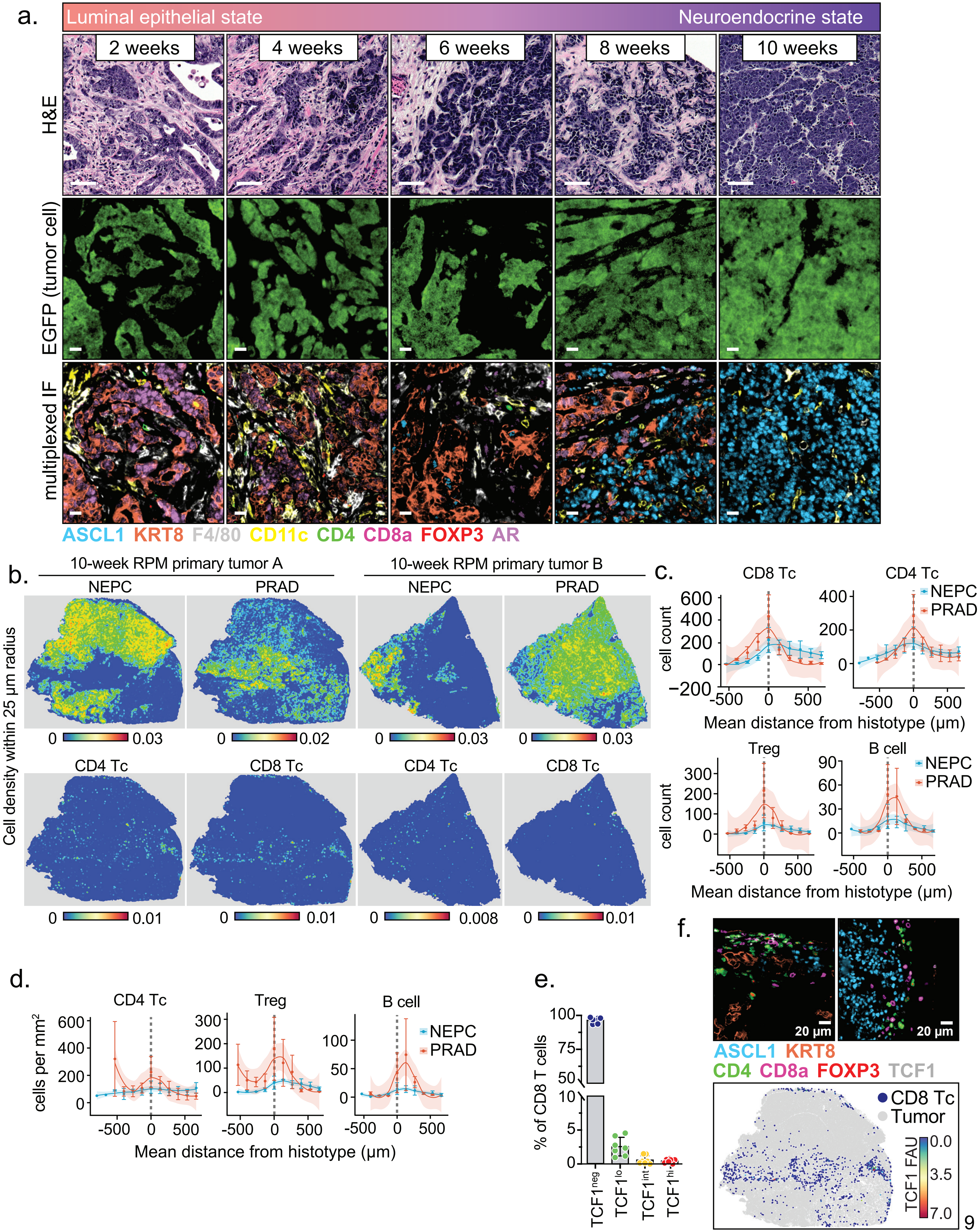

**Extended Data Figure 5:**

**a.** Representative (top) H&E, (middle) EGFP, and multicolor immunofluorescence (bottom) of RPM tumors isolated at the indicated time points. EGFP and Immunofluorescence images are matched sections. Data representative of *n*>3 tumors stained by COMET. Scale bar for all images indicates 20 µm. Pseudo-coloring listed within the figure panel. **b.** Spatial cell type density (cell density within 25 µm radius) for the indicated tumor cell types and lymphocytes across two independent 10-week RPM tumors. Heatmap represents average cell density (cell number/$\mu m^2$). **c.** Mean lymphocyte cell count relative to nearest PRAD or NEPC boundary. **d.** Data as in panel c but normalized to the binned tumor area. **e.** Percentage of TCF1 negative (neg), intermediate (int), or high (hi) CD8 T cells within *n*=5 independent RPM tumors. Data points represent the mean number of indicated CD8 T cells across TCF1 expression groups and error bars denote standard deviation. **f.** (Top) Representative immunofluorescence stains from RPM 10-week tumors of the indicated lymphocyte markers. (Bottom) Segmented FoV where each dot represents a CD8 T cell coordinates within a 10-week RPM tumor section. Each dot is color coded based on predetermined thresholds for TCF1 expression (FAU=fluorescence arbitrary units). Data representative of *n*=5 independent tumors and is related to Extended Data Fig. 5e. Dotted line in panels c-d represents the boundary of the histotype to a different histotype or the edge of a tumor. Positive values indicate cells found outside the histotype boundary; negative values indicate cells found inside the histotype boundary. Data in panels c-d derived from *n*=3 biologically independent samples. Error bars denote mean and standard error of the mean. Smoothened data curve fit by Loess method.

### EXTENDED DATA FIGURE 6:

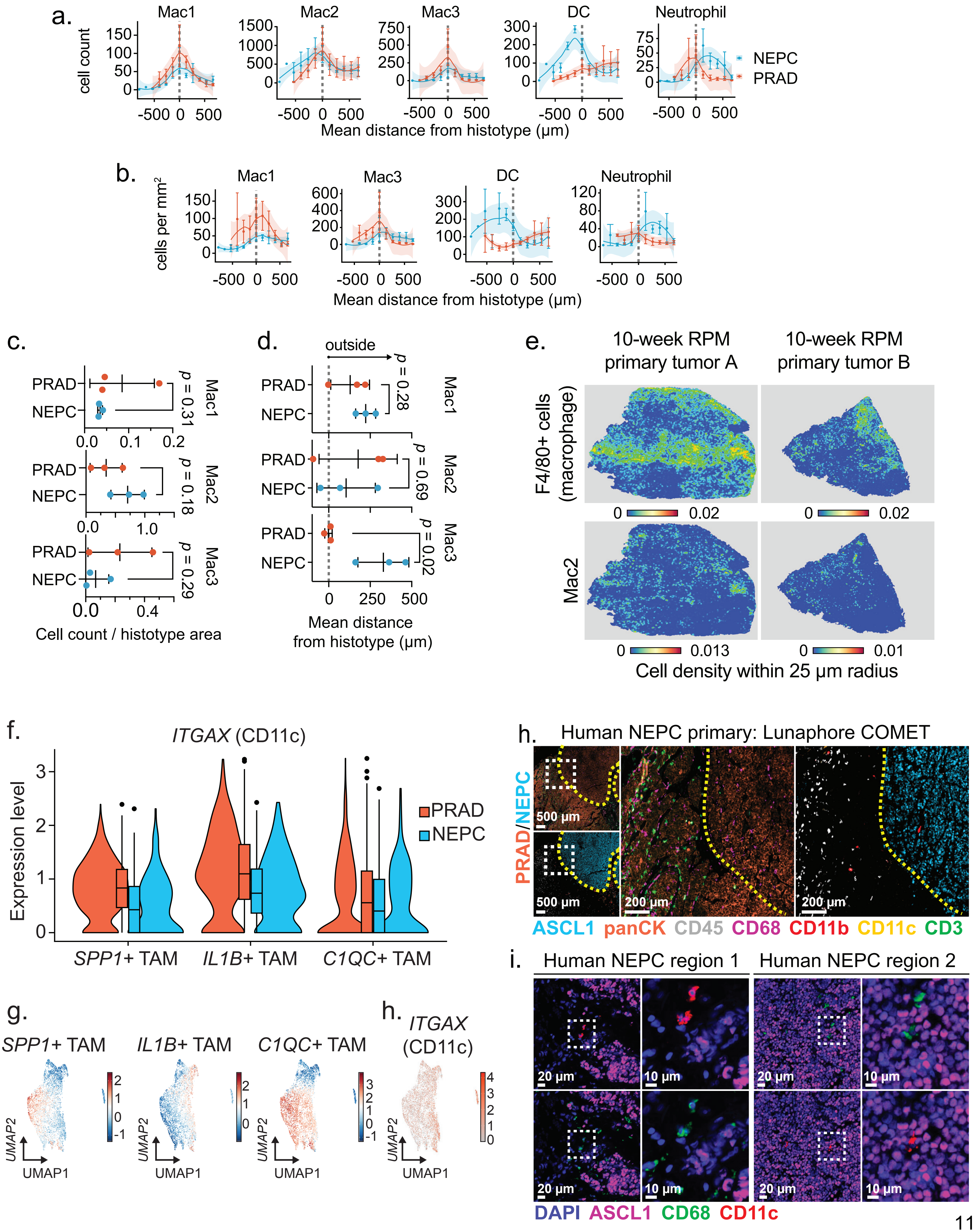

**Extended Data Figure 6:**

**a.** Frequency distribution of each indicated cell type within each binned distance outside or inside the defined interface region (NEPC or PRAD). Error bar denotes mean and standard error. **b.** Frequency distribution of each indicated cell type density within each binned distance from defined interface region (NEPC or PRAD). Error bar denotes mean and standard error. **c.** Dot plot depicting mean cell density away from the boundary of NEPC or PRAD tumor regions for the indicated macrophage subtypes. Error bar denotes mean and standard deviation. **d.** Dot plot depicting mean cell distance (μm) away from the boundary of NEPC or PRAD tumor regions for the indicated macrophage subtypes. Error bar denotes mean and standard deviation. **e.** Spatial cell type density (cell density within 25-μm radius) for the indicated myeloid cell types across two independent 10-week RPM tumors. Heatmap represents average cell density (cell number/$\mu m^2$). **f.** *ITGAX* (CD11c) expression across previously established tumor-associated macrophage populations identified within human PRAD and NEPC samples sequenced by scRNAseq. Violin plot depicts the median and first and third quartiles of *ITGAX* expression. **g.** Gene expression modules of TAM subsets identified within human PRAD and NEPC samples displayed in UMAP space, related to Extended Fig. 6f. Scale bar represents module score. **h.** *ITGAX* (CD11c) expression across all myeloid cell types identified within human PRAD and NEPC samples displayed in UMAP space, related to Extended Fig. 6f-g. Scale bar represents raw expression counts. **i.** Representative multiplexed IF of a human prostatectomy verified to contain mixed PRAD and NEPC pathology. Left zoomed out panels contain white dotted line indicating zoomed in regions on the right. Dotted yellow line represents the boundary of a panCK+ PRAD and ASCL1+ NEPC. **j.** Representative multiplexed IF of two distinct regions within the human NEPC sample shown in Extended Data Fig. 6i. Dotted square indicates magnified inset shown adjacent to lower magnification view. Dotted line in panels a-d represents the boundary of the histotype to a different histotype or the edge of a tumor. Positive values indicate cells found outside the histotype boundary; negative values indicate cells found inside the histotype boundary. Data in panels a-d derived from *n*=3 biologically independent samples. Smoothened data curve fit by Loess method.

### EXTENDED DATA FIGURE 7:

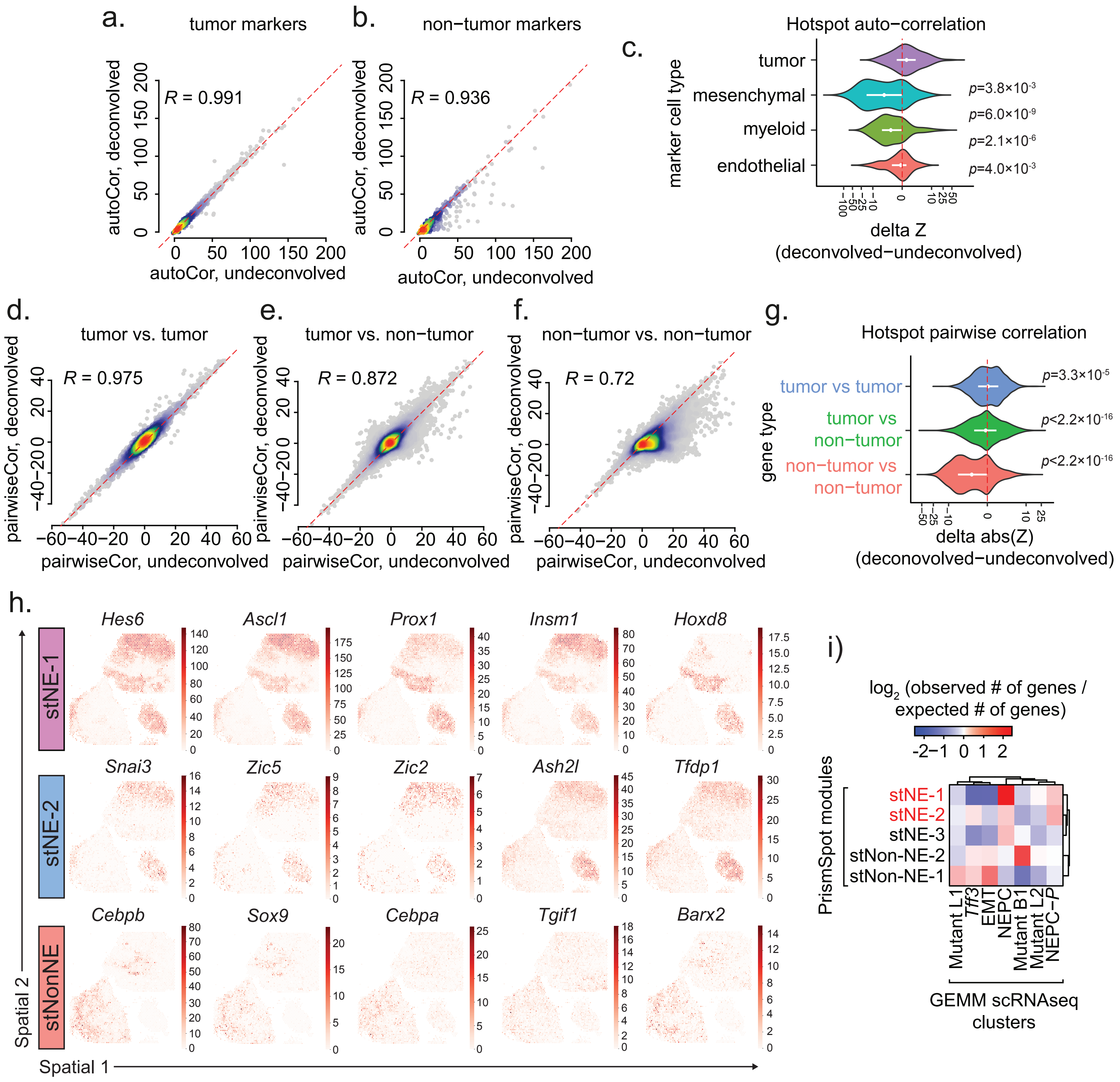

**Extended Data Figure 7:**

**a.** Density plot shows the Hotspot autocorrelation *Z*-scores for BayesPrism deconvolved (y-axis) vs undeconvolved expression (x-axis) across markers of tumor cells derived from GEMM scRNA-seq. Each dot in the density plot represents a tumor marker gene. Red dashed line marks y=x. **b.** Same as panel a. but across non-tumor cell types derived from GEMM scRNA-seq, including endothelial, myeloid, and mesenchymal cells. Autocorrelation computed using the deconvolved expression shows smaller value than that from the undeconvolved expression. **c.** Violin plot shows the distribution of the difference between *Z*-scores of PrismSpot and un-deconvolved Hotspot for marker genes of each cell type. Red dashed line marks zero. **d.** Density plot shows the Hotspot pairwise local correlation *Z*-scores for BayesPrism deconvolved (y-axis) vs undeconvolved expression (x-axis) between pairs of tumor marker genes defined using GEMM scRNA-seq. Each dot in the density plot represents a pair of tumor marker genes. Red dashed line marks y=x. **e.** Same as panel d. but between a tumor marker gene and marker gene from any non-tumor cell types. Using deconvolved expressions shrinks local correlation *Z*-scores towards zero. **f.** Same as panel e. but between a pair of marker genes from any non-tumor cell types. Using deconvolved expressions shrinks local correlation *Z*-scores towards zero. **g.** Violin plot shows the distribution of the difference between the absolute value of *Z*-scores of PrismSpot and un-deconvolved Hotspot for genes of each category. Red dashed line marks zero. **h.** Spatial expression (Visium) across the indicated tumor spatial modules identified by PrismSpot (related to Fig. 5e-f). Scale bar represents raw expression counts. **i.** Heatmap of the observed gene overlap normalized to expected gene overlap between PrismSpot modules and several tumor clusters derived by previously published GEMM models of NEPC.

EXTENDED DATA FIGURE 8:

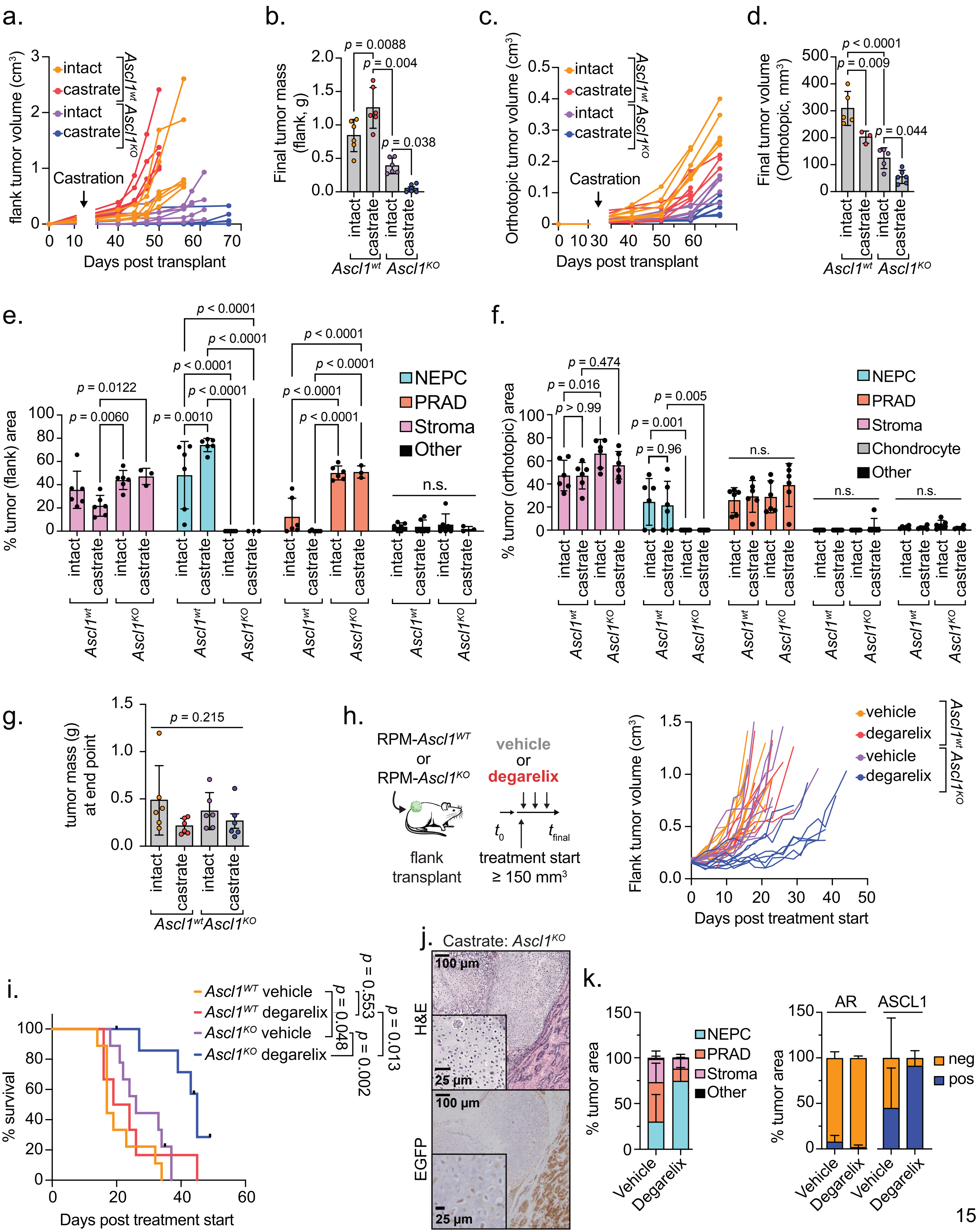

**Extended Data Figure 8:**

**a.** Individual longitudinal SQ tumor volumes as determined by caliper. Castration or sham surgery was performed 14 days post organoid transplantation. Data related to Fig. 6b. *n*=6 independent tumors across each group. Related to Fig. 6b. **b.** Final SQ tumor mass at experimental endpoint. *n*=6 independent tumors across each group. Statistics derived by one-way ANOVA with Sidak's multiple comparisons correction. Error bars denote mean and standard deviation. **c.** Individual longitudinal OT tumor volumes determined by ultrasound. *n*=6 independent tumors per group. Castration or sham surgery was performed 14 days post organoid transplantation. **d.** Final OT prostate tumor volumes determined by ultrasound. $Ascl1^{wt}$ intact *n*=5; $Ascl1^{wt}$ castrate *n*=3; $Ascl1^{KO}$ intact *n*=5; $Ascl1^{KO}$ castrate *n*=6. Statistics derived by one-way ANOVA with Sidak's multiple comparisons correction. Error bars denote mean and standard deviation. **e.** Bar charts representing percentage of SQ tumor area composed of the histological categories depicted in the figure legend. Data related to Fig. 6e. Each dot represents the average area per mouse. Statistics derived by two-way ANOVA with Tukey's multiple comparisons correction. Error bar denotes mean and standard deviation. *n*=6 independent tumors per group. **f.** Bar charts representing percentage of OT tumor area composed of the histological categories depicted in the figure legend. Data related to Fig. 6f. Each dot represents the average area per mouse. Statistics derived by two-way ANOVA with Tukey's multiple comparisons correction. Error bar denotes mean and standard deviation. *n*=6 independent tumors per group. **g.** Final OT tumor mass at experimental end point. *n*=6 per group. Statistics derived by one-way ANOVA. Error bars denote mean and standard deviation. Data related to Extended Data Fig. 8c; however, after cessation of ultrasound measurements, tumors were isolated and weighed once palpable or distress was observed. **h.** (Left) schematic of SQ transplantation assay by which mice are randomized into vehicle or degarelix treatment arms after tumors have established as determined by caliper. RPM-$Ascl1^{WT}$ vehicle and degarelix treated *n*=8, RPM-$Ascl1^{KO}$ vehicle *n*=9, RPM-$Ascl1^{KO}$ degarelix *n*=8 independent tumors per group. **i.** Survival of vehicle or degarelix treated mice with SQ transplants of the indicated RPM organoid genotypes. Statistics derived from the Log-rank (Mantel-Cox) test for each pair-wise comparison. Data related to Fig. 6c and Extended Data Fig. 8h. RPM-$Ascl1^{WT}$ vehicle *n*=9, RPM-$Ascl1^{WT}$ degarelix *n*=6, RPM-$Ascl1^{KO}$ vehicle *n*=9, and RPM-$Ascl1^{KO}$ degarelix *n*=8 independent mice. **j.** Representative (left) hematoxylin and eosin stain and (right) immunohistochemistry of EGFP of a single RPM-$Ascl1^{KO}$ tumor growth in a castrated host demonstrating chondrosarcomatoid histopathology. Scale bars denoted within the figure. **k.** (Left) Stacked bar charts representing percentage of SQ tumor area composed of the histological categories depicted in the figure legend. Data are quantified histology of RPM tumors derived from tumor bearing mice treated with vehicle (*n*=5) or degarelix (*n*=4) for 4 weeks after tumor establishment (≥150 $mm^3$) and represent average tumor area. (Right) Stacked bar charts of the percentage of AR- and ASCL1-pos (positive) or neg (negative) tumor cells (defined as EGFP+; CD45-; VIMENTIN-) within vehicle or degarelix treated RPM SQ tumors. Vehicle *n*=5, Degarelix *n*=4.

### EXTENDED DATA FIGURE 9:

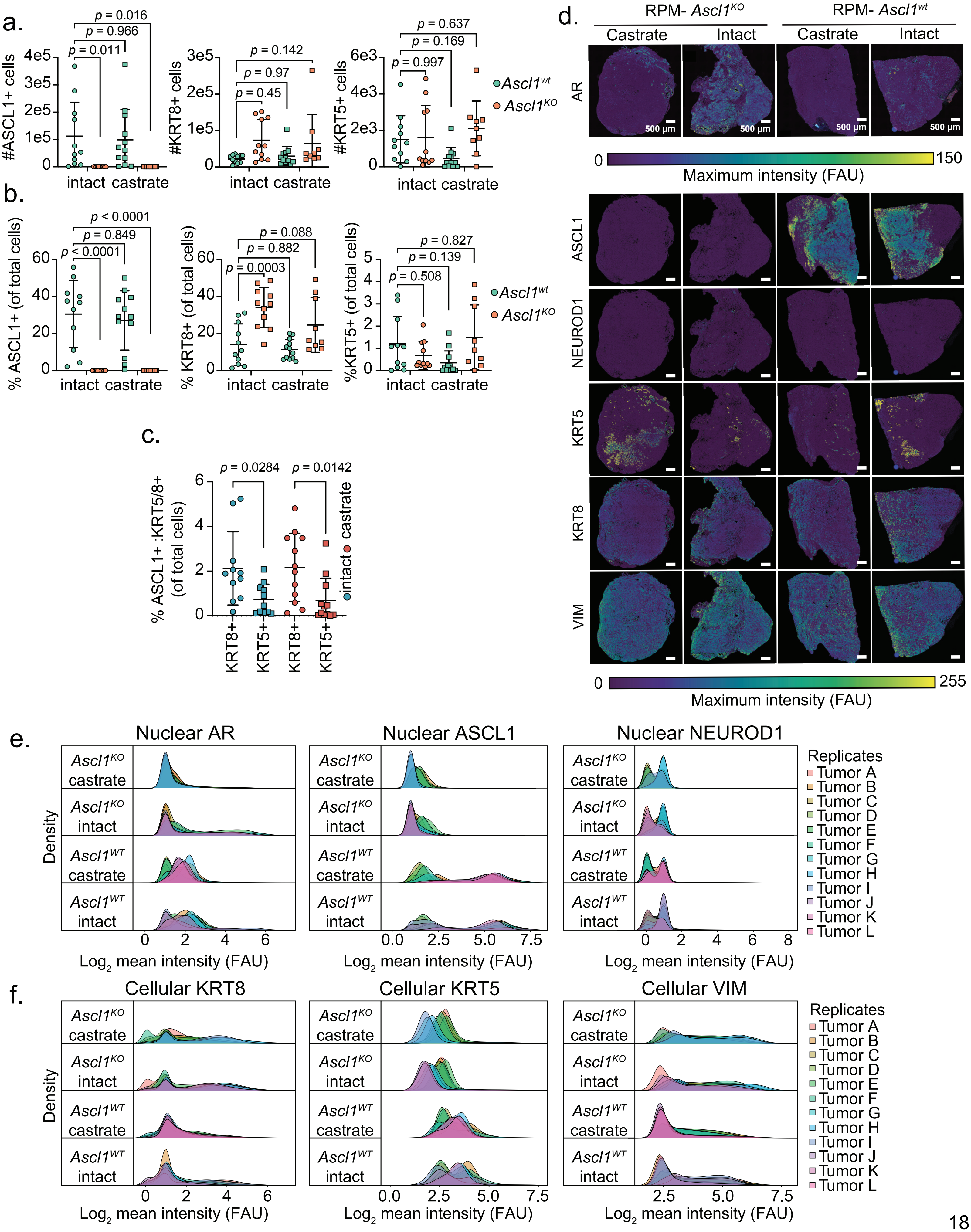

**Extended Data Figure 9:**

**a.** Number of ASCL1+ nuclei (left), KRT8+ cells (middle) and KRT5+ cells (right) in the indicated genotypes (legend on right hand-side) and treatment groups. **b.** Percentage of ASCL1+ nuclei (left), KRT8+ cells (middle) and KRT5+ cells (right) relative to total cells (DAPI+ nuclei) in the indicated genotypes (legend on right hand-side) and treatment groups. **c.** Percentage of ASCL1+ cells (DAPI+ nuclei) co-expressing either KRT8 or KRT5 within RPM-$Ascl1^{WT}$ tumors in either intact or castrated hosts. Statistics in panels a-c derived from two-way ANOVA with Tukey's multiple comparisons correction. Error bars in panels a-c denote mean and standard deviation. **d.** Field of view images depicting maximum intensity score for all segmented cells within representative RPM tumors of the indicated genotype and treatment group. FAU=fluorescence arbitrary units. Data representative of n≥10 individual tumors. **e.** Density plots of the $\log_2(x+1)$ transformed mean fluorescence intensity for each nuclear protein. Each density plot represents signal intensity of tumor cells across an independent tumor. **f.** Density plots of the $\log_2(x+1)$ transformed mean fluorescence intensity for each cytoplasmic protein. Each density plot represents signal intensity of tumor cells across an independent tumor.

EXTENDED DATA FIGURE 10:

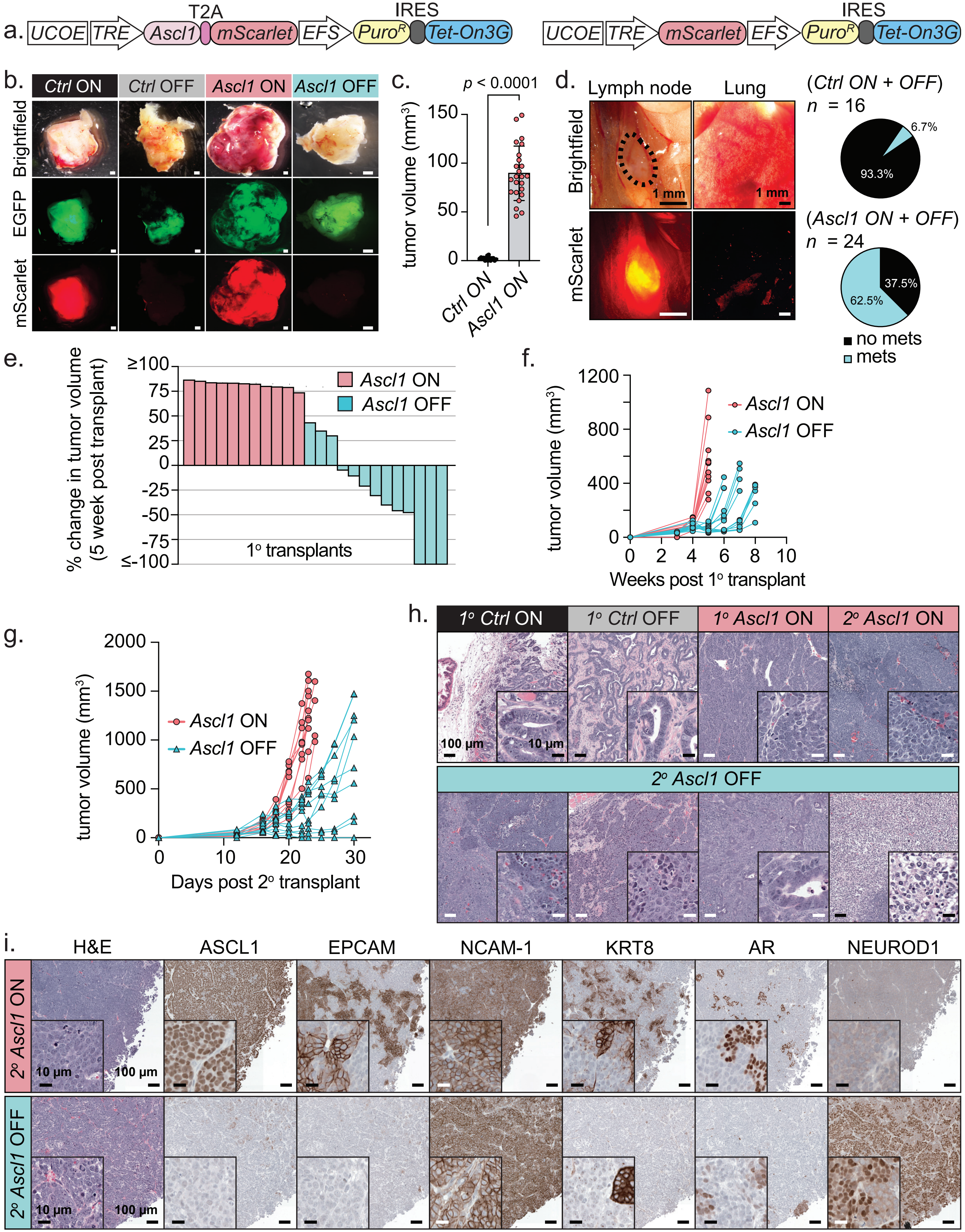

**Extended Data Figure 10:**

**a.** Schematic of the dox-inducible lentiviral vector used within this conditionally overexpress *Ascl1-P2A-mScarlet* or (not shown) *mScarlet* in RPM-$Ascl1^{KO}$ organoids. Related to Fig. 8. **b.** Representative stereoscopic images (brightfield and fluorescent) of OT tumors isolated from the indicated dox maintained or withdrawn conditions. Scale bar represents 1 mm. **c.** Tumor volumes determined by ultrasound 4 weeks post OT engraftment of the indicated groups. All mice were maintained on dox chow. Statistics derived from two-sided *t*-test and error bars denote mean and standard deviation. *Ctrl* ON *n*=15, *Ascl1* ON *n*=13. **d.** (Left) Representative stereoscopic images (brightfield and fluorescence) of the draining lymph nodes and lungs of mice bearing OT *Ascl1* ON tumors. (Right) Pie charts indicating frequency of regional or distal micro-metastatic dissemination. **e.** Percent change in primary recipient (1$^{o}$) OT tumor volume between 4-5 weeks post OT transplantation of *Ascl1* ON organoids, as determined by ultrasound. *Ascl1* OFF cohort has been withdrawn from dox-chow for 1 week. **f.** Longitudinal primary OT tumor volumes determined by ultrasound for the indicated groups. *Ascl1* ON *n*=11, *Ascl1* OFF *n*=13). **g.** Longitudinal secondary recipient (2$^{o}$) SQ tumor volumes determined by caliper for the indicated groups. *Ascl1* ON and OFF *n*=10. **h.** Representative H&E images of the indicated groups spanning both primary recipient (OT) and secondary recipient (SQ) transplanted mice. **i.** Representative serially sectioned tumors stained for H&E and IHC of the indicated markers across *Ascl1* ON and *Ascl1* OFF secondary transplant recipient mice (SQ). Images displayed represent regions maintaining NEPC histology and high fraction of NCAM-1 marker expression. All scale bars depicted in the figure panels.
