## Supplementary Figures 1-11 for "The neuroendocrine transition in prostate cancer is dynamic and dependent on ASCL1"

SUPPLEMENTARY FIGURE 1:

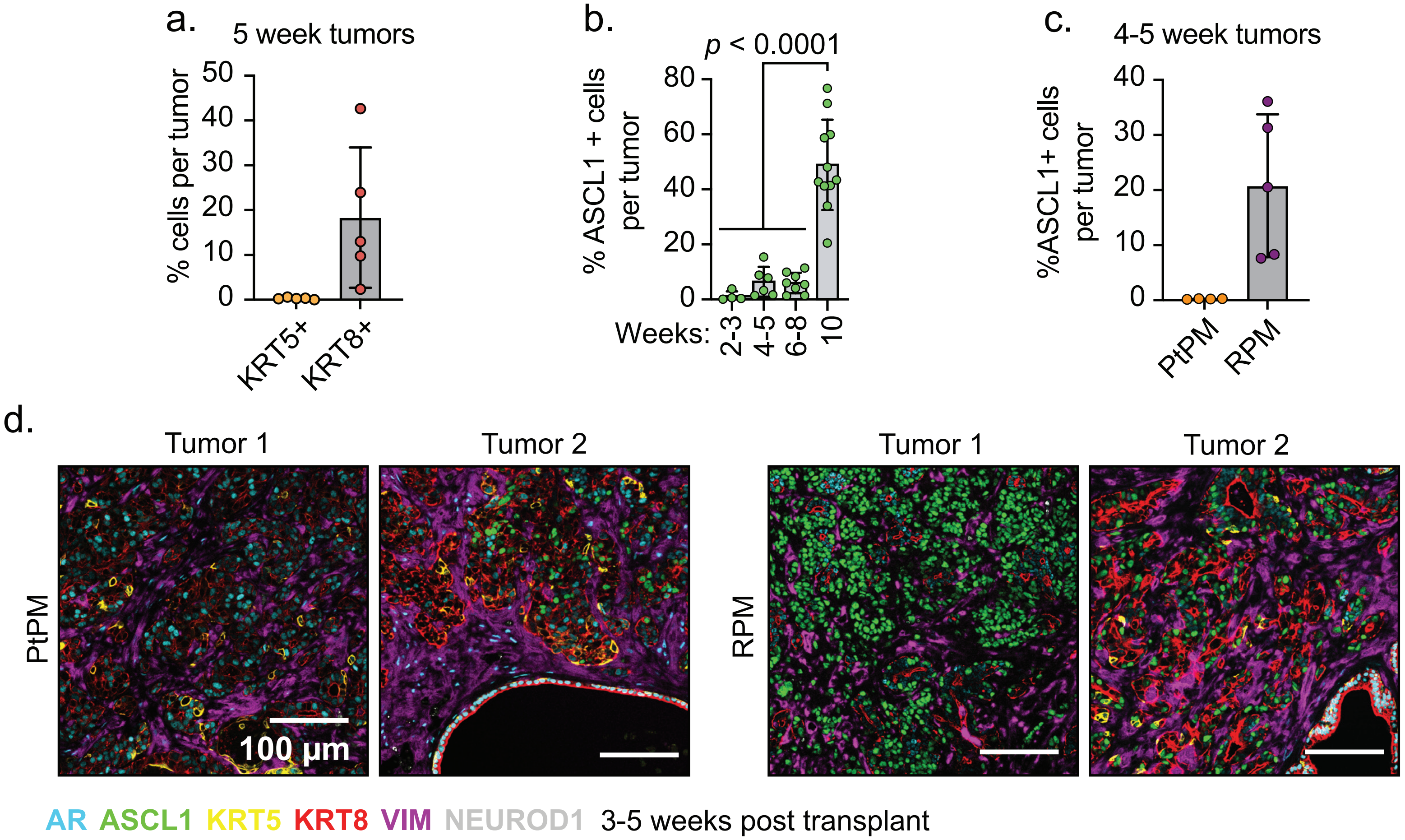


**Supplementary Figure 1:**

a. Average percentage of KRT8+ or KRT5+ tumor cells relative to total detected cells (DAPI+ nuclei) within individual RPM OT tumors 5 weeks post-transplantation. Each dot represents the average marker positive cell per mouse tumor. Error bars denote mean and standard deviation. Data related to Fig. 2a-b and represent the average positive cell number per tumor. Data derived from $n=5$ independent RPM OT tumors.

b. Average percentage of total ASCL1+ cells relative to total detected cells (DAPI+ nuclei) within individual RPM tumors at the indicated time points. Each dot represents the average marker positive cell per mouse tumor. 2-3 weeks, $n=4$; 4-5 weeks, $n=6$; 6-8 weeks, $n=8$; 10 weeks, $n=11$. Statistics derived using one-way ANOVA with Tukey's multiple comparisons correction. Error bars denote mean and standard deviation.

c. Percentage of cells staining positively for nuclear ASCL1 from tumors harvest 4 to 5 weeks post orthotopic transplantation. Data related to Fig. 2a-b and represent the average positive cell number per tumor. Data derived from $n=4$ (PtPM) and $n=5$ (RPM) independent OT tumors per group.

SUPPLEMENTARY FIGURE 2:

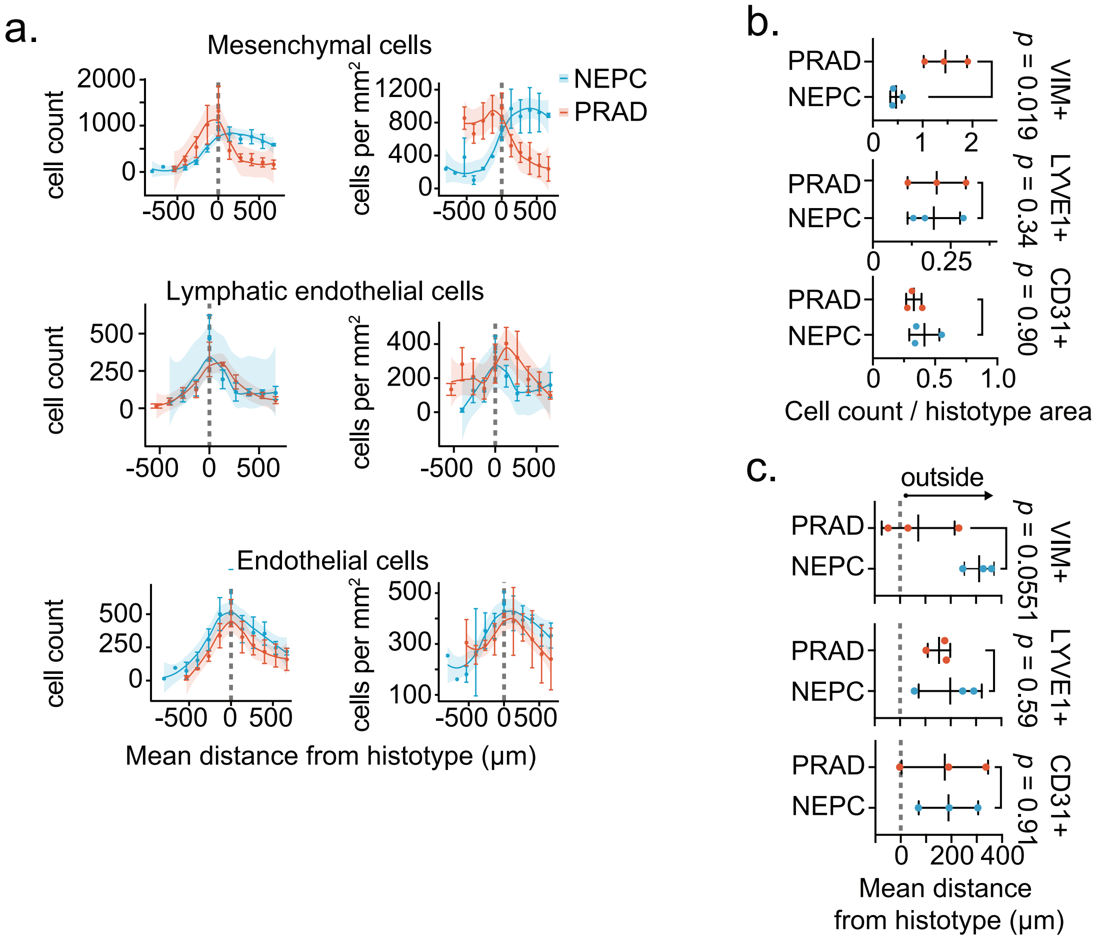


**Supplementary Figure 2:**

a. Cell number of each indicated non-immune stromal cell type within each binned distance outside or inside the defined interface region (NEPC or PRAD). Shaded region approximated through Loess method. Scale bar represents mean and standard error of the mean of the cell count per bin. $n=3$ unique regions collected from 3 independent tumors.

b. Frequency distribution of each indicated non-immune stromal cell type within each binned distance outside or inside the defined interface region (NEPC or PRAD). Shaded region approximated through Loess method. Scale bar represents mean and standard error of the mean of the cell count per bin. $n=3$ unique regions collected from 3 independent tumors.

c. Dot plot depicting mean cell density outside or inside NEPC or PRAD tumor regions for the indicated non-immune stromal cell types. VIM+, mesenchymal cells: LYVE1+, lymphatic endothelial cells; CD31+, endothelial cells. $n=3$ unique regions collected from 3 independent tumors. Dotted line in panels a-c indicate the boundary of a PRAD or NEPC tumor region.

### SUPPLEMENTARY FIGURE 3:

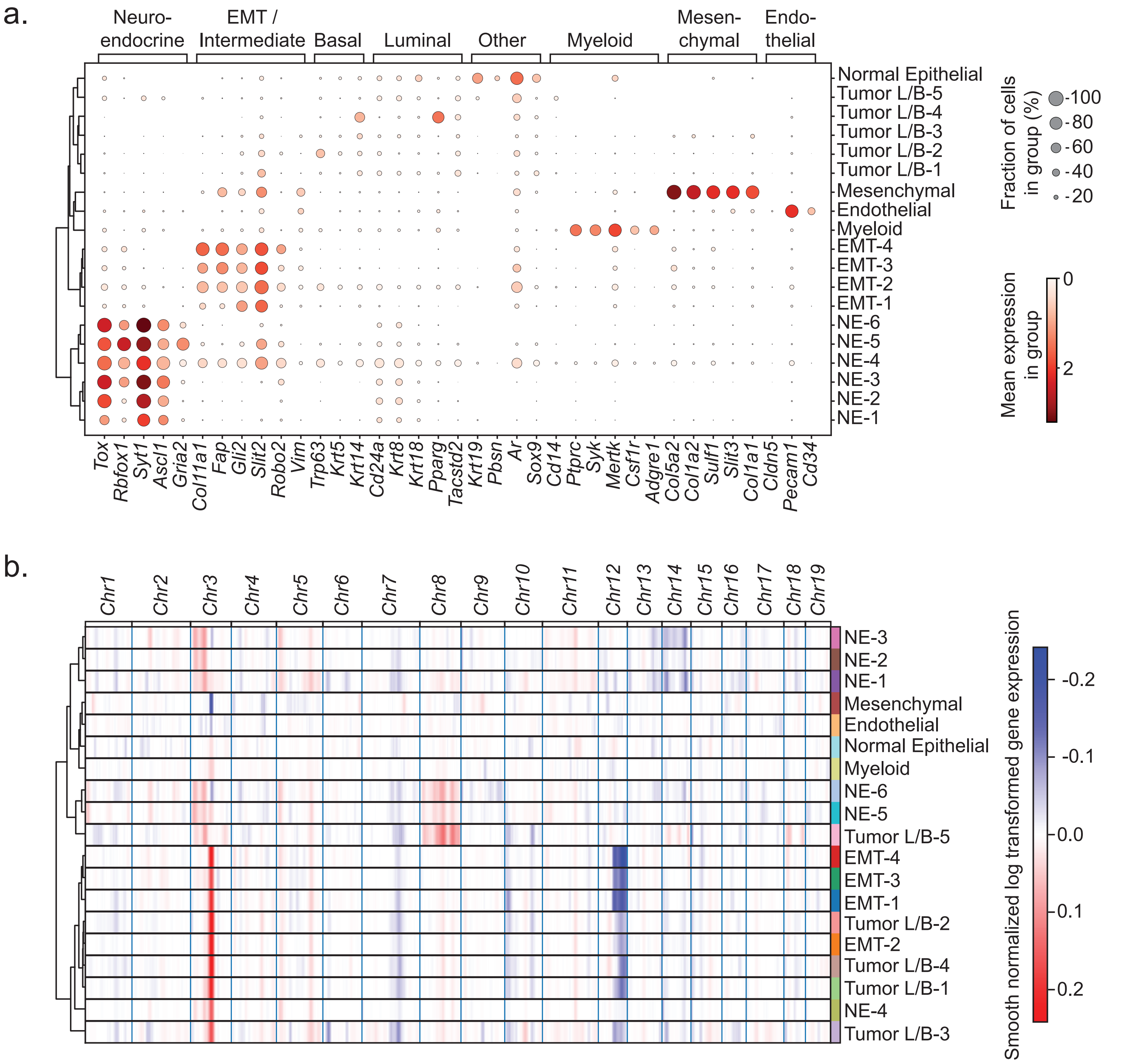


**Supplementary Figure 3:**

a. Dot plot of selected marker genes (DEGs) associated with each of the indicated cell types from snRNA-seq profiled from RPM tumors. Data represent the mean normalized $\log_2(X+1)$ expression scaled from 0 to 1. Dot size represents percent of cells expressing a given gene.

b. Inferred copy number variation (CNV) applying inferCNV to snRNAseq profiled from RPM tumors. Data represent the smooth normalized and log-transformed gene expression values across each murine chromosome. Inferred CNV for each cell type identified in panel a listed on the right.

SUPPLEMENTARY FIGURE 4:

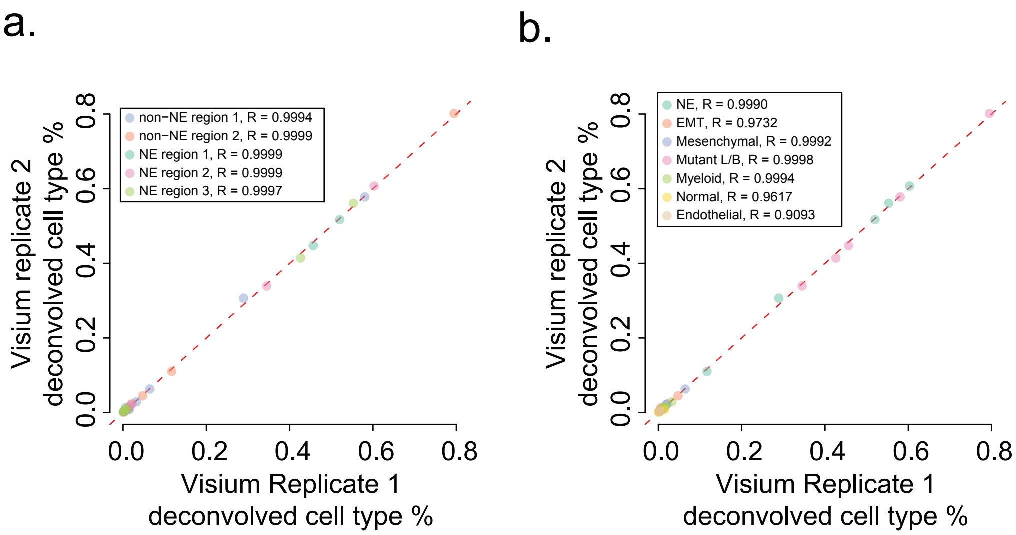


**Supplementary Figure 4:**

a. Scatter plot shows the mean cell type fraction inferred by BayesPrism across two technical replicates for each cell type in each region, colored by regions defined by histology. NE = neuroendocrine.

b. Same as panel a. but colored by cell type. NE = neuroendocrine, EMT = epithelial to mesenchymal transition tumor cells. Tumor L/B = tumor cells with markers associated with luminal and basal lineages.

### SUPPLEMENTARY FIGURE 5:

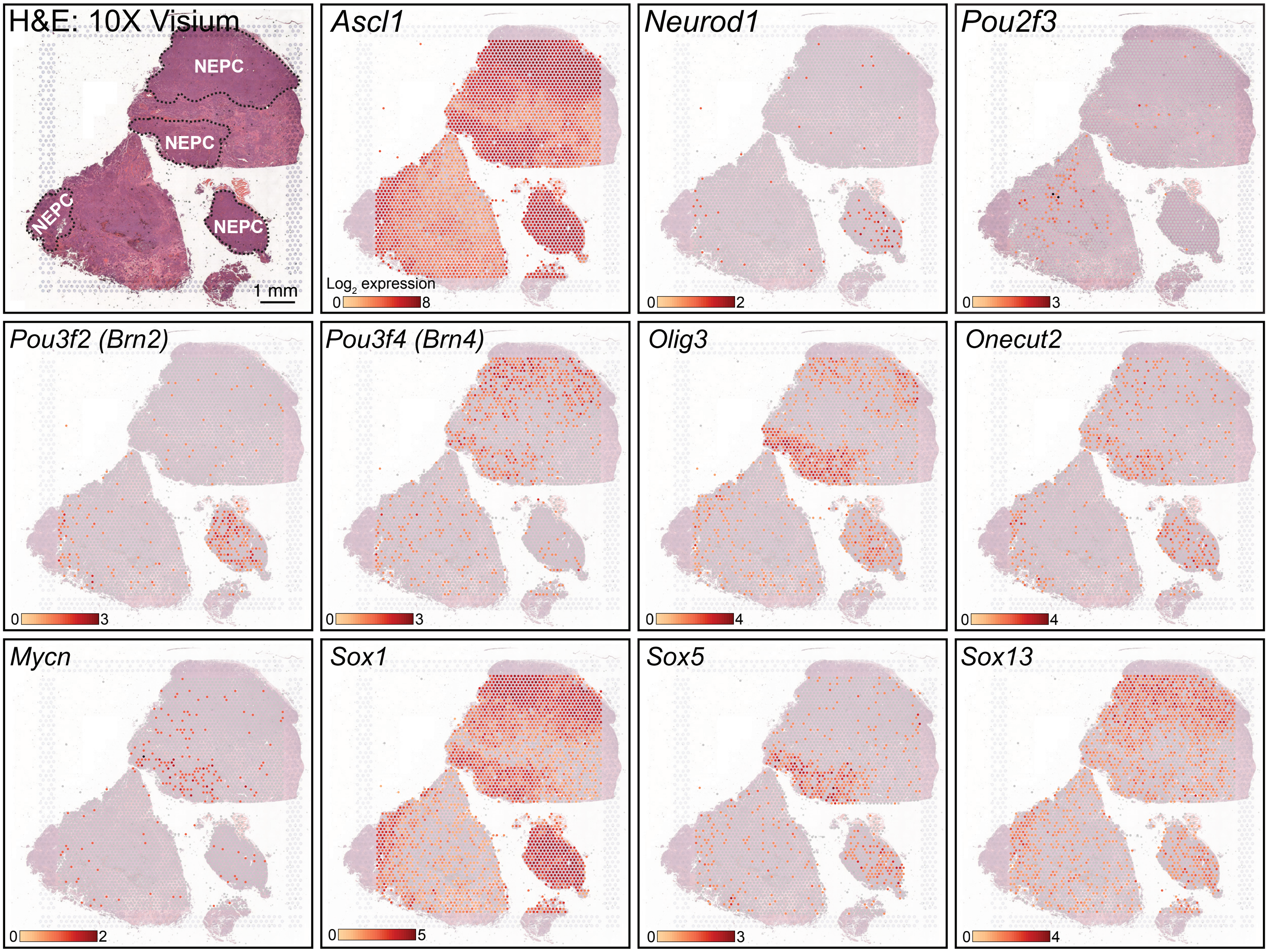


**Supplementary Figure 5:**

Log$_2$ spatial expression (Visium) across the indicated genes. Top Left: H&E depicting NEPC tumor regions within two independent 10-week RPM tumors (related to Fig. 5c).

SUPPLEMENTARY FIGURE 6:

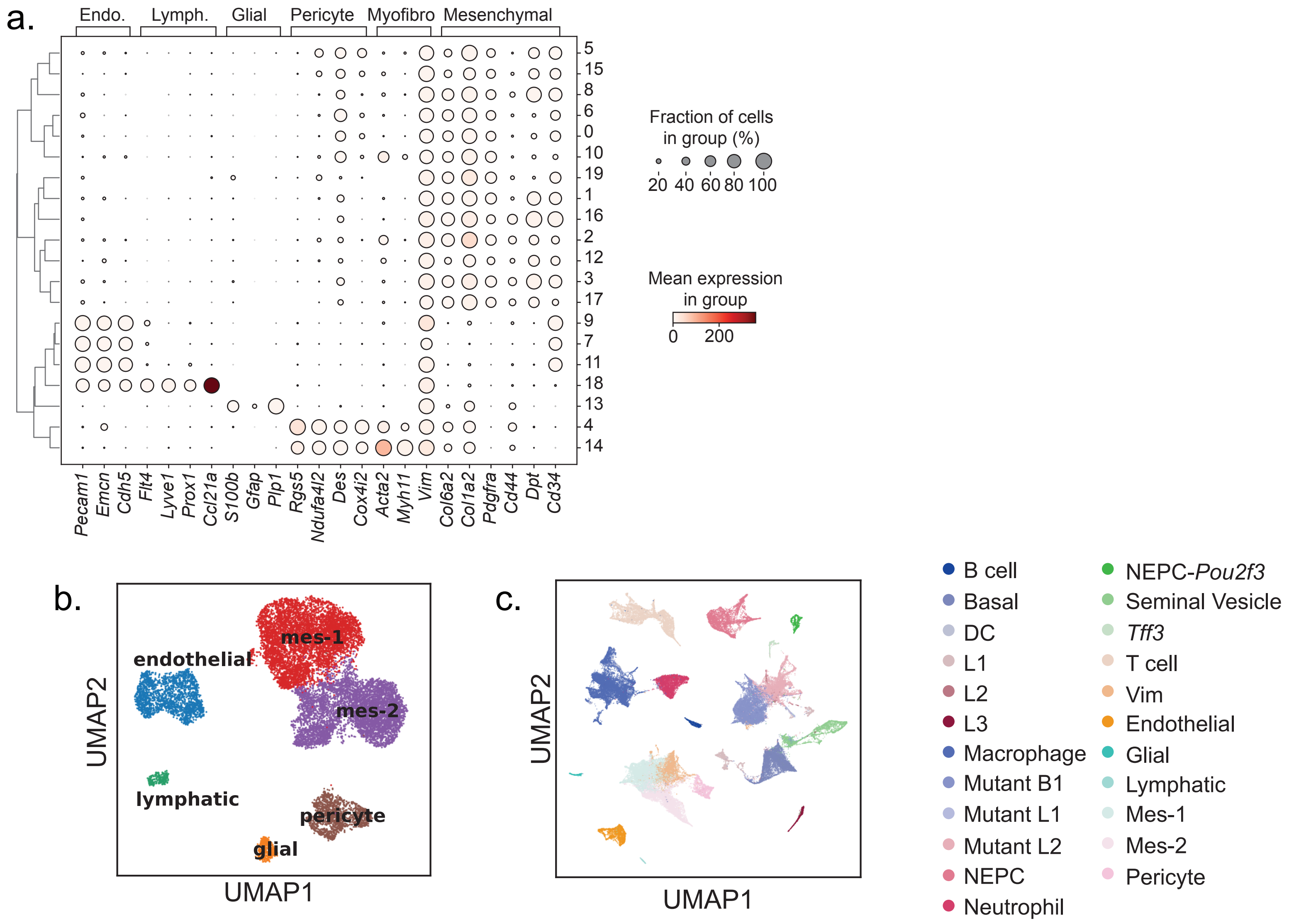


**Supplementary Figure 6:**

a. Dot plot of marker genes associated with each of the indicated cell types across phenograph clusters. Data represent the mean normalized $\log_2(X+1)$ expression scaled from 0 to 1. Dot size represents percent of cells expressing a given gene.

b. UMAP of non-tumor cell populations in GEMM scRNA-seq dataset, colored by cell types. Mes = Mes-enchymal.

c. UMAP of the original GEMM scRNA-seq dataset, colored by cell types with the GFP-negative stromal populations replaced with finer-grained cell types.

SUPPLEMENTARY FIGURE 7:

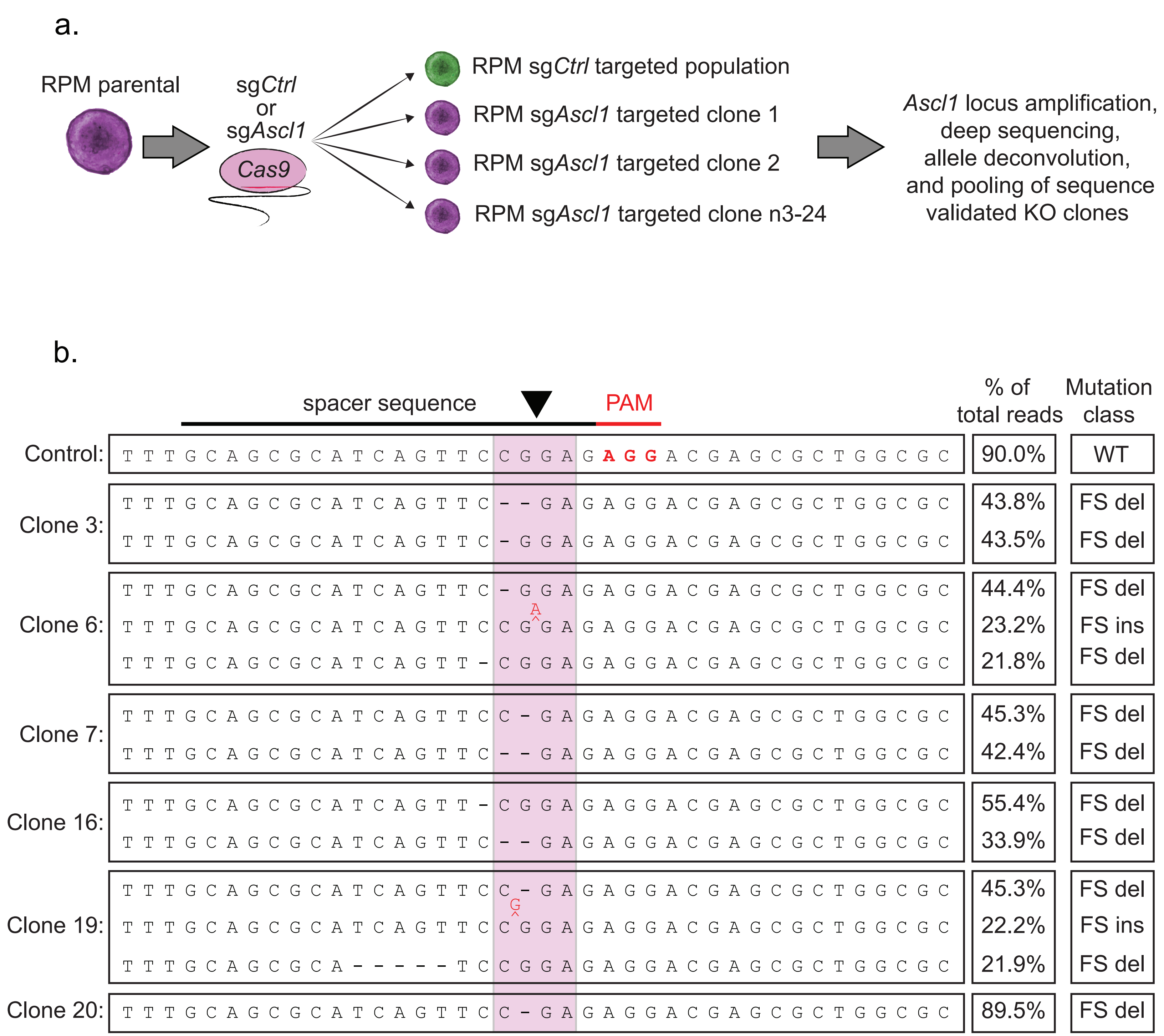


**Supplementary Figure 7:**

a. Schematic of the derivation and validation of RPM control and *Ascl1*$^{KO}$ organoids. A total of 24 organoid clones were selected and expanded after sg*Ascl1* Cas9 ribonucleoprotein electroporation for Illumina-based deep sequencing analysis of targeted *Ascl1* locus.

b. Representative alleles obtained from sequence validated RPM-*Ascl1*$^{KO}$ clones. Plot summarizes the mutational analyses of locus-specific deep sequencing showing the *Ascl1* wild-type locus containing the sg*Ascl1.1* binding site (black, spacer sequence) and protospacer adjacent motif (PAM) sequence (red) along with representative mutant alleles. Dashes indicate deletion events and red arrows indicate insertion events. Percentage of total alleles and mutation class shown on the right. WT = wild type, FS del = frameshift deletion, FS ins = frameshift insertion. Related to Supplementary Table 9.

### SUPPLEMENTARY FIGURE 8:

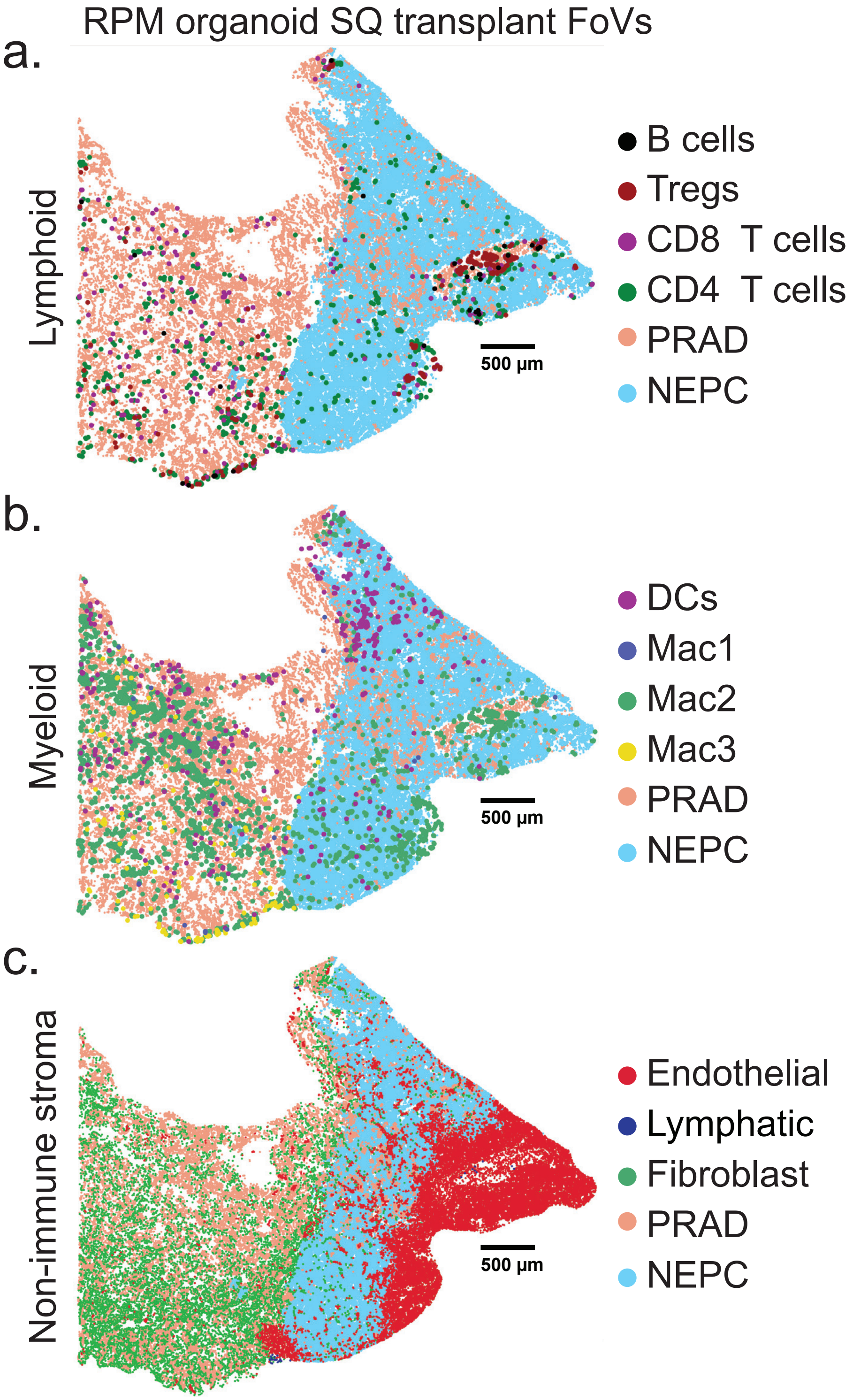


**Supplementary Figure 8:**

Representative segmented field of view (FoV) for the indicated general **a.** lymphoid cell, **b.** Myeloid, and **c.** non-immune stromal types in 8-week SQ RPM tumor. Data are representative of 5 independent SQ tumors. SQ = subcutaneous tumor.

### SUPPLEMENTARY FIGURE 9:

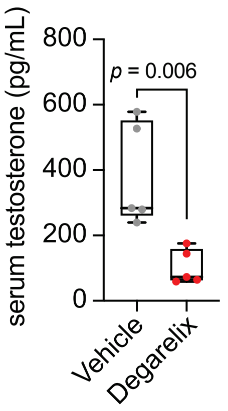


**Supplementary Figure 9:**

Serum testosterone concentrations assessed by ELISA collected 2 weeks post SQ single dose of degarelix (15mg/kg) or vehicle (5% mannitol). Box and whisker plot denotes mean and top and bottom quartiles. Statistics derived using two-sided Student's *t*-test. SQ = subcutaneous tumor.

SUPPLEMENTARY FIGURE 10:

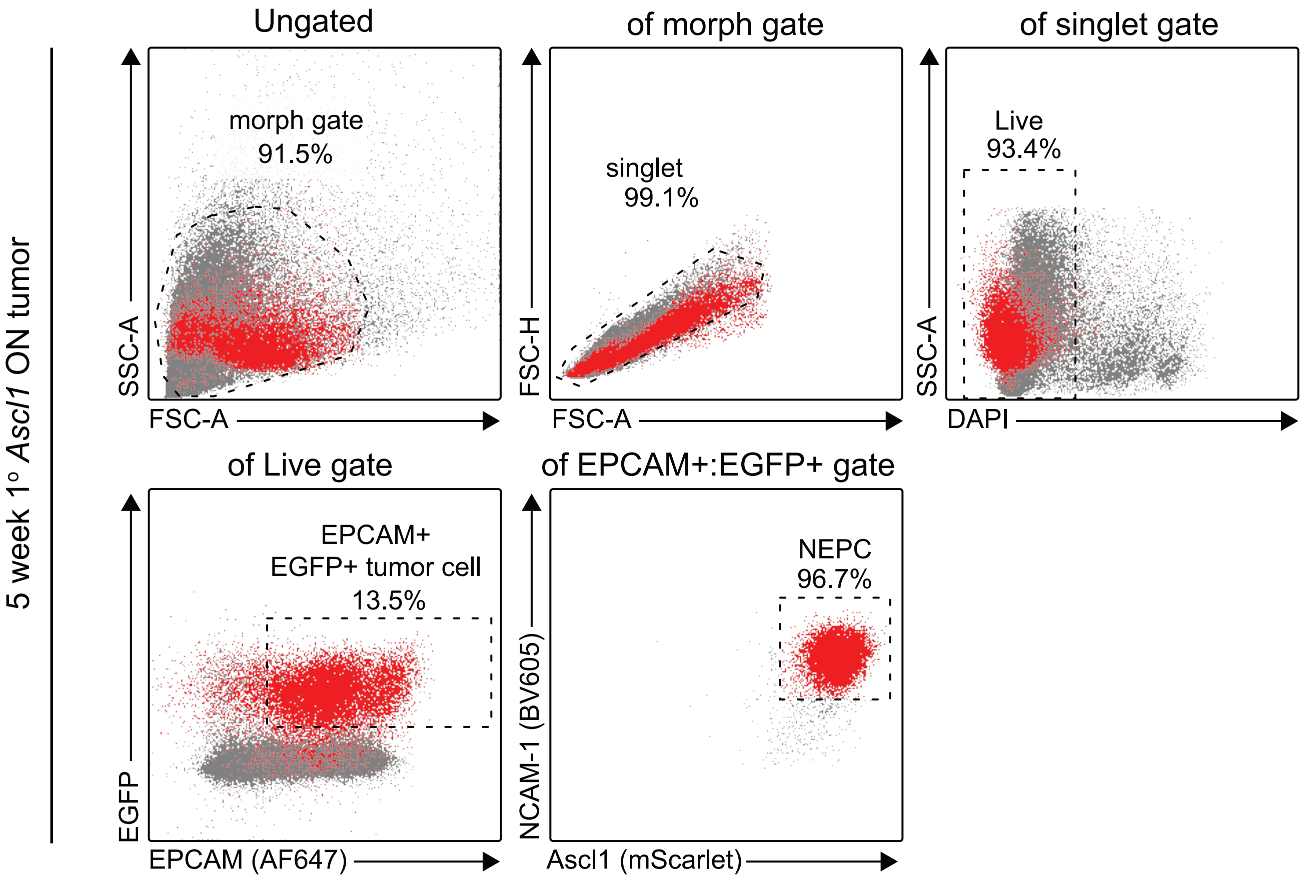


**Supplementary Figure 10:**

Gating strategy used to purify RPM-NEPC tumor fraction from primary (1°) *Ascl1 ON* mice on dox for 5 weeks post transplantation. Data related to Fig. 8c. See methods for more information.

SUPPLEMENTARY FIGURE 11:

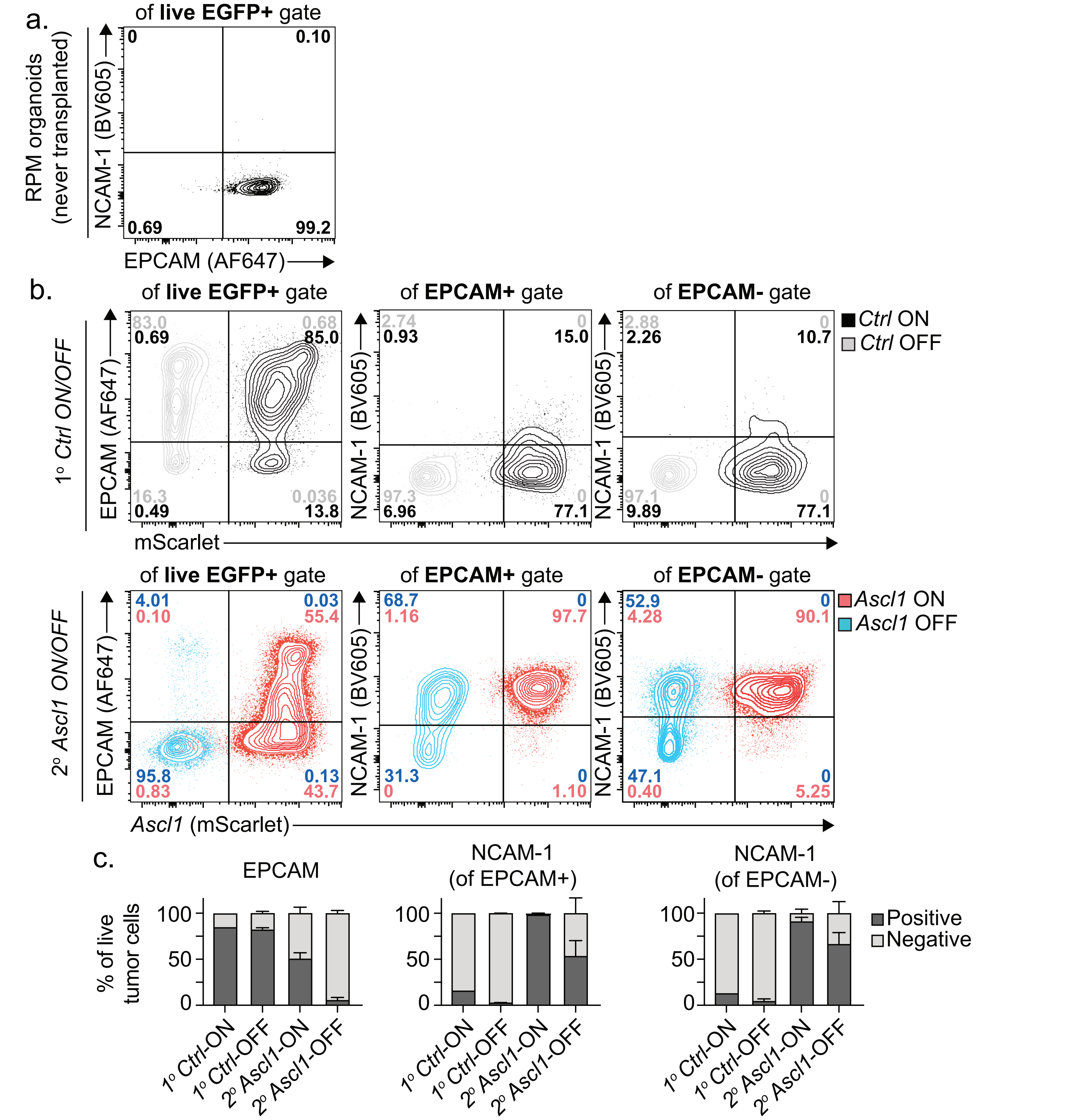


**Supplementary Figure 11:**

a. Representative flow plot of control (DAPI-negative) RPM organoids (never transplanted, EGFP+) stained with NCAM-1 or EPCAM and used for positive and negative gating in panels b-c.

b. (Top) Representative OT *Ctrl ON* or *Ctrl OFF* primary (1°) tumors (endogenous *Ascl1*$^{KO}$, can only initiate and progress as PRAD) stained for the depicted markers. Used as negative controls for NCAM-1 staining. (Bottom) Representative SQ *Ascl1 ON* or *Ascl1 OFF* secondary (2°) tumors stained for the depicted markers. Data representative of $n$=5 tumors per group. Outliers marked as dots. Note 95.8% of *Ascl1 OFF* tumors (dox withdrawal for 2 weeks) have lost EPCAM expression.

c. Stacked bar charts for the indicated tumors staining positively or negatively for (left) EPCAM, (middle) NCAM-1 of EPCAM+ gate, or (right) NCAM-1 of EPCAM- gate. Related to panel b. Error bars represents mean and standard deviation and are derived from $n$=5 independent tumors per group.
