## Supplementary figures and images for "The neuroendocrine transition in prostate cancer is dynamic and dependent on ASCL1"

### Source Data File: Unedited western blots

Source Data Fig 1:

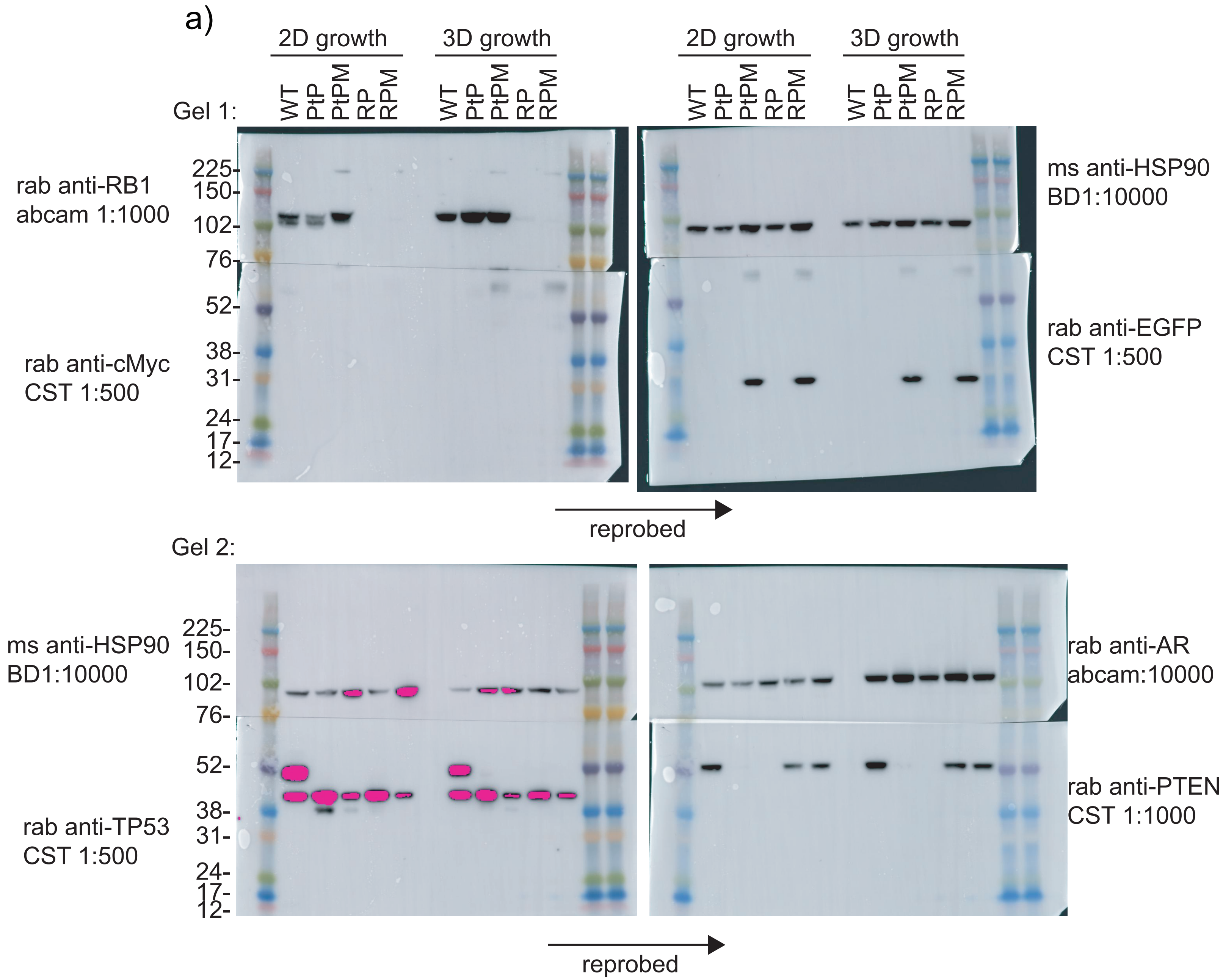
